# Resting state changes in aging and Parkinson’s disease are shaped by underlying neurotransmission – a normative modeling study

**DOI:** 10.1101/2023.10.18.562677

**Authors:** Jan Kasper, Svenja Caspers, Leon D. Lotter, Felix Hoffstaedter, Simon B. Eickhoff, Juergen Dukart

## Abstract

Human healthy and pathological aging is linked to a steady decline in brain resting state activity and connectivity measures. The neurophysiological mechanisms underlying these changes remain poorly understood. Making use of recent developments in normative modeling and availability of in vivo maps for various neurochemical systems, we test in the UK Biobank cohort (N=25,917) if and how age- and Parkinson’s disease related resting state changes in commonly applied local and global activity and connectivity measures co-localize with underlying neurotransmitter systems. We find the distributions of several major neurotransmitter systems including serotonergic, dopaminergic, noradrenergic and glutamatergic neurotransmission to explain age-related changes as observed across functional activity and connectivity measures. Co-localization patterns in Parkinson’s disease deviate from normative aging trajectories for these, as well as for cholinergic and GABAergic neurotransmission. The deviation from normal co-localization of brain function and GABAa correlates with disease duration. These findings provide new insights into molecular mechanisms underlying age- and Parkinson’s related brain functional changes. Combining normative modeling and neurotransmitter mapping may aid future research and drug development through deeper understanding of neurophysiological mechanisms underlying specific clinical conditions.

## Introduction

Understanding the neurophysiological mechanisms underlying healthy and pathological brain aging is an essential component for development of successful prevention, detection, and intervention strategies for age-related diseases and cognitive decline. Despite ample evidence for age-related decline in various brain functional measures there is only limited understanding of the neurophysiological mechanisms underlying these changes. As the interplay of different neurotransmitter systems is the major contributor to the blood oxygen level dependent (BOLD) signal it is plausible to assume that changes in respective systems also manifest in age-related functional alterations as observed using resting state functional magnetic resonance imaging (rs-fMRI).

Most commonly applied rs-fMRI measures estimate either the temporal change in the regional amplitude of the BOLD signal as a measure of local activity, or compute correlations of BOLD time series across different brain regions as a measure of synchronicity. Previous rs-fMRI studies reported aging related reductions in local brain activity primarily in medial and frontal regions^1–4^. These alterations are complemented by reduced local synchronicity in cortical, sub- cortical, and cerebellar motor structures^5^ and increased local synchronicity mainly in hippocampal and thalamic regions^4,6^. In parallel, positron emission tomography (PET) studies of aging reported reduced serotonergic^7–11^, dopaminergic^12–15^, glutamatergic^16–18^, cholinergic^19^ and norepinephrinergic^20^ neurotransmission whilst evidence for increased availability was found for GABAa^21^ and µ-opioid^22^ receptors. Whilst both modalities point to complex functional re-organization during aging, the relationship between the respective rs-fMRI and PET findings remains poorly understood.

Testing for associations between the regional availability of specific neurotransmitter receptors or transporters as derived from positron emission tomography (PET) and rs-fMRI derived functional signal has been shown to be a promising way to estimate their impact on the observed brain phenotypes^23,24^. To date, age-related changes in these associations have not been systematically addressed. Such age-related changes in co-localization patterns may reflect the altered availability or functional decline of cell populations that exhibit the respective neurochemical properties. Following this logic, disease related changes in such associations would support the notion of respective neurotransmitter systems being particularly affected by the respective neuropathology^25^.

Understanding of typical age-related co-localization changes of brain function and neurotransmitter systems can inform the study of pathological deviations from such as for example observed in Parkinson’s disease^25^ (PD). Previous studies provided evidence for increased vulnerability of specific neurotransmitter systems in PD including dopaminergic^26–31^, serotonergic^32–34^, glutamatergic^35,36^, GABAergic^30^, histaminergic^37^, cholinergic^38–40^, and norepinephrinergic^41,42^ neurotransmission. In our previous work, deviations from normal brain function in PD were related to the availability of D2 and 5-HT1b receptors, supporting the notion of specific vulnerability of these neurotransmitter systems^25^. However, if and how far these PD-related alterations deviate from typical age-related co-localization changes remains to be shown.

To address these questions, we adopt a normative modeling approach to test for aging-effects on co-localizations between brain resting state functional measures and underlying neurotransmission in a large cohort drawn from the UK Biobank (n = 25,917). We then replicate and extend the previous evidence of increased vulnerability of specific neurotransmitter systems in PD (n = 58). We test for co-localization of PET-derived distributions for all major neurotransmitter systems with commonly deployed rs-fMRI derived activity and connectivity measures.

## Methods

### Cohorts

We included 25,917 adult subjects from the UK Biobank who were not diagnosed with any of the listed diseases (Supplementary Table 1) as a control cohort for normative modeling of aging effects. Subjects with psychiatric, cognitive, and neurological disorders with known effect on brain structure (like demyelinating diseases or atrophies) and function (like intellectual disabilities, psychoactive substance use, or schizophrenia) were excluded. We additionally identified a group of 58 subjects from the UK Biobank who were diagnosed with idiopathic Parkinson’s disease (ICD-10, G20) before their imaging session. An overview of both groups is provided in Table 1.

**Table 1:**
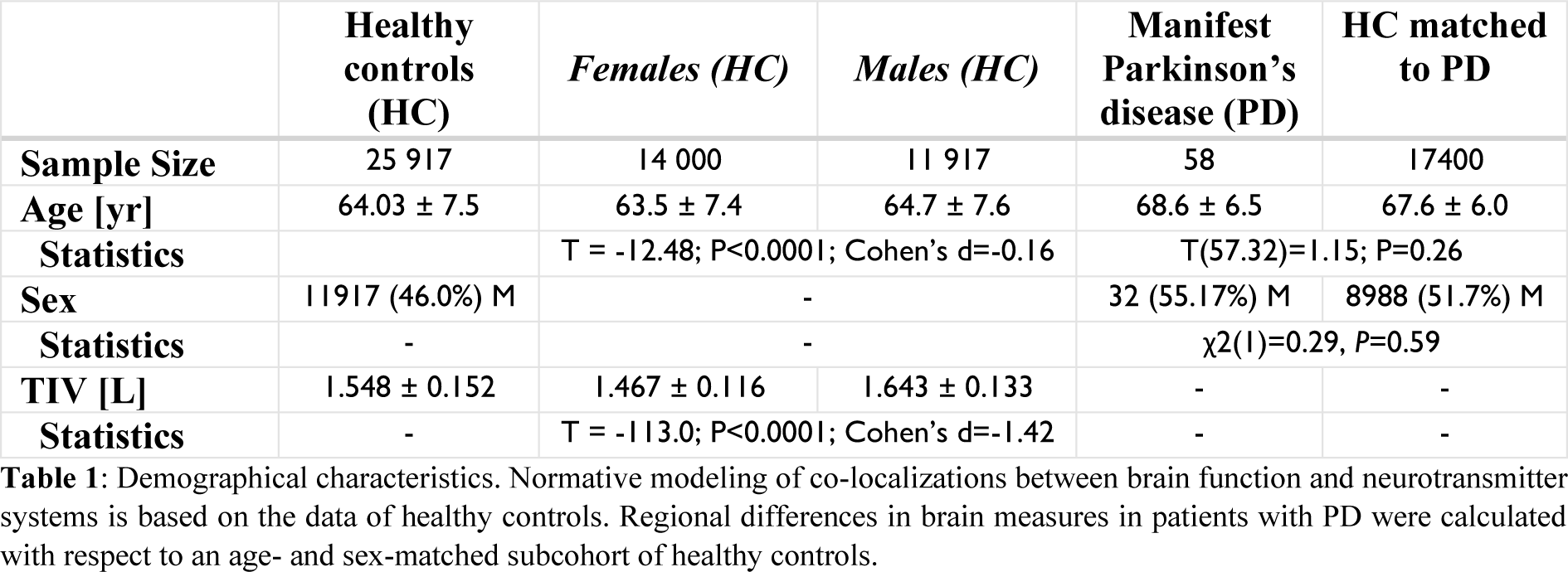
Demographical characteristics. Normative modeling of co-localizations between brain function and neurotransmitter systems is based on the data of healthy controls. Regional differences in brain measures in patients with PD were calculated with respect to an age- and sex-matched subcohort of healthy controls.

### Preprocessing of resting-state functional imaging data

We used resting-state functional MRI data provided and processed by the UK Biobank (referred to as “filtered_func_data_clean.nii”, cf. UK Biobank Brain Imaging Documentation for a detailed description^43^). The pipeline consisted of primary T1 quality control, gradient distortion correction, motion correction, grand-mean intensity normalization, high-pass temporal filtering, echo planar imaging unwarping, gradient distortion correction unwarping, and the removal of structural artefacts via ICA+FIX. Further processing was conducted with SPM12^44^, the FMIRB Software Library (FSL v5.0^45^), and the CONN toolbox^46^ implemented in MATLAB (v2020b). Functional images were transformed into MNI space using a general reference template provided by FSL and a subject specific warping image. After resampling (3mm^3^ isotropic) and smoothing (Gaussian kernel with 4 mm FWHM), we applied a bandpass filter (0.008 – 0.09 Hz) to the BOLD signal, discarded the first five frames to ensure signal equilibrium, and regressed out 24 parameters of motion^47^, as well as the mean signal from white matter and cerebrospinal fluid. Distorted images and artifacts were identified by low correlations (r < 0.9) between individual and a reference, preprocessed mean rs-fMRI image of 200 subjects. Visual inspection confirmed that failed spatial normalization or insufficient brain coverage in individual images were responsible for low correlation coefficients. We additionally excluded data from subjects with excessive in-scanner motion (maximum frame- wise rotation > 2° and movement > 3 mm).

### Measures of local brain activity and synchronicity

Three complementary maps of brain function, including measures of neuronal activity and synchronicity were derived from individual, preprocessed rs-fMRI data using the CONN toolbox. As a proxy for spontaneous local neuronal activity, we calculated voxel-wise maps of the fractional amplitude of low-frequency fluctuations (fALFF). fALFF is defined as the power ratio of low-frequency (0.008 – 0.09 Hz) oscillations to the total detectable frequency range in the BOLD signal^48^. Maps of local and global correlation (LCOR^49^ and GCOR^50^) represent proxies for both BOLD activity and synchronicity. LCOR is defined as the normalized sum of correlation coefficients of the BOLD signal in the voxel of interest with other voxels in its vicinity, with distances weighted by a Gaussian kernel (25 mm FWHM). GCOR is calculated in the same way as LCOR, but without distance-dependent weighting of the individual correlation coefficients. Thus, GCOR represents a measure of the global synchronicity of a voxel, whereas LCOR represents a measure of local coherence. Unlike fALFF, LCOR and GCOR do not depend on the amplitude of the BOLD signal, but rather on the similarity of the BOLD time series of all considered voxels. These three metrics provide a complementary characterization of the BOLD signal providing information about local neural activity as well as local and global functional connectivity.

### Aging effects and sex differences in fALFF, LCOR, and GCOR

We estimated voxel-wise aging effects by general linear modeling of (t-)contrast maps using a family-wise-error corrected voxel-wise threshold of P < 0.05 combined with a cluster-defining threshold of k > 20, including sex as a covariate. Using voxel-wise beta weights of the aging effects, we generated maps of annual changes in fALFF, LCOR and GCOR. Additionally, we assessed voxel-wise sex effects in fALFF, LCOR, and GCOR between women and men by general linear modeling of (t-)contrast maps using SPM12, including age and total intracranial volume (TIV) as covariates (thresholding as above).

### Spatial co-localization of brain function and neurotransmitter systems and effects of aging in the healthy population

We analyzed to what extent unthresholded group-level aging effects (maps of annual change) in fALFF, LCOR, and GCOR co-localize with specific neurotransmitter systems. For this, Spearman correlation coefficients were derived using the default Neuromorphometrics atlas (119 regions) estimating the similarity of aging effects in fALFF, LCOR, and GCOR with 19 distinct neurotransmitter maps as included in the JuSpace toolbox^25^. To approximate a normal distribution, correlation coefficients were Fisher’s z-transformed. We choose the Neuromorphometrics atlas as it provides a neuroanatomically plausible delineation of cortical and subcortical structures. As shown in our previous study^51^, the choice of atlas (with a comparable number of parcels) has a neglectable effect on the observed co-localization patterns of brain dysfunction and PET maps.

Included PET maps covered serotonergic receptors (5-HT1a^52^, 5-HT1b^53^, 5-HT2a^52^, 5-HT4^52^, 5-HT6^11^), dopaminergic receptors (D1^54^, D2^55^), histamine receptor 3 (H3^56^), dopamine uptake (DAT^24^), serotonin (SERT^52^), norepinephrine (NET^20^), vesicular acetylcholine (VAChT^57^) transporters, as well as the acetylcholinergic receptors M1^58^ and A4B2^59^, the glutamate receptors mGluR5^60^ and NMDA^61^, the cannabinoid CB1^62^, opioid µ^57^, and the GABAa^24^ receptor. Source publications, age, sex, and sample sizes characteristics of each PET map are provided in Supplementary Table 2. 95% confidence intervals of Spearman correlation coefficients were estimated using the Bonett-Wright^63^ procedure.

To better understand if and how these group level aging effects are also reflected in terms of the magnitude and the spread across all individual data, we re-computed the correlations on the single-subject level using individual measures of fALFF, LCOR, and GCOR. Prior to testing for aging effects across individual co-localizations, we examined whether Fisher’s z-transformed Spearman correlation coefficients of the healthy population were significantly different from a null distribution (one-sample t-test, alpha = 0.05). Aging effects on co- localization strengths were then estimated using linear regression analyses considering sex as a confound.

### Higher variation in co-localization between brain function and neurotransmitter systems with aging

Distinct aging trajectories from healthy aging to the effects of diseases and impairments are known to manifest in altered brain function. Depending on the underlying neurophysiological processes such changes are likely to be architecturally aligned with the spatial patterns of affected neurotransmitter systems. Correspondingly, one would expect to observe an increased variation in co-localization between brain function and receptor/transporter distribution in older as compared to younger participants. To test this hypothesis, we examined the heteroscedasticity in co-localization to identify neurotransmitter systems that are affected by such age-related brain functional changes.

For this, we performed two complementary tests. Using the White test, we first identified all pairs of brain function measures and PET maps whose correlation coefficients did not exhibit constant variance across age. In a post-hoc analysis, we tested for each co-localization pair with non-constant variance (P_FDR_ < 0.05) whether the variance for individuals in the upper third age-range is higher as compared to those in the lower third age-range using the Goldfeld-Quandt- test (one-sided, i.e., increasing variance). To account for potential confounding effects of sex between younger and older adults, we regressed out sex effects from the Fisher’s z-transformed Spearman correlation coefficients prior to the comparisons.

### Normative modeling of brain function - neurotransmitter co- localization and deviations in Parkinson’s disease

To model the effects of aging on the observed co-localization patterns we generated normative models based on the Fisher’s transformed correlation coefficients (co-localization strengths between neurotransmitter and rs-fMRI measures) derived from the healthy UK Biobank subjects using the PCNtoolkit^64^. To account for non-linear trajectories and non-Gaussian variance of the normal co-localization levels, we used Bayesian linear regression (5 knot basis splines and sinh-arcsinh warping) with age and sex as covariates (cf. Figure 1 for a methodological overview).

**Figure 1:**
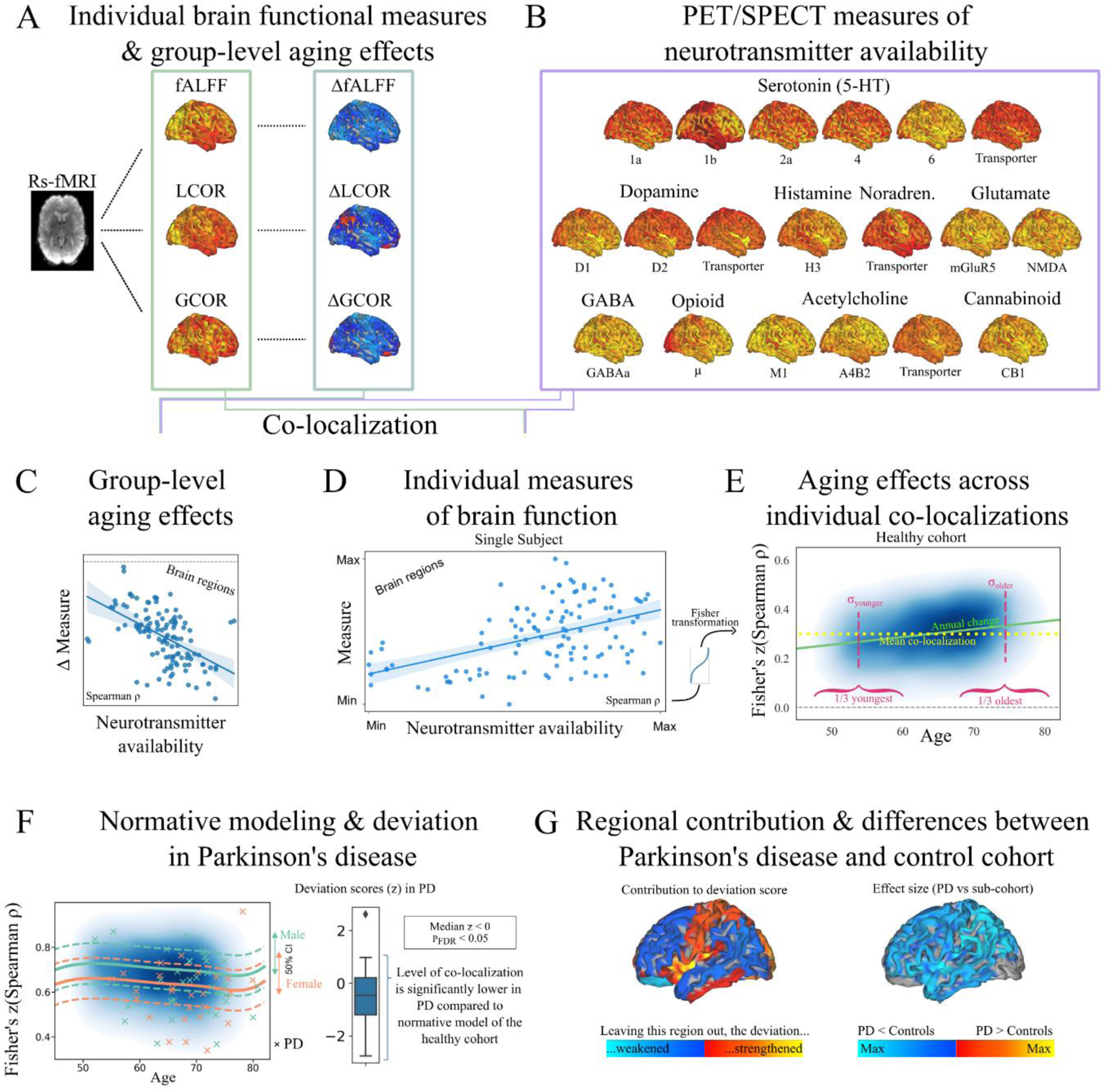
Methodological overview. We derived voxel-wise maps of fALFF, LCOR, and GCOR from individual rs-fMRI data (**A**). First, this data was used to explore group-level voxel-wise aging changes in fALFF, LCOR and GCOR in the healthy cohort (n_HC_=25,917, **A**, right column). Second, we used PET maps of 19 neurotransmitter systems (**B**) to calculate the spatial correlation (co-localization) with both, the group-level aging effects (**C**) and individual fALFF, LCOR, and GCOR (**D**). We Fisher (z-)transformed the Spearman correlation coefficients ρ to ensure a normal distribution and examined the effects of age on the co-localization data of the healthy cohort (**E**). The blue cloud illustrates co-localization strengths (kernel density estimation of all transformed Spearman correlation coefficients) of the healthy cohort. For each pair of measure and neurotransmitter map, we analyzed mean co-localizations (yellow), linear aging effects (green), as well as differences in variances across subjects in the youngest (44 - 60 years) and oldest (68 – 82 years) third (n_Both_ = 8,639; red). Vertical dashed red lines were added for illustration only and do not correspond to the actual variance of the respective subpopulation. Afterwards, we normatively modeled the co-localization strengths depending on age and sex (**F**, left) in order to calculate the deviation in subjects with manifest Parkinson’s disease (n_PD_=58; crosses). Here, we show the predicted means (solid lines) and 25% and 75% percentile (dashed lines) derived from the normative model for both men (blue) and women (orange). We analyzed whether the deviation (z-)scores of subjects with Parkinson’s disease were significantly different from a null distribution (**F**, box plot). In this example, the distribution was significantly below a null distribution, indicating that patients with PD exhibit lower Spearman correlation coefficients than the norm. Last, we quantified the mean regional contribution to the observed deviation score across subjects with Parkinson’s disease (**G**, left), as well as the functional differences in patients with Parkinson’s disease compared to an age- and sex-matched subcohort of healthy controls (n_HCmatched_ = 17,900; **G**, right). The functional differences were quantified by calculating the regional effect sizes (Cohen’s d).

Deviations (z-scores) from these normative aging models were derived per neurotransmitter map for subjects with PD. By comparing the deviation scores to a null distribution (t-test, alpha = 0.05), we identified neurotransmitter co-localization pairs for which patients with PD differed from normative models derived from healthy controls. For the significant (P_FDR_ < 0.05) deviations, we tested which regions contributed strongest to the observed deviations. To this end, we repeated the spatial correlation analyses in the data of PD using a leave-one-region-out approach^65^. As a measure of regional contribution to the deviation we calculated differences in squared correlation coefficients (Δρ^2^) between the reduced (n_Regions_ = 118: ρ_LOO_ ^2^) and the full (n_Regions_ = 119: ρ ^2^) set of regions. We set Δρ^2^ positive if omitting this specific region resulted in a more normal correlation coefficient (closer to the mean of the normative model), and negative if omitting led to stronger deviation from the normative model. We further evaluated whether these regional contributions to the observed co-localization strengths were spatially related to regional alterations in fALFF, LCOR, or GCOR by computing Pearson correlations between maps of Δρ^2^ and the regional effect size (Cohen’s d) in fALFF, LCOR, GCOR for differences between PD and an age- and sex-matched subgroup of HC (n = 17,400). In order to evaluate the extent of significant functional alterations in PD, regions with significant differences in fALFF, LCOR, and GCOR in PD compared with the matched controls were identified using the Mann-Whitney-U test.

All analyses were corrected for multiple comparisons using either the Benjamini-Hochberg procedure or, in case of highly inflated P-values due to large sample sizes, Bonferroni-Holm correction. To ensure that the observed functional co-localization effects are not driven by the underlying atrophy we repeated all analyses after regressing out the individual voxel-wise grey matter volumes (spatially smoothed with a 4 mm FWHM Gaussian kernel) from all functional maps.

## Results

### Demographic characterization

From a total pool of 30,035 subjects from the UK Biobank for which all necessary functional images and movement data was available, our analysis was based on data of 25,917 subjects for whom no diseases with known effect on brain function were reported.

Three subjects had no structural data available. Thus, the repetition of all analysis after controlling for age-related atrophy was performed on the data of 25,914 subjects. 75 subjects of the total pool had reported a diagnosis of Parkinson’s disease. In 58 of them, the first mention of the diagnosis (ICD-10: G20) was dated before their imaging session, so we classified them as “manifest” and included them for further analysis. In order to compare regional measures of brain function in PD with those of the healthy controls, we defined an age- and sex-matched subcohort consisting of 17,400 subjects (PD mean ± SD age in years: 68.6 ± 6.5, HC_matched_ mean ± SD age: 67.6 ± 6.0, P>0.26; 55.17% male PD and 51.7% male HC_matched_, χ^2^(1)=0.29, P=0.59).

An overview of the groups is provided in Table 1.

### Group-level aging effects in resting state measures and their co- localization to underlying neurotransmission

All three resting state measures decreased with aging in most cortical, sub-cortical, and cerebellar regions. Each measure showed a wide-spread but distinct spatial pattern of age-related alterations with few regional increases (Figure 2A, Supplementary Table 3 and Supplementary Table 4).

**Figure 2:**
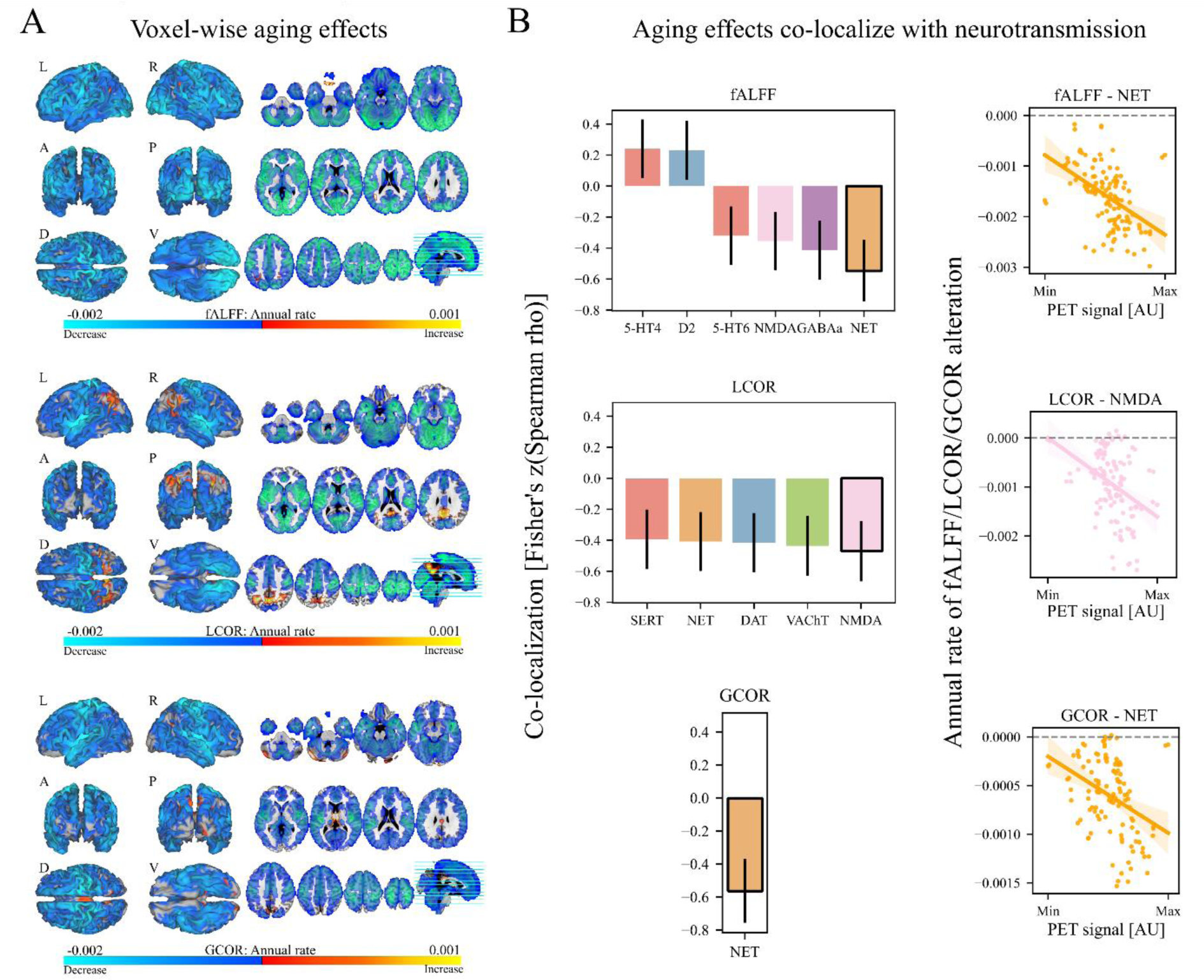
Group-level voxel-wise aging effects in each functional measure and associations with neurotransmitter systems. **A**: Colors in the voxel-wise plots of thresholded group-level aging effects indicate annual decrease (blue) or increase (red) in fALFF (top), LCOR (middle), and GCOR (bottom). **B, left column:** Significant (P_FDR_< 0.05) linear correlations between annual rate in brain functional measure and neurotransmission. Vertical black lines indicate the uncertainty of Fisher’s z-transformed Spearman correlation coefficient estimated according to Bonett & Wright^63^. **B, right column:** Exemplary scatter plots show how the annual change in fALFF, LCOR, or GCOR spatially correlate with the PET signal of specific neurotransmitter systems. Colors group receptors and transporters of the same neurotransmitter system, i.e., serotonin (red), dopamine (blue), acetylcholine (green), glutamate (pink) and GABA (purple), cannabinoid (mint), opioid (yellow), norepinephrine (orange), and histamine (turquoise).

Next, we aimed to understand if the topography of these age-related changes was linked to the distribution of specific neurotransmitter systems. For this, we derived voxel-wise maps of age-related linear (annual) changes in all three resting state measures and tested for their spatial co- localization with the neurotransmitter systems. The rate at which fALFF and LCOR changed during aging correlated significantly (P_FDR_<0.05) with serotonergic, dopaminergic, norepinephrinergic (also GCOR) and glutamate neurotransmission (Figure 2B, Supplementary Figures 1-2, Supplementary Tables 5-6). In addition, fALFF changes correlated with GABAergic and LCOR changes with cholinergic neurotransmission. Except for the correlation between fALFF changes and NMDA all findings remained significant after correcting for age-related atrophy. Scatter plots of the strongest correlations are displayed in Figure 2B. Results for voxel-wise sex differences are summarized in Supplementary Figures 3-5 and Supplementary Tables 7-10.

### Individual co-localization of resting state measures and neurotransmitter systems covaries with age

The extent to which a specific neurotransmitter system contributes to the measured brain function was evaluated by its correlation strength. We first computed the individual co- localization strengths between each subject’s resting state measure and each available neurotransmitter map. Due to the large cohort size, even very small effects in all resting state measures were significantly associated with the 19 PET maps (all P_Bonferroni-Holm_<0.0001, median absolute Spearman correlation coefficient ranged from 0.03 to 0.68) with the different neurotransmitter systems explaining between 0.1% and 46% of variance in the respective resting state measures (Supplementary Figure 6, Supplementary Tables 11-13, left columns). The direction of the correlations was highly similar across the three functional measures. Positive correlations were found for the norepinephrinergic, muscarinic, glutamatergic, GABAergic systems, as well as serotonergic receptors 5-HT1b, -2a, and -6. Negative correlations were found for the dopaminergic, histaminergic and opioid neurotransmitter systems, as well as for serotonin receptors 5-HT1a, 5-HT4 and serotonin and vesicular acetylcholine transporters (SERT and VAChT). All associations remained significant after controlling for age-related atrophy (Supplementary Tables 14-16). If age-related changes in brain functions are indeed primarily driven by specific neurotransmitter systems, the correlation coefficients should systematically (that is, in simplest approximation, linearly) in- or decrease during aging. Thus, we tested whether and to what extent aging effects and their co-localizations with neurotransmitter systems observed at the cohort level are also reflected in the individual co-localization strength. Most of the observed correlations were significantly associated with age explaining up to 3%, 4% and 1% of the co-localization strength between fALFF (with NMDA and SERT), LCOR (with SERT) and GCOR (with VAChT) and the respective neurotransmitter systems (Figure 3A, Supplementary Tables 11-13, middle columns). Correction for atrophy lowered the correlation strengths for most associations but the findings remained largely significant (Supplementary Tables 14-16, middle columns).

**Figure 3:**
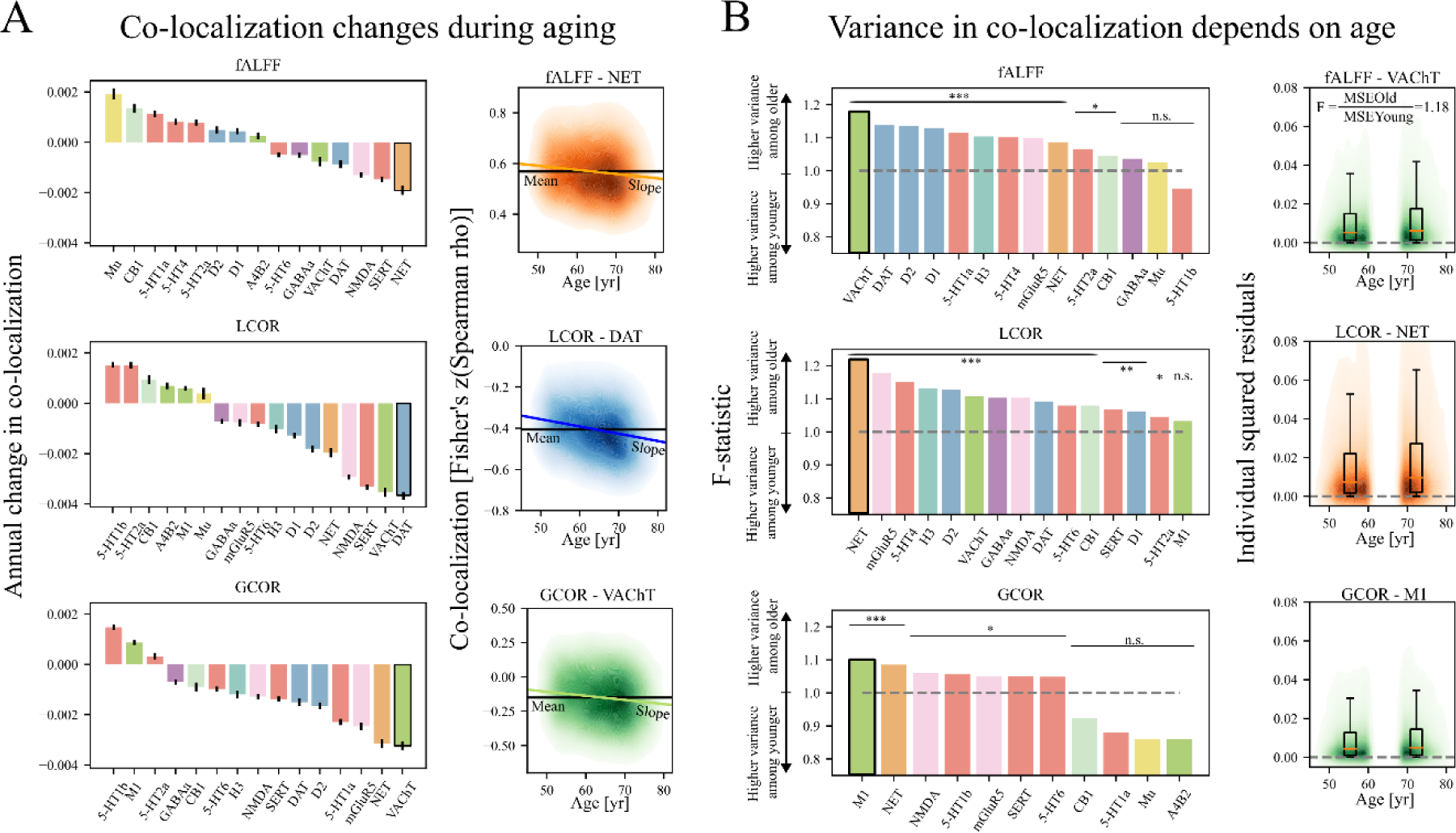
Single-subject co-localizations between brain functional measure and neurotransmitter systems depend on age. **A, left column:** Overview about all significant linear aging effects (P_Bonferroni-Holm_<0.0001) in the co-localization strengths (Fisher’s z-transformed Spearman correlation coefficients) between each pair of brain functional measure (fALFF, LCOR, GCOR) and neurotransmitter system. Error bars correspond to the standard error of parameter (slope) estimation. **A, right column:** Exemplary plots show how co-localization strengths between brain functional measures (fALFF: top, LCOR: middle, GCOR: bottom) and specific neurotransmitter systems depend on age. The black, horizontal line indicate the population mean co- localization (cf. Supplemd Spentary Figure 6 for an overview about all co-localization means). The colored clouds show the kernel density estimation of Fisher’s z-transformed Spearman correlation coefficients of the healthy cohort. The slope of the colored line (linear fits) corresponds to the bar plots in the left column. **B, left column:** Each plot shows the effect size (F- statistic) of mean squared error (MSE) differences between younger and older adults. We show bar plots regarding neurotransmitter systems whose co-localization with the respective brain functional measure was previously shown to be significantly non-constant (White-test, P_FDR_<0.05). All pairs of brain function and PET map whose co-localization variance was significantly (P_FDR_<0.05) different between the older and the younger subpopulations are highlighted by asterisks (*: P_FDR_<0.05; **: P_FDR_<0.01; ***: P_FDR_<0.001). F-statistics (MSE of older divided by MSE of younger adults) above 1 correspond to a larger variance in the older subpopulation. **B, right column:** Exemplary plots visualize the individual squared errors of the co-localizations between younger and older adults. Box plots visualize both distributions. Note, that per definition, the squared errors are positive. Due to the kernel density estimation of all squared errors the colored clouds exceed the null level. Colors group receptors and transporters of the same neurotransmitter system according to the same scheme as described in Figure 2.

### Variance in co-localization changes during aging

As aging may not only be associated with changes in average co-localization strengths but also with increased variance in such (i.e., due to yet undetected neurodegenerative processes in a subpopulation) we tested for such changes using a two-step procedure. Using the White-test, we first identified significant non-constant variance in co-localization strength. For fALFF and LCOR, non-constant variance was observed for all classes of neurotransmitters except for LCOR and opioid system. For GCOR, significant non-constant variance in co-localization was found for serotoninergic, norepinephrinergic, cannabinoid, opioid, glutamatergic and cholinergic neurotransmission (Supplementary Table 17, left columns). These effects remained significant after controlling for age-related atrophy except for GCOR and 5-HT1b, 5-HT6, and SERT (Supplementary Table 18, left columns). As the above analysis only detects differences in variance over age but not their direction, we aimed to better understand the respective findings by computing the Goldfeld-Quandt test comparing the co-localization variance between the youngest (44 – 60 years) and oldest (68 – 82 years) third (n_Both_ =8,639) of the study population for the significant associations identified using the White-test (Figure 3B).

For fALFF and LCOR, higher variability in co-localization was found in the elderly sub- population for serotonergic, dopaminergic, noradrenergic, histaminergic, cannabinoid, glutamatergic and cholinergic neurotransmission. In addition, for LCOR, we found significantly higher variance in the older population in co-localization with the GABA system. For GCOR, higher variability in co-localization was found regarding the serotonergic, noradrenergic, glutamatergic, and cholinergic system. For fALFF, 11 out of 14, for LCOR, 14 out of 15 and for GCOR, 7 out of 11 neurotransmitter co-localization pairs identified using the White-test displayed a higher variance in the older subpopulation (Figure 3B, Supplementary Table 17, right columns). The effects remained largely similar after controlling for atrophy (Supplementary Table 18, right columns).

### Deviations from normal co-localization in manifest Parkinson’s disease

Having established this reference for co-localization of normal age-related changes with different neurotransmitter systems, we now aimed to test if and how functional changes caused by progressive neurodegeneration deviate from the non-pathological co-localization patterns. For this, we adopted a normative modeling approach using the healthy aging population as a reference allowing for non-linear changes with age (models are visualized in Supplementary Figures 7-8). A UK Biobank subgroup of PD patients served as an example for the clinical relevance of our findings. For each PD patient, we derived individual deviation (z-)scores quantifying their deviation from the normative model. For fALFF, PD patients had a lower co- localization strength with serotonergic, GABAergic, muscarinic and glutamatergic neurotransmission (Figure 4D). For LCOR and GCOR, PD showed lower co-localizations with serotonergic, dopaminergic, GABAergic, histaminergic, norepinephrinergic, glutamatergic and cholinergic neurotransmitter systems (Figure 4E-F, Supplementary Table 19). The deviation in co-localization strength regarding LCOR and GABAa (illustrated in Figure 4A-B) correlated negatively with disease duration with higher deviations being indicative of longer disease duration (Pearson r = -0.38, P_FDR_ = 0.027; Figure 4C, Supplementary Tables 21-22). After atrophy correction, all deviations remained significant except for the LCOR-5-HT1b and LCOR-VAChT associations (Supplementary Table 20).

**Figure 4:**
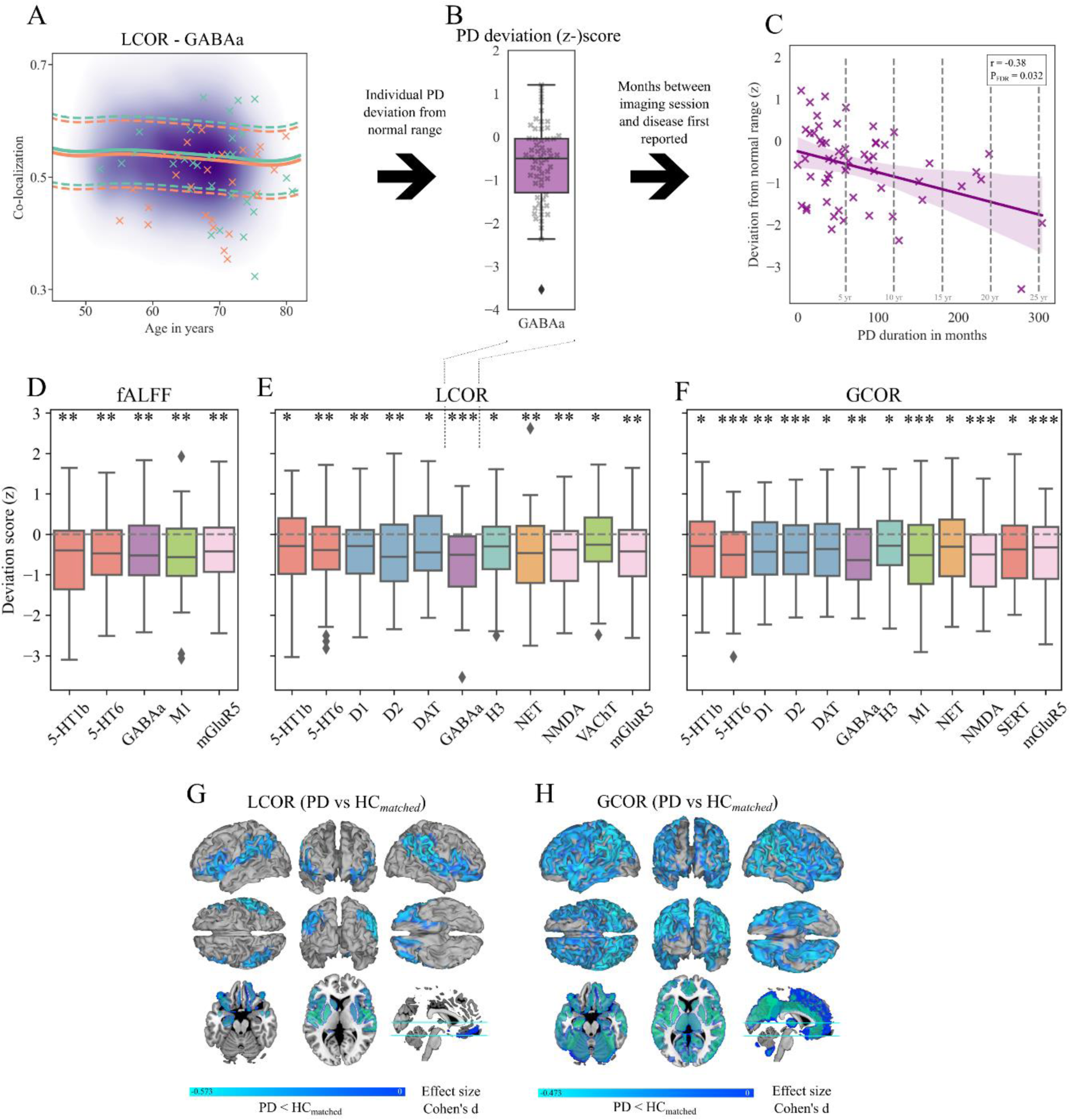
Subjects with PD deviate from normative models of co-localization between brain function and neurotransmitter systems. **A**: The purple cloud shows the kernel-density-plot of Fisher’s z-transformed Spearman correlation coefficients of the healthy cohort regarding the spatial correlation of LCOR and GABAa. Solid and dashed lines show the predicted mean and predicted 25% or 75% percentile of men (turquoise) and women (orange) derived from the normative models. Crosses indicate the co-localization levels of patients with PD. **B**: Box plot showing the significant deviation from the norm (null) in patients with PD. **C**: Deviation scores in PD are significantly correlated with disease duration. **D-F**: Box plots showing the deviation scores that were significantly different from the norm (null) regarding fALFF (**D**), LCOR (**E**), and GCOR (**F**). *, **, and *** indicate Bonferroni-Holm corrected significant deviations of the distributions from a null distribution with exact P<0.05, P<0.01, and P<0.001. Colors in box plots group receptors and transporters of the same neurotransmitter system according to the same scheme as described in Figure 2. **G-H:** Significant (P_FDR_<0.05) regional differences (effect sizes, Cohen’s d) between PD and the matched subgroup of healthy controls in LCOR (**G)** and GCOR (**H**). Lower values in PD are indicated by blue areas. Effect sizes of all region are provided in Supplementary Figures 17-19.

Lastly, we aimed to understand which regions contribute most to the observed co-localization alterations in PD. For this, we first analyzed how leaving out each region of the deployed atlas changed the observed correlations. For fALFF, main contributing regions to co-localization changes in PD were the basal ganglia, insula and occipital regions. For LCOR and GCOR, main contributing regions to co-localization changes in PD were the basal ganglia, subcallosal areas, thalamus (LCOR only), basal forebrain, as well as pre- and postcentral, insula and occipital regions (Supplementary Figures 9-11). The effects remained largely similar after controlling for atrophy (Supplementary Figures 12-14). Regional contribution to the deviations found regarding the glutamatergic system significantly correlated with regional alteration in PD in both synchronicity measures (P_FDR_<0.05, Supplementary Figures 15-16, Supplementary Tables 23-24). Effect sizes in regions that exhibit significant (P_FDR_<0.05) differences in LCOR and GCOR in PD compared to the age- and sex-matched subgroup of HC are provided in Figure 4G-H. Statistics of regional comparison of fALFF, LCOR, and GCOR in PD compared to the matched control group are provided in Supplementary Table 25-26. All regional effect sizes in fALFF, LCOR, and GCOR between PD and the age- and sex-matched subgroup of HC are shown in Supplementary Figures 17-19.

## Discussion

Here, we tested how age-related changes in commonly applied resting activity and connectivity measures co-localize with underlying neurotransmission. Consistent with previous studies of aging effects on brain function, we find widespread age-related decreases but also few increases in the three evaluated measures^2,3,66^. These age-related changes display a robust co-localization pattern with various major neurotransmitter systems, including monoamines, glutamate, choline, and GABA, at group- and single-subject level. Variance in the co-localization patterns of these systems increases over age. PD patients display significant deviations from typical age-related co-localization patterns in neurotransmitter systems related to the disease. The deviation in co-localization strength regarding the GABAergic system correlates with disease duration.

In line with most studies reporting aging effects in the brain, we find wide-spread age-related decreases in all three evaluated resting state measures^1–5^. The extent of the decreases is substantially higher in our study covering basically all of the brain with few exceptions as discussed below. Considering that these findings are based on over 25,000 subjects, the increased statistical power as compared to previous studies with at most few hundred participants is the most likely explanation for the observed discrepancy. In parallel, we observe spatially distinct age-related increases across the three evaluated resting state measures covering parietal, precuneal, thalamic, gyrus rectus and cerebellar regions. With respect to directionality, these findings are consistent with several previous studies, reporting presumably compensation-related increases in different connectivity metrics in the aging population^66,67^. Consistent with this interpretation, we find the local activity and connectivity to be increased in parietal regions involved in cross-modal sensory integration, potentially compensating for the prominent loss in modality-specific sensory processing during aging^68,69^.

Supporting the previously reported complex re-organization of the excitation/inhibition balance during aging^70^, we find the group-level aging effects on brain function to be associated with glutamatergic and GABAergic neurotransmission. The additional co-localization of the aging effects with monoaminergic and cholinergic systems may be supportive of the underlying changes to be related to learning, memory, and other higher cognitive functions affected by aging^7,15,67,71^. In contrast, the age-related global connectivity is primarily increased in thalamic and cerebellar regions and the topography of these changes only aligns with the norepinephrinergic neurotransmission. Both regions and in particular the thalamus show a high expression of norepinephrine receptors^72,73^. Whilst the modulatory role of norepinephrine in the cerebellum has been repeatedly associated with motor learning^74,75^, its contribution to aging is controversially discussed with its activity being associated with prevention but also acceleration of the production and accumulation of amyloid-β and tau across the brain^76^. On a functional level, these findings may originate in the functional decline of the norepinephrinergic system, which is considered as key factor in maintaining arousal and cognitive adaptation and control^77^_,78_.

When testing for co-localization of resting state measures with neurotransmission at the single subject level, we find age-related alterations in all three measures to co-localize primarily with monoaminergic neurotransmission. These findings are complemented by increases in variance observed for a variety of evaluated neurotransmitter systems. Considering the reportedly high prevalence of neuropathology in a cognitively normal elderly population^79,80^ the individual co- localization changes - aside with increased variance - may indeed reflect such yet undetected neurodegenerative processes. To test the sensitivity of the co-localization patterns to such neurodegenerative processes we further adopted a normative modeling approach. A major advantage of normative models is their ability to represent population heterogeneity in the phenotype under investigation by means of normalized deviation scores^64^. We applied this approach in patients with a diagnosis of Parkinson’s disease, which was previously linked to monoaminergic neurotransmission as well as more recently to a dysbalance of GABA and glutamate^81–83^. Indeed, across all three resting state metrics we find the co-localization patterns in PD patients to significantly deviate from age- and sex-adjusted normative models across a variety of neurotransmitter systems. Deviations in co-localization of local activity in PD are found primarily with respect to serotonergic, GABA, and glutamatergic neurotransmission, whilst deviations in co-localization of both local and global connectivity are also present with respect to dopamine neurotransmission. We find that only the deviation in co-localization strength of local connectivity with GABAa receptors predicts disease duration supporting the suggested relevance of GABA pathology for clinical progression^81^.

It is important to acknowledge that the studied cohort was recruited as a representative sample of healthy residents of the UK. Because the metrics we studied target local brain function, subjects with diseases primarily affecting brain structure or function were excluded. Given the prevalence for mild depressive symptoms of the UK population of 11 %^84^ and that 12.32 % of our analyzed participants of the UK Biobank have a reference to ICD-10: F32, their exclusion would have biased the cohort. Additional healthy control biases in the UK biobank^85^ include a high average socioeconomic status, low alcohol and tobacco consume. Although, we estimate that deviations from the overall population results are small with respect to these primary biases, further sampling of a more diverse population is needed for replication. In addition, clinical scores superior to disease duration as an estimate of disease severity should be used to strengthen evidence of the observed association with GABAergic neurotransmission. The use of PET maps from differently aged healthy populations may introduce a further bias into our findings as proteomics such as receptor and transporter distributions may change during aging^77,86^.

## Conclusions

Here, we provided a detailed overview on the effects of aging on macroscopic brain functioning as observed using rs-fMRI derived commonly measures of local activity and local and global connectivity. We link these age-related changes to the distribution of various major neurotransmitter systems demonstrating a decline in co-localization strength aside with increased variance during aging. By adopting a normative modeling approach, we further demonstrate on the example of Parkinson’s disease the feasibility of using co-localization strength as a sensitive measure of underlying neurodegeneration providing potentially valuable insight into the underlying neuropathological processes.

