## Supplementary material for "Resting state changes in aging and Parkinson’s disease are shaped by underlying neurotransmission – a normative modeling study"

**Supplementary Table 1**: List of exclusion criteria for the control group.

| | **Subgroup** | **ICD-10** |
|---|---|---|
| **Mental and Behavioural disorders (F-labeled)** | Physiological conditions | F01 - F03 |
| | Psychoactive substance use | F10 – F19 |
| | Schizophrenia, schizotypal, delusional, and schizoaffective disorder | F20 - F22, F25 |
| | Mood affective disorders | F31, F33 |
| | Intellectual disabilities | F70 – F72 |
| **Diseases of the central nervous system (G-labeled)** | Inflammatory diseases | G04 – G07 |
| | Systematic atrophies | G10 - G13, G14 |
| | Extrapyramidal and movement disorders | G20 – G25 |
| | Other degenerative diseases | G30-G32 |
| | Demyelinating diseases | G35-G37 |
| | Episodic and paroxysmal disorders | G40, G41, G45 |
| **Neoplasms (C-labeled)** | Malignant neoplasm of the brain | C71 |

**Supplementary Table 2**: Characteristics of neurotransmitter PET maps used.

| Atlas | Tracer | Sample size | % Male | Age (μ +- σ) | Source DOI |
|---|---|---|---|---|---|
| 5-HT1a | [11C]CUMI-101 | 8 | 37.50 | 28.4 +- 8.8 | 10.1523/JNEUROSCI.2830-16.2016 |
| 5-HT1b | [11C]P943 | 65 | 75.38 | 33.7 +- 9.7 | 10.1038/jcbfm.2009.195 |
| 5-HT2a | [11C]CIMBI-36 | 29 | 51.72 | 22.6 +- 2.7 | 10.1523/JNEUROSCI.2830-16.2016 |
| 5-HT4 | [11C]SB207145 | 59 | 69.49 | 25.9 +- 5.3 | 10.1523/JNEUROSCI.2830-16.2016 |
| 5-HT6 | [11C]GSK215083 | 30 | 100 | 36.6 +- 9 | 10.2967/jnumed.117.206516 |
| SERT | [11C]DASB | 100 | 29 | 25.1 +- 5.8 | 10.1523/JNEUROSCI.2830-16.2016 |
| D1 | [11C]SCH23390 | 13 | 46 | 33 +- 13 | 10.1007/s00259-017-3645-0 |
| D2 | [11C]FLB457 | 55 | 47.27 | 32.5 +- 9.7 | 10.1038/jcbfm.2014.237 |
| DAT | [123I]FP-CIT | 174 | 62.64 | 61 +- 11 | 10.1038/s41598-018-22444-0 |
| H3 | [11C]GSK189254 | 8 | 87.5 | 31.7 +- 9 | 10.1177/0271678X16650697 |
| NET | [11C]MRB | 77 | 64.94 | 33.4 +- 9.2 | 10.1002/syn.20696 |
| M1 | [11C]LSN3172176 | 24 | 54.17 | 40.50 +- 11.7 | 10.2967/jnumed.120.246967 |
| A4B2 | [18F]FLUBATINE | 30 | 66.67 | 33.5 +- 10.7 | 10.1016/j.neuroimage.2016.07.026 |
| VAChT | [18F]FEOBV | 18 | 27.78 | 66.8 +- 6.8 | 10.1038/mp.2017.183 |
| mGluR5 | [11C]ABP688 | 73 | 34.25 | 19.9 +- 3.04 | 10.1007/s00259-018-4252-4 |
| NMDA | [18F]GE-179 | 29 | 72 | 41 +- 13 | 10.1101/2021.12.04.21267226 |
| CB1 | [11C]OMAR | 77 | 63.64 | 30 +- 8.9 | 10.1038/jcbfm.2015.46 |
| Opioid mu | [11C]CARFENTANIL | 204 | 64.71 | 32.3 +- 10.8 | 10.1038/mp.2017.183 |
| GABAa | [11C]FLUMAZENIL | 6 | 100 | 43+-4 | 10.1038/s41598-018-22444-0 |

**Supplementary Table 3**: Regions covered by cluster of significant aging effects in fALFF, LCOR, and GCOR – *before atrophy correction*.

| Anatomical region | Cluster size | Cluster p-values (corrected) | Peak T-values | Peak MNI-Coordinates |
|---|---|---|---|---|
| **fALFF: Decreasing with age** | | | | |
| **Bilateral**: Superior frontal gyrus (dorsolateral), Middle temporal gyrus, Middle frontal gyrus, Postcentral gyrus, Middle occipital gyrus, Inferior temporal gyrus, Precuneus, Precentral gyrus, Superior temporal gyrus, Fusiform gyrus, Superior frontal gyrus (medial), Cerebellum (Crus 1), Inferior frontal gyrus (pars triangularis), Calcarine fissure and surrounding cortex, Inferior parietal gyrus, Lingual gyrus, Supplementary motor area, Cerebellum (8), Middle cingulate & paracingulate gyri, Insula, Cerebellum (6), Supramarginal gyrus, Superior parietal gyrus, Cerebellum (Crus 2), Cuneus, Angular gyrus, Rolandic operculum, Parahippocampal gyrus, Superior occipital gyrus, Inferior frontal gyrus (pars opercularis), Cerebellum (4,5), Superior temporal gyrus (pole), Putamen, Paracentral lobule, Hippocampus, Inferior occipital gyrus, Superior frontal gyrus (medial orbital), Cerebellum (9), Middle temporal gyrus (pole), Caudate nucleus, Anterior cingulate cortex (supracallosal), Gyrus rectus, Inferior frontal gyrus (pars orbitalis), Medial orbital gyrus, Anterior cingulate cortex (pregenual), Posterior orbital gyrus, Anterior orbital gyrus, Cerebellum (7b), Vermis (4,5), Olfactory cortex, Vermis (6), Amygdala, Posterior cingulate gyrus, Vermis (8), Heschl's gyrus, Lateral orbital gyrus, Thalamus (pulvinar medial), Vermis (9), Anterior cingulate cortex (subgenual), Ventral striatum, Vermis (7), Cerebellum (3), Vermis (3), Thalamus (mediodorsal medial magnocellular), Pallidum, Thalamus (ventral posterolateral), Thalamus (mediodorsal lateral parvocellular), Thalamus (ventral lateral), Vermis (1,2), Thalamus (pulvinar inferior), Thalamus (pulvinar anterior), Vermis (10), Thalamus (pulvinar lateral), Thalamus (lateral posterior), Thalamus (anteroventral nucleus), Thalamus (lateral geniculate), Thalamus (intralaminar), Substantia nigra pars compacta, Raphe nucleus, Thalamus (medial geniculate), Red nucleus, Cerebellum (10)<br>**Left:** Ventral tegmental area, Thalamus (ventral anterior) | 48649 | <0.001 | 49.3 | **45 0 -3** |
| **fALFF: Increasing with age** | | | | |
| **Left**: Angular gyrus, Middle occipital gyrus, Inferior parietal gyrus, Middle temporal gyrus | 118 | <0.001 | 20.83 | **-36 -54 24** |
| **Right**: Supramarginal gyrus, Angular gyrus | 33 | <0.001 | 11.39 | **30 -45 36** |
| **LCOR: Decreasing with age** | | | | |
| **Bilateral**: Middle temporal gyrus, Superior frontal gyrus (dorsolateral), Postcentral gyrus, Precentral gyrus, Superior temporal gyrus, Fusiform gyrus, Middle frontal gyrus, Calcarine fissure and surrounding cortex, Lingual gyrus, Supplementary motor area, Inferior frontal gyrus (pars triangularis), Inferior temporal gyrus, Superior frontal gyrus (medial), Cerebellum (8), Insula, Cerebellum (6), Cerebellum (Crus 1), Middle occipital gyrus, Rolandic operculum, Cuneus, Middle cingulate & paracingulate gyri, Parahippocampal gyrus, Supramarginal gyrus, Inferior frontal gyrus (pars opercularis), Cerebellum (4,5), Cerebellum (Crus 2), Putamen, Superior temporal gyrus (pole), Paracentral lobule, Inferior parietal gyrus, Hippocampus, Inferior occipital gyrus, Precuneus, Cerebellum (9), Caudate nucleus, Superior occipital gyrus, Anterior cingulate cortex (supracallosal), Superior parietal gyrus, Inferior frontal gyrus (pars orbitalis), Anterior cingulate cortex (pregenual), Middle temporal | 38017 | <0.001 | 44.64 | **45 3 -6** |

| | | | | |
|---|---|---|---|---|
| gyrus (pole), Superior frontal gyrus (medial orbital), Posterior orbital gyrus, Cerebellum (7b), Vermis (4,5), Gyrus rectus, Vermis (6), Amygdala, Olfactory cortex, Vermis (8), Heschl's gyrus, Vermis (9), Thalamus (pulvinar medial), Anterior cingulate cortex (subgenual), Vermis (7), Cerebellum (3), Vermis (3), Medial orbital gyrus, Thalamus (mediodorsal medial magnocellular), Ventral striatum, Pallidum, Lateral orbital gyrus, Thalamus (ventral posterolateral), Thalamus (mediodorsal lateral parvocellular), Anterior orbital gyrus, Thalamus (ventral lateral), Vermis (1,2), Thalamus (pulvinar inferior), Thalamus (pulvinar anterior), Vermis (10), Thalamus (pulvinar lateral), Thalamus (lateral posterior), Thalamus (anteroventral nucleus), Thalamus (lateral geniculate), Thalamus (intralaminar), Substantia nigra pars compacta, Raphe nucleus, Thalamus (medial geniculate), Red nucleus, Cerebellum (10)<br>**Left:** Angular gyrus, Ventral tegmental area, Thalamus (ventral anterior) | | | | |
| **LCOR: Increasing with age** | | | | |
| **Bilateral**: Precuneus, Angular gyrus, Inferior parietal gyrus, Posterior cingulate gyrus, Middle occipital gyrus, Middle cingulate & paracingulate gyri, Superior parietal gyrus, Cuneus, Supramarginal gyrus, Middle temporal gyrus, Superior occipital gyrus, Postcentral gyrus<br>**Left**: Calcarine fissure and surrounding cortex | 1796 | <0.001 | 24.87 | **-33 -57 33** |
| **Left**: Gyrus rectus | 27 | <0.001 | 8.25 | **-6 21 -30** |
| **GCOR: Decreasing with age** | | | | |
| **Bilateral**: Middle temporal gyrus, Superior frontal gyrus (dorsolateral), Postcentral gyrus, Middle frontal gyrus, Precentral gyrus, Superior temporal gyrus, Inferior temporal gyrus, Superior frontal gyrus (medial), Fusiform gyrus, Inferior frontal gyrus (pars triangularis), Supplementary motor area, Middle occipital gyrus, Inferior parietal gyrus, Cerebellum (8), Insula, Lingual gyrus, Precuneus, Supramarginal gyrus, Middle cingulate & paracingulate gyri, Superior parietal gyrus, Cerebellum (6), Calcarine fissure and surrounding cortex, Rolandic operculum, Inferior frontal gyrus (pars opercularis), Cerebellum (4,5), Parahippocampal gyrus, Superior temporal gyrus (pole), Cuneus, Paracentral lobule, Superior occipital gyrus, Hippocampus, Middle temporal gyrus (pole), Angular gyrus, Cerebellum (9), Superior frontal gyrus (medial orbital), Anterior cingulate cortex (supracallosal), Inferior occipital gyrus, Putamen, Inferior frontal gyrus (pars orbitalis), Cerebellum (Crus 1), Anterior cingulate cortex (pregenual), Cerebellum (Crus 2), Gyrus rectus, Vermis (4,5), Caudate nucleus, Posterior orbital gyrus, Cerebellum (7b), Vermis (6), Amygdala, Vermis (8), Heschl's gyrus, Anterior cingulate cortex (subgenual), Olfactory cortex, Lateral orbital gyrus, Vermis (9), Vermis (7), Vermis (3), Cerebellum (3), Posterior cingulate gyrus, Thalamus (pulvinar medial), Ventral striatum, Pallidum, Medial orbital gyrus, Thalamus (mediodorsal medial magnocellular), Thalamus (ventral posterolateral), Vermis (1,2), Thalamus (pulvinar inferior), Anterior orbital gyrus, Thalamus (pulvinar anterior), Vermis (10), Thalamus (lateral geniculate), Thalamus (pulvinar lateral), Thalamus (intralaminar), Raphe nucleus, Thalamus (medial geniculate), Thalamus (mediodorsal lateral parvocellular), Cerebellum (10)<br>**Left**: Thalamus (lateral posterior) | 38929 | <0.001 | 33.04 | **45 9 -9** |
| **GCOR: Increasing with age** | | | | |
| **Bilateral**: Thalamus (mediodorsal medial magnocellular), Caudate nucleus, Thalamus (ventral lateral), Thalamus (pulvinar medial), Thalamus (mediodorsal lateral parvocellular), | 206 | <0.001 | 22.15 | **-3 -3 9** |

| | | | | |
|---|---|---|---|---|
| Thalamus (anteroventral nucleus), Thalamus (lateral posterior), Hippocampus, Thalamus (intralaminar)<br>**Left**: Thalamus (ventral anterior)<br>**Right**: Thalamus (pulvinar anterior), Thalamus (pulvinar lateral), Posterior cingulate gyrus | | | | |
| **Left**: Angular gyrus, Middle occipital gyrus | 45 | <0.001 | 16.7 | **-33 -57 33** |
| **Left**: Cerebellum (Crus 1), Cerebellum (Crus 2), Cerebellum (6) | 350 | <0.001 | 15.56 | **-45 -72 -30** |
| **Bilateral**: Precuneus, Superior parietal gyrus<br>**Right**: Angular gyrus, Superior occipital gyrus, Cuneus | 217 | <0.001 | 13.5 | **18 -60 45** |
| **Bilateral**: Posterior cingulate gyrus<br>**Right**: Middle cingulate & paracingulate gyri | 70 | <0.001 | 12.07 | **0 -24 27** |
| **Right**: Cerebellum (Crus 1), Lingual gyrus, Cerebellum (6), Calcarine fissure and surrounding cortex | 164 | <0.001 | 10.23 | **12 -99 -12** |

**Supplementary Table 4**: Regions covered by cluster of significant aging effects in fALFF, LCOR, and GCOR – *after atrophy correction*.

| **Anatomical region** | **Cluster size** | **Cluster p-values (corrected)** | **Peak T-values** | **Peak MNI-Coordinates** |
|---|---|---|---|---|
| **fALFF: Decreasing with age** | | | | |
| **Bilateral**: Superior frontal gyrus (dorsolateral), Middle frontal gyrus, Middle temporal gyrus, Postcentral gyrus, Middle occipital gyrus, Precentral gyrus, Inferior temporal gyrus, Precuneus, Superior temporal gyrus, Superior frontal gyrus (medial), Fusiform gyrus, Cerebellum (Crus 1), Inferior parietal gyrus, Inferior frontal gyrus (pars triangularis), Calcarine fissure and surrounding cortex, Lingual gyrus, Supplementary motor area, Cerebellum (8), Middle cingulate & paracingulate gyri, Insula, Cerebellum (6), Supramarginal gyrus, Superior parietal gyrus, Cerebellum (Crus 2), Angular gyrus, Cuneus, Rolandic operculum, Parahippocampal gyrus, Inferior frontal gyrus (pars opercularis), Superior occipital gyrus, Cerebellum (4,5), Superior temporal gyrus (pole), Putamen, Paracentral lobule, Hippocampus, Inferior occipital gyrus, Superior frontal gyrus (medial orbital), Middle temporal gyrus (pole), Cerebellum (9), Caudate nucleus, Anterior cingulate cortex (supracallosal), Gyrus rectus, Inferior frontal gyrus (pars orbitalis), Medial orbital gyrus, Anterior cingulate cortex (pregenual), Posterior orbital gyrus, Anterior orbital gyrus, Cerebellum (7b), Vermis (4,5), Posterior cingulate gyrus, Olfactory cortex, Vermis (6), Amygdala, Vermis (8), Heschl's gyrus, Lateral orbital gyrus, Thalamus (pulvinar medial), Vermis (9), Anterior cingulate cortex (subgenual), Ventral striatum, Vermis (7), Cerebellum (3), Vermis (3), Thalamus (mediodorsal medial magnocellular), Pallidum, Thalamus (ventral posterolateral), Thalamus (mediodorsal lateral parvocellular), Thalamus (ventral lateral), Vermis (1,2), Thalamus (pulvinar inferior), Thalamus (pulvinar anterior), Vermis (10), Thalamus (pulvinar lateral), Thalamus (lateral posterior), Thalamus (anteroventral nucleus), Thalamus (lateral geniculate), Thalamus (intralaminar), | 49470 | <0.001 | 42.83 | **0 -36 24** |

| | | | | |
|---|---|---|---|---|
| Substantia nigra pars compacta, Raphe nucleus, Thalamus (medial geniculate), Red nucleus, Cerebellum (10).<br><br>**Left**: Ventral tegmental area, Thalamus (ventral anterior) | | | | |
| **fALFF: Increasing with age** | | | | |
| **Bilateral**: Gyrus rectus | 30 | <0.001 | 7.87 | **6 21 -30** |
| **LCOR: Decreasing with age** | | | | |
| **Bilateral:** Middle temporal gyrus, Superior frontal gyrus (dorsolateral), Postcentral gyrus, Precentral gyrus, Superior temporal gyrus, Middle frontal gyrus, Fusiform gyrus, Calcarine fissure and surrounding cortex, Lingual gyrus, Supplementary motor area, Inferior frontal gyrus (pars triangularis), Inferior temporal gyrus, Superior frontal gyrus (medial), Cerebellum (8), Insula, Cerebellum (6), Middle occipital gyrus, Cerebellum (Crus 1), Rolandic operculum, Middle cingulate & paracingulate gyri, Cuneus, Inferior frontal gyrus (pars opercularis), Parahippocampal gyrus, Supramarginal gyrus, Cerebellum (4,5), Putamen, Superior temporal gyrus (pole), Cerebellum (Crus 2), Paracentral lobule, Hippocampus, Inferior parietal gyrus, Inferior occipital gyrus, Cerebellum (9), Caudate nucleus, Precuneus, Superior occipital gyrus, Anterior cingulate cortex (supracallosal), Inferior frontal gyrus (pars orbitalis), Superior parietal gyrus, Anterior cingulate cortex (pregenual), Middle temporal gyrus (pole), Superior frontal gyrus (medial orbital), Posterior orbital gyrus, Cerebellum (7b), Vermis (4,5), Vermis (6), Gyrus rectus, Amygdala, Olfactory cortex, Vermis (8), Heschl's gyrus, Vermis (9), Thalamus (pulvinar medial), Anterior cingulate cortex (subgenual), Vermis (7), Cerebellum (3), Ventral striatum, Vermis (3), Thalamus (mediodorsal medial magnocellular), Medial orbital gyrus, Pallidum, Lateral orbital gyrus, Thalamus (ventral posterolateral), Anterior orbital gyrus, Thalamus (mediodorsal lateral parvocellular), Thalamus (ventral lateral), Vermis (1,2), Thalamus (pulvinar inferior), Thalamus (pulvinar anterior), Vermis (10), Thalamus (pulvinar lateral), Thalamus (lateral posterior), Thalamus (anteroventral nucleus), Thalamus (lateral geniculate), Thalamus (intralaminar), Substantia nigra pars compacta, Raphe nucleus, Thalamus (medial geniculate), Red nucleus, Cerebellum (10)<br><br>**Left**: Angular gyrus, Ventral tegmental area, Thalamus (ventral anterior) | 38027 | <0.001 | 34.38 | **48 6 -9** |
| **LCOR: Increasing with age** | | | | |
| **Bilateral**: Precuneus, Angular gyrus, Inferior parietal gyrus, Posterior cingulate gyrus, Middle occipital gyrus, Middle cingulate & paracingulate gyri, Superior parietal gyrus, Cuneus<br><br>**Left**: Middle temporal gyrus, Calcarine fissure and surrounding cortex<br><br>**Right**: Supramarginal gyrus, Superior occipital gyrus | 1670 | <0.001 | 16.87 | **-33 -60 33** |
| **Left**: Gyrus rectus | 30 | <0.001 | 8.23 | **-6 21 -30** |
| **Right**: Anterior orbital gyrus, Medial orbital gyrus | 26 | <0.001 | 7.7 | **30 51 -18** |
| **GCOR: Decreasing with age** | | | | |
| **Bilateral**: Middle temporal gyrus, Superior frontal gyrus (dorsolateral), Postcentral gyrus, Middle frontal gyrus, Precentral gyrus, Superior temporal gyrus, Inferior temporal gyrus, Superior frontal gyrus (medial), Inferior frontal gyrus (pars triangularis), Fusiform gyrus, | 39066 | <0.001 | 28.32 | **45 9 -12** |

| | | | | |
|---|---|---|---|---|
| Middle occipital gyrus, Supplementary motor area, Inferior parietal gyrus, Insula, Cerebellum (8), Lingual gyrus, Precuneus, Supramarginal gyrus, Middle cingulate & paracingulate gyri, Superior parietal gyrus, Calcarine fissure and surrounding cortex, Cerebellum (6), Rolandic operculum, Inferior frontal gyrus (pars opercularis), Cerebellum (4,5), Parahippocampal gyrus, Cuneus, Superior temporal gyrus (pole), Paracentral lobule, Superior occipital gyrus, Hippocampus, Middle temporal gyrus (pole), Angular gyrus, Cerebellum (9), Inferior occipital gyrus, Superior frontal gyrus (medial orbital), Anterior cingulate cortex (supracallosal), Inferior frontal gyrus (pars orbitalis), Putamen, Anterior cingulate cortex (pregenual), Cerebellum (Crus 2), Gyrus rectus, Cerebellum (Crus 1), Vermis (4,5), Posterior orbital gyrus, Caudate nucleus, Cerebellum (7b), Vermis (6), Amygdala, Vermis (8), Heschl's gyrus, Olfactory cortex, Anterior cingulate cortex (subgenual), Lateral orbital gyrus, Vermis (9), Vermis (3), Cerebellum (3), Vermis (7), Posterior cingulate gyrus, Thalamus (pulvinar medial), Medial orbital gyrus, Ventral striatum, Pallidum, Thalamus (mediodorsal medial magnocellular), Thalamus (ventral posterolateral), Vermis (1,2), Thalamus (pulvinar inferior), Anterior orbital gyrus, Thalamus (pulvinar anterior), Thalamus (lateral geniculate), Vermis (10), Thalamus (pulvinar lateral), Thalamus (intralaminar), Raphe nucleus, Thalamus (medial geniculate), Thalamus (mediodorsal lateral parvocellular)<br>**Left**: Cerebellum (10), Thalamus (lateral posterior | | | | |
| **GCOR: Increasing with age** | | | | |
| **Bilateral**: Thalamus (mediodorsal medial magnocellular), Caudate nucleus, Thalamus (ventral lateral), Thalamus (pulvinar medial), Thalamus (mediodorsal lateral parvocellular), Thalamus (anteroventral nucleus), Thalamus (lateral posterior), Hippocampus, Thalamus (intralaminar)<br>**Left**: Thalamus (ventral anterior)<br>**Right**: Thalamus (pulvinar anterior), Posterior cingulate gyrus | 206 | <0.001 | 19.61 | **3 -12 12** |
| **Left:** Cerebellum (Crus 1), Cerebellum (Crus 2), Cerebellum (6) | 380 | <0.001 | 15.75 | **-45 -72 -30** |
| **Bilateral**: Precuneus<br>**Left**: Superior parietal gyrus<br>**Right**: Superior occipital gyrus, Cuneus | 180 | <0.001 | 12.52 | **15 -63 42** |
| **Bilateral**: Posterior cingulate gyrus´<br>**Right**: Middle cingulate & paracingulate gyri | 68 | <0.001 | 12.14 | **0 -24 27** |
| **Right**: Cerebellum (Crus 1), Cerebellum (6), Cerebellum (Crus 2), Lingual gyrus, Fusiform gyrus, Calcarine fissure and surrounding cortex | 199 | <0.001 | 11.78 | **12 -99 -12** |

**Supplementary Table 5**: Co-localization of **aging in brain functional measures** (unthresholded voxel-wise maps of annual change) and neurotransmitter systems – *before atrophy correction*.

| | fALFF | | LCOR | | GCOR | |
|---|---|---|---|---|---|---|
| Neurotransmitter system | Rho | $P_{FDR}$ | Rho | $P_{FDR}$ | Rho | $P_{FDR}$ |
| 5-HT1a | 0.1 | 0.6343 | 0.1 | 0.6216 | 0.1 | 0.1429 |
| 5-HT1b | -0.15 | 0.5644 | -0.15 | 0.5798 | -0.15 | 0.659 |
| 5-HT2a | -0.12 | 0.592 | -0.12 | 0.5798 | -0.12 | 0.9195 |
| 5-HT4 | **0.24** | **0.0399** | 0.24 | 0.5798 | 0.24 | 0.5239 |
| 5-HT6 | **-0.31** | **0.0044** | -0.31 | 0.3043 | -0.31 | 0.5239 |
| A4B2 | -0.05 | 0.7282 | -0.05 | 0.9755 | -0.05 | 0.9195 |
| CB1 | 0.17 | 0.5644 | 0.17 | 0.5798 | 0.17 | 0.9195 |
| D1 | 0.05 | 0.7282 | 0.05 | 0.8403 | 0.05 | 0.1178 |
| D2 | **0.23** | **0.0421** | 0.23 | 0.5798 | 0.23 | 0.9739 |
| DAT | 0.02 | 0.8802 | **0.02** | **0.0006** | 0.02 | 0.7522 |
| GABAa | **-0.39** | **0.0038** | -0.39 | 0.5798 | -0.39 | 0.1985 |
| H3 | <0.01 | 0.9941 | <0.01 | 0.6836 | <0.01 | 0.9195 |
| M1 | -0.14 | 0.5644 | -0.14 | 0.5798 | -0.14 | 0.7522 |
| Mu | 0.31 | 0.2216 | 0.31 | 0.5798 | 0.31 | 0.8353 |
| NET | **-0.5** | **0.0038** | **-0.5** | **0.0209** | **-0.5** | **0.0171** |
| NMDA | **-0.34** | **0.0421** | **-0.34** | **0.0033** | -0.34 | 0.9195 |
| SERT | -0.09 | 0.5644 | **-0.09** | **0.0006** | -0.09 | 0.9195 |
| VAChT | -0.03 | 0.833 | **-0.03** | **0.0006** | -0.03 | 0.1411 |
| mGluR5 | -0.3 | 0.1183 | -0.3 | 0.6216 | -0.3 | 0.1411 |

Bold numbers highlight significant correlations. Rho: Spearman correlation coefficient.

**Supplementary Table 6**: Co-localization of **aging in brain functional measures** (unthresholded voxel-wise maps of annual change) and neurotransmitter systems – *after atrophy correction*.

| | fALFF | | LCOR | | GCOR | |
|---|---|---|---|---|---|---|
| Neurotransmitter system | Rho | $P_{FDR}$ | Rho | $P_{FDR}$ | Rho | $P_{FDR}$ |
| 5-HT1a | 0.09 | 0.6088 | 0.09 | 0.6497 | 0.09 | 0.1254 |
| 5-HT1b | -0.17 | 0.4982 | -0.17 | 0.5704 | -0.17 | 0.6017 |
| 5-HT2a | -0.14 | 0.6017 | -0.14 | 0.5704 | -0.14 | 0.8934 |
| 5-HT4 | **0.23** | **0.0407** | 0.23 | 0.5704 | 0.23 | 0.6017 |
| 5-HT6 | **-0.29** | **0.0095** | -0.29 | 0.3616 | -0.29 | 0.6017 |
| A4B2 | -0.06 | 0.6088 | -0.06 | 0.9926 | -0.06 | 0.9099 |
| CB1 | 0.15 | 0.6017 | 0.15 | 0.5704 | 0.15 | 0.8934 |
| D1 | 0.07 | 0.6017 | 0.07 | 0.9573 | 0.07 | 0.0703 |
| D2 | **0.24** | **0.0337** | 0.24 | 0.5704 | 0.24 | 0.9762 |
| DAT | 0.05 | 0.6387 | **0.05** | **0.0006** | 0.05 | 0.6017 |
| GABAa | **-0.38** | **0.0038** | -0.38 | 0.5704 | -0.38 | 0.2346 |
| H3 | <0.01 | 0.9796 | <0.01 | 0.729 | <0.01 | 0.8934 |
| M1 | -0.13 | 0.6017 | -0.13 | 0.5704 | -0.13 | 0.7047 |
| Mu | 0.29 | 0.2878 | 0.29 | 0.5704 | 0.29 | 0.8381 |
| NET | **-0.48** | **0.0085** | **-0.48** | **0.0228** | **-0.48** | **0.0114** |
| NMDA | -0.31 | 0.0731 | **-0.31** | **0.0057** | -0.31 | 0.8934 |
| SERT | -0.08 | 0.6017 | **-0.08** | **0.0006** | -0.08 | 0.8934 |
| VAChT | -0.01 | 0.9232 | **-0.01** | **0.0006** | -0.01 | 0.1254 |
| mGluR5 | -0.28 | 0.1664 | -0.28 | 0.6497 | -0.28 | 0.1254 |

Bold numbers highlight significant correlations. Rho: Spearman correlation coefficient.

**Supplementary Table 7**: Anatomical regions covered by sex-differences (T-contrasts) - *before atrophy correction*.

| Anatomical region | Cluster size | Cluster p-values (corrected) | Peak T-values | Peak MNI-Coordinates |
|---|---|---|---|---|
| **FALFF: Female < Male** | | | | |
| **Bilateral**: Postcentral gyrus, middle frontal gyrus, precentral gyrus, middle occipital gyrus, superior temporal gyrus (incl. pole), calcarine fissure and surrounding cortex, dorsolateral and medial superior frontal gyrus, lingual gyrus, middle temporal gyrus (incl. pole), inferior parietal gyrus, insula, supplementary motor area, fusiform gyrus, supramarginal gyrus, inferior frontal gyrus (pars triangularis & orbitalis), inferior temporal gyrus, superior parietal gyrus, rolandic operculum, cuneus, superior occipital gyrus, cerebellum (4,5,6,7b,8,9,10) crus1, crus2), vermis (4,5,6,8) middle cingulate gyrus, paracentral lobule, inferior occipital gyrus, putamen, precuneus, orbital gyrus (posterior, medial, anterior, lateral), parahippocampal gyrus, anterior cingulate cortex (subgenual, pregenual, supracallosal), hippocampus, Heschl gyrus, angular gyrus, amygdala, gyrus rectus, thalamus (pulvinar medial, mediodorsal medial magnocellular, ventral posterolateral, mediodorsal lateral parvocellular, pulvinar lateral, ventral lateral, pulvinar anterior, anteroventral nucleus, intralaminar, lateral geniculate, medial geniculate), caudate, pallidum, olfactory cortex, ventral striatum, raphe nucleus (dorsal), substantia nigra (pars compacta), red nucleus,<br>**Left**: Thalamus (ventral anterior), Ventral tegmental area | 29898 | <0.001 | 29.31 | **-57 -3 15** |
| **Right:** Cerebellum (7b, 8, crus1, crus2) | 58 | <0.001 | 19.43 | **54 -54 -42** |
| **Right:** Caudate | 47 | <0.001 | 11.11 | **18 -9 27** |
| **FALFF: Female > Male** | | | | |
| **Bilateral:** Cerebellum 9, Vermis 9 | 182 | <0.001 | 23.01 | **0 -54 -51** |
| **Bilateral:** Frontal gyrus (medial pars orbitalis, superior dorsolateral & medial), gyrus rectus, anterior cingulate gyrus (subgenual), orbital gyrus (anterior & medial)<br>**Right**: Anterior cingulate gyrus (pregenual) | 497 | <0.001 | 22.65 | **-15 69 9** |
| **Bilateral:** Precuneus, cingulate gyrus (middle & posterior), calcarine fissure and surrounding cortex, vermis (4,5)<br>**Left:** Cuneus<br>**Right:** Lingual gyrus | 974 | <0.001 | 20.82 | **6 -57 21** |
| **Right:** Temporal gyrus (inferior, middle, superior) | 574 | <0.001 | 19.08 | **66 -15 -15** |
| **Bilateral:** Superior frontal gyrus (medial)<br>**Left:** Frontal gyrus (superior, middle), supplementary motor area | 369 | <0.001 | 15.14 | **-27 18 57** |
| **Bilateral:** Superior frontal gyrus (medial)<br>**Left:** Frontal gyrus (superior, middle), supplementary motor area | 390 | <0.001 | 15.03 | **30 18 57** |

| | | | | |
|---|---|---|---|---|
| **Bilateral:** Vermis (7)<br>**Right:** Cerebellum (6, 7b, 8, crus1, crus2) | 421 | <0.001 | 12.17 | **27 -75 -33** |
| **Left:** Temporal gyrus (middle & inferior) | 319 | <0.001 | 11.98 | **-69 -27 -9** |
| **Left:** Cerebellum (6,7b,8, crus1, crus2) | 315 | <0.001 | 11.63 | **-36 -63 -39** |
| **Left:** Parietal gyrus (superior & inferior), occipital gyrus (middle), angular gyrus | 104 | <0.001 | 11.51 | **-36 -72 39** |
| **Right:** Gyrus rectus, orbital gyrus (medial) | 24 | <0.001 | 10.29 | **9 21 -24** |
| **Right:** Angular gyrus, occipital gyrus (middle) | 51 | <0.001 | 10.00 | **45 -66 33** |
| **Left:** Frontal gyrus (superior, middle, inferior pars orbitalis) | 37 | <0.001 | 9.96 | **-27 36 -9** |
| **Left:** Orbital gyrus (medial), gyrus rectus | 20 | <0.001 | 9.93 | **-12 21 -24** |
| **Left:** Parahippocampal gyrus, fusiform gyrus, temporal gyrus (inferior) | 34 | <0.001 | 9.05 | **-33 -39 -9** |
| **Right:** Parahippocampal gyrus, fusiform gyrus, hippocampus | 28 | <0.001 | 8.98 | **33 -36 -12** |
| **Right:** Frontal gyrus (middle, inferior pars orbitalis), orbital gyrus (anterior) | 25 | <0.001 | 8.00 | **33 39 -6** |
| **LCOR: Female < Male** | | | | |
| **Bilateral**: Middle frontal gyrus, Postcentral gyrus, Middle occipital gyrus, Middle temporal gyrus, Precentral gyrus, Superior frontal gyrus (dorsolateral), Superior temporal gyrus, Inferior parietal gyrus, Inferior frontal gyrus (pars triangularis), Precuneus, Fusiform gyrus, Supplementary motor area, Calcarine fissure and surrounding cortex, Lingual gyrus, Insula, Supramarginal gyrus, Superior parietal gyrus, Cerebellum (Crus 1), Cerebellum (6), Middle cingulate & paracingulate gyri, Cerebellum (8), Superior frontal gyrus (medial), Cuneus, Angular gyrus, Cerebellum (Crus 2), Rolandic operculum, Inferior frontal gyrus (pars opercularis), Superior occipital gyrus, Cerebellum (4,5), Inferior temporal gyrus, Putamen, Superior temporal gyrus (pole), Paracentral lobule, Inferior occipital gyrus, Parahippocampal gyrus, Inferior frontal gyrus (pars orbitalis), Cerebellum (9), Middle temporal gyrus (pole), Cerebellum (7b), Anterior cingulate cortex (supracallosal), Anterior cingulate cortex (pregenual), Posterior orbital gyrus, Vermis (4,5), Hippocampus, Vermis (8), Heschl's gyrus, Vermis (6), Amygdala, Lateral orbital gyrus, Anterior orbital gyrus, Vermis (7), Cerebellum (3), Vermis (9), Pallidum, Thalamus (pulvinar medial), Medial orbital gyrus, Vermis (3), Thalamus (mediodorsal medial magnocellular), Thalamus (ventral posterolateral), Caudate nucleus, Thalamus (mediodorsal lateral parvocellular), Thalamus (pulvinar anterior), Thalamus (ventral lateral), Thalamus (intralaminar), Thalamus (pulvinar lateral), Thalamus (pulvinar inferior), Thalamus (medial geniculate), Thalamus (lateral geniculate), Olfactory cortex, Vermis (10), Vermis (1,2), Cerebellum (10)<br>**Right:** Gyrus rectus, Superior frontal gyrus (medial orbital), Thalamus (lateral posterior), Thalamus (anteroventral nucleus), Posterior cingulate gyrus | 37792 | <0.001 | 30.05 | **48 -15 15** |
| **Bilateral**: Substantia nigra pars compacta, Red nucleus<br>**Left**: Ventral tegmental area | | <0.001 | 9.79 | **0 -18 -12** |
| **LCOR: Female > Male** | | | | |
| **Bilateral**: Superior frontal gyrus (medial orbital), Superior frontal gyrus (dorsolateral), Gyrus rectus, Caudate nucleus, Superior frontal gyrus (medial), Medial orbital gyrus, Anterior cingulate cortex (subgenual), Anterior orbital gyrus, Middle frontal gyrus, Olfactory cortex, Thalamus (lateral posterior)<br>**Left**: Anterior cingulate cortex (pregenual), Anterior cingulate cortex (supracallosal), Ventral striatum, Thalamus (pulvinar medial), Thalamus (ventral lateral), Thalamus (anteroventral nucleus) | 1465 | <0.001 | 34.53 | **15 69 6** |
| **Right**: Middle temporal gyrus, Inferior temporal gyrus, Cerebellum (Crus 1), Cerebellum (6), Superior temporal gyrus, Fusiform gyrus, Cerebellum (Crus 2), Middle temporal gyrus (pole) | 491 | <0.001 | 22.7 | 69 -21 -18 |
| **Left**: Hippocampus, Parahippocampal gyrus | 51 | <0.001 | 16.17 | -33 -39 -3 |

| | | | | |
|---|---|---|---|---|
| **Left**: Inferior temporal gyrus, Middle temporal gyrus, Cerebellum (Crus 1), Cerebellum (7b), Cerebellum (6) | 508 | <0.001 | 15.84 | -66 -27 -21 |
| **Right**: Hippocampus, Parahippocampal gyrus | 46 | <0.001 | 14.41 | 33 -36 0 |
| **Bilateral**: Cerebellum (9), Vermis (9) | 47 | <0.001 | 13.55 | 0 -48 -48 |
| **Right**: Thalamus (mediodorsal medial magnocellular) | 24 | <0.001 | 13.14 | 0 -18 0 |
| **Bilateral:** Precuneus, Posterior cingulate gyrus, Middle cingulate & paracingulate gyri, Calcarine fissure and surrounding cortex<br>**Left**: Cuneus | 269 | <0.001 | 12.06 | -9 -54 18 |
| **Right**: Medial orbital gyrus, Olfactory cortex, Gyrus rectus, Posterior orbital gyrus | 83 | <0.001 | 11.58 | 9 15 -24 |
| **Left**: Medial orbital gyrus, Olfactory cortex, Posterior orbital gyrus, Gyrus rectus, Parahippocampal gyrus | 66 | <0.001 | 10.09 | -12 15 -24 |
| **GCOR: Female < Male** | | | | |
| **Bilateral**: Middle temporal gyrus, Middle frontal gyrus, Superior frontal gyrus (dorsolateral), Postcentral gyrus, Precuneus, Middle occipital gyrus, Precentral gyrus, Superior temporal gyrus, Inferior parietal gyrus, Superior frontal gyrus (medial), Inferior frontal gyrus (pars triangularis), Fusiform gyrus, Supplementary motor area, Calcarine fissure and surrounding cortex, Middle cingulate & paracingulate gyri, Lingual gyrus, Inferior temporal gyrus, Insula, Supramarginal gyrus, Cerebellum (8), Superior parietal gyrus, Angular gyrus, Cuneus, Cerebellum (6), Rolandic operculum, Inferior frontal gyrus (pars opercularis), Superior occipital gyrus, Cerebellum (4,5), Superior temporal gyrus (pole), Putamen, Parahippocampal gyrus, Paracentral lobule, Inferior occipital gyrus, Anterior cingulate cortex (supracallosal), Inferior frontal gyrus (pars orbitalis), Cerebellum (9), Superior frontal gyrus (medial orbital), Anterior cingulate cortex (pregenual), Hippocampus, Middle temporal gyrus (pole), Cerebellum (Crus 2), Anterior orbital gyrus, Posterior orbital gyrus, Vermis (4,5), Cerebellum (7b), Posterior cingulate gyrus, Caudate nucleus, Gyrus rectus, Medial orbital gyrus, Heschl's gyrus, Vermis (8), Vermis (6), Cerebellum (Crus 1), Amygdala, Lateral orbital gyrus, Thalamus (pulvinar medial), Vermis (9), Cerebellum (3), Ventral striatum, Anterior cingulate cortex (subgenual), Pallidum, Vermis (7), Vermis (3), Thalamus (mediodorsal medial magnocellular), Olfactory cortex, Thalamus (ventral posterolateral), Thalamus (mediodorsal lateral parvocellular), Thalamus (pulvinar inferior), Thalamus (pulvinar anterior), Thalamus (ventral lateral), Thalamus (pulvinar lateral), Thalamus (intralaminar), Thalamus (lateral geniculate), Substantia nigra pars compacta, Vermis (1,2), Thalamus (medial geniculate), Thalamus (lateral posterior), Thalamus (anteroventral nucleus), Vermis (10), Red nucleus, Raphe nucleus, Cerebellum (10)<br>**Left**: Ventral tegmental area, Thalamus (ventral anterior) | 43115 | <0.001 | 26.55 | **45 -15 9** |
| **GCOR: Female > Male** | | | | |
| **Right**: Caudate nucleus | 65 | <0.001 | 18.82 | **12 0 21** |
| **Left**: Hippocampus | 42 | <0.001 | 17.68 | **-30 -39 3** |
| **Left**: Caudate nucleus, Thalamus (lateral posterior), Thalamus (pulvinar medial) | 63 | <0.001 | 16.76 | **-12 -6 21** |
| **Right**: Superior frontal gyrus (medial), Superior frontal gyrus (dorsolateral), Superior frontal gyrus (medial orbital) | 60 | <0.001 | 15.08 | **15 69 3** |
| **Right**: Hippocampus | 33 | <0.001 | 14.35 | **33 -36 0** |
| **Left**: Superior frontal gyrus (dorsolateral), Superior frontal gyrus (medial), Superior frontal gyrus (medial orbital) | 47 | <0.001 | 13.77 | **-12 69 6** |
| **Left**: Cerebellum (Crus 1), Cerebellum (Crus 2), Cerebellum (7b) | 305 | <0.001 | 12.84 | **-51 -66 -30** |

| | | | | |
|---|---|---|---|---|
| **Right**: Cerebellum (Crus 1), Cerebellum (Crus 2), Cerebellum (7b), Inferior temporal gyrus, Cerebellum (6) | 194 | <0.001 | 12.78 | **54 -60 -33** |
| **Bilateral**: Cerebellum (9) | 25 | <0.001 | 10.74 | **0 -51 -54** |
| **Left**: Medial orbital gyrus, Posterior orbital gyrus, Olfactory cortex, Parahippocampal gyrus, Gyrus rectus | 29 | <0.001 | 8.71 | **-12 12 -24** |
| **Right**: Medial orbital gyrus, Olfactory cortex, Gyrus rectus | 23 | <0.001 | 8.02 | **12 12 -24** |

**Supplementary Table 8**: Anatomical regions covered by sex-differences (T-contrasts) - *after atrophy correction.*

| Anatomical region | Cluster size | Cluster p-values (corrected) | Peak T-values | Peak MNI-Coordinates |
|---|---|---|---|---|
| **FALFF: Female < Male** | | | | |
| **Bilateral**: Postcentral gyrus, Middle frontal gyrus, Precentral gyrus, Middle occipital gyrus, Superior temporal gyrus, Calcarine fissure and surrounding cortex, Superior frontal gyrus (dorsolateral), Lingual gyrus, Middle temporal gyrus, Inferior parietal gyrus, Insula, Fusiform gyrus, Supplementary motor area, Supramarginal gyrus, Inferior frontal gyrus (pars triangularis), Inferior temporal gyrus, Superior parietal gyrus, Rolandic operculum, Cuneus, Superior occipital gyrus, Cerebellum (6), Inferior frontal gyrus (pars opercularis), Middle cingulate & paracingulate gyri, Cerebellum (8), Paracentral lobule, Cerebellum (4,5), Inferior occipital gyrus, Superior temporal gyrus (pole), Putamen, Precuneus, Cerebellum (Crus 1), Superior frontal gyrus (medial), Middle temporal gyrus (pole), Inferior frontal gyrus (pars orbitalis), Parahippocampal gyrus, Posterior orbital gyrus, Vermis (4,5), Cerebellum (9), Cerebellum (Crus 2), Anterior orbital gyrus, Caudate nucleus, Hippocampus, Anterior cingulate cortex (supracallosal), Medial orbital gyrus, Anterior cingulate cortex (pregenual), Vermis (8), Heschl's gyrus, Angular gyrus, Amygdala, Vermis (6), Gyrus rectus, Cerebellum (7b), Lateral orbital gyrus, Thalamus (pulvinar medial), Thalamus (mediodorsal medial magnocellular), Cerebellum (3), Pallidum, Olfactory cortex, Vermis (9), Vermis (7), Ventral striatum, Vermis (3), Thalamus (ventral posterolateral), Thalamus (mediodorsal lateral parvocellular), Superior frontal gyrus (medial orbital), Thalamus (pulvinar inferior), Thalamus (ventral lateral), Thalamus (pulvinar anterior), Thalamus (anteroventral nucleus), Thalamus (pulvinar lateral), Thalamus (intralaminar), Vermis (10), Thalamus (lateral geniculate), Thalamus (lateral posterior), Substantia nigra pars compacta, Raphe nucleus, Anterior cingulate cortex (subgenual), Thalamus (medial geniculate), Red nucleus, Cerebellum (10)<br>**Left**: Ventral tegmental area, Thalamus (ventral anterior) | 30583 | <0.001 | 29.07 | **48 -15 9** |
| **fALFF: Female > Male** | | | | |
| **Bilateral**: Superior frontal gyrus (medial orbital), Superior frontal gyrus (dorsolateral), Superior frontal gyrus (medial), Gyrus rectus, Anterior cingulate cortex (subgenual), Anterior orbital gyrus, Medial orbital gyrus<br>**Left**: Anterior cingulate cortex (pregenual), Olfactory cortex | 494 | <0.001 | 22.53 | **15 69 6** |
| **Bilateral**: Precuneus, Middle cingulate & paracingulate gyri, Posterior cingulate gyrus, Cuneus, Calcarine fissure and surrounding cortex, Vermis (4,5)<br>**Right**: Lingual gyrus | 934 | <0.001 | 21.06 | **6 -57 21** |
| **Bilateral:** Cerebellum (9) | 147 | <0.001 | 20.43 | **-3 -54 -48** |
| **Right**: Middle temporal gyrus, Inferior temporal gyrus, Superior temporal gyrus | 597 | <0.001 | 20.27 | **63 -15 -15** |

| | | | | |
|---|---|---|---|---|
| **Bilateral**: Superior frontal gyrus (medial)<br>**Right**: Superior frontal gyrus (dorsolateral), Middle frontal gyrus, Supplementary motor area | 361 | <0.001 | 14.73 | **30 18 57** |
| **Left**: Superior frontal gyrus (dorsolateral), Middle frontal gyrus, Supplementary motor area, Superior frontal gyrus (medial) | 356 | <0.001 | 14.72 | **-30 18 57** |
| **Left**: Middle temporal gyrus, Inferior temporal gyrus | 384 | <0.001 | 12.76 | **-69 -27 -9** |
| **Right**: Cerebellum (Crus 1), Cerebellum (Crus 2), Cerebellum (7b), Cerebellum (6), Cerebellum (8) | 286 | <0.001 | 11.02 | **30 -75 -33** |
| **Right**: Gyrus rectus, Medial orbital gyrus | 23 | <0.001 | 10.87 | **12 21 -27** |
| **Right**: Angular gyrus, Middle occipital gyrus | 49 | <0.001 | 10.75 | **45 -66 30** |
| **Left**: Inferior parietal gyrus, Angular gyrus, Middle occipital gyrus, Superior parietal gyrus | 80 | <0.001 | 10.49 | **-36 -72 39** |
| **Left**: Cerebellum (Crus 2), Cerebellum (Crus 1), Cerebellum (7b), Cerebellum (8), Cerebellum (6) | 178 | <0.001 | 10.25 | **-30 -75 -33** |
| **Left**: Inferior frontal gyrus (pars orbitalis), Middle frontal gyrus, Superior frontal gyrus (dorsolateral) | 24 | <0.001 | 7.68 | **-27 39 -9** |
| **LCOR: Female < Male** | | | | |
| **Bilateral**: Middle frontal gyrus, Postcentral gyrus, Middle occipital gyrus, Middle temporal gyrus, Precentral gyrus, Superior frontal gyrus (dorsolateral), Superior temporal gyrus, Inferior parietal gyrus, Inferior frontal gyrus (pars triangularis), Precuneus, Fusiform gyrus, Calcarine fissure and surrounding cortex, Supplementary motor area, Lingual gyrus, Insula, Supramarginal gyrus, Superior parietal gyrus, Cerebellum (Crus 1), Cerebellum (6), Middle cingulate & paracingulate gyri, Cerebellum (8), Superior frontal gyrus (medial), Cuneus, Cerebellum (Crus 2), Angular gyrus, Rolandic operculum, Inferior frontal gyrus (pars opercularis), Superior occipital gyrus, Cerebellum (4,5), Inferior temporal gyrus, Superior temporal gyrus (pole), Putamen, Paracentral lobule, Inferior occipital gyrus, Parahippocampal gyrus, Inferior frontal gyrus (pars orbitalis), Cerebellum (9), Middle temporal gyrus (pole), Anterior cingulate cortex (supracallosal), Cerebellum (7b), Anterior cingulate cortex (pregenual), Hippocampus, Posterior orbital gyrus, Vermis (4,5), Vermis (6), Vermis (8), Heschl's gyrus, Amygdala, Anterior orbital gyrus, Lateral orbital gyrus, Vermis (9), Vermis (7), Cerebellum (3), Pallidum, Thalamus (pulvinar medial), Medial orbital gyrus, Vermis (3), Thalamus (mediodorsal medial magnocellular), Thalamus (ventral posterolateral), Thalamus (mediodorsal lateral parvocellular), Caudate nucleus, Thalamus (pulvinar anterior), Thalamus (ventral lateral), Thalamus (pulvinar inferior), Thalamus (intralaminar), Vermis (10), Thalamus (pulvinar lateral), Thalamus (lateral geniculate), Thalamus (medial geniculate), Olfactory cortex, Vermis (1,2), Cerebellum (10)<br>**Right**: Gyrus rectus, Superior frontal gyrus (medial orbital), Thalamus (lateral posterior), Thalamus (anteroventral nucleus), Posterior cingulate gyrus | 38244 | <0.001 | 32.67 | **45 -15 12** |
| **Bilateral:** Substantia nigra pars compacta, Red nucleus<br>**Left**: Ventral tegmental area | 36 | <0.001 | 9.74 | **0 -18 -12** |

| | | | | |
|---|---|---|---|---|
| **LCOR: Female > Male** | | | | |
| **Bilateral**: Superior frontal gyrus (medial orbital), Superior frontal gyrus (dorsolateral), Gyrus rectus, Superior frontal gyrus (medial), Anterior cingulate cortex (subgenual), Medial orbital gyrus, Anterior orbital gyrus, Middle frontal gyrus, Olfactory cortex<br>**Left**: Anterior cingulate cortex (pregenual) | 1171 | <0.001 | 34.43 | **-15 69 6** |
| **Right**: Middle temporal gyrus, Inferior temporal gyrus, Cerebellum (Crus 1), Cerebellum (6), Superior temporal gyrus, Fusiform gyrus, Cerebellum (Crus 2), Middle temporal gyrus (pole) | 487 | <0.001 | 22.36 | **69 -21 -18** |
| **Bilateral**: Caudate nucleus, Thalamus (lateral posterior)<br>**Left:** Ventral striatum, Thalamus (pulvinar medial), Thalamus (ventral lateral), Thalamus (anteroventral nucleus) | 270 | <0.001 | 17 | **-3 -6 15** |
| **Left**: Middle temporal gyrus, Inferior temporal gyrus, Cerebellum (Crus 1), Cerebellum (6), Cerebellum (7b) | 489 | <0.001 | 15.81 | **-66 -24 -21** |
| **Left**: Hippocampus, Parahippocampal gyrus | 46 | <0.001 | 14.74 | **-33 -36 -3** |
| **Right**: Hippocampus, Parahippocampal gyrus | 37 | <0.001 | 13.22 | **33 -36 0** |
| **Bilateral**: Precuneus, Posterior cingulate gyrus, Middle cingulate & paracingulate gyri, Calcarine fissure and surrounding cortex<br>**Left**: Cuneus | 269 | <0.001 | 12.87 | **-9 -54 18** |
| **Right**: Medial orbital gyrus, Olfactory cortex, Gyrus rectus, Posterior orbital gyrus | 80 | <0.001 | 11.73 | **9 15 -27** |
| **Left:** Medial orbital gyrus, Olfactory cortex, Posterior orbital gyrus, Gyrus rectus, Parahippocampal gyrus | 69 | <0.001 | 10.75 | **-12 15 -24** |
| **Bilateral:** Cerebellum (9) | 22 | <0.001 | 10.71 | **0 -48 -48** |
| **GCOR: Female < Male** | | | | |
| **Bilateral**: Middle temporal gyrus, Middle frontal gyrus, Superior frontal gyrus (dorsolateral), Postcentral gyrus, Precuneus, Middle occipital gyrus, Precentral gyrus, Superior temporal gyrus, Inferior parietal gyrus, Superior frontal gyrus (medial), Inferior frontal gyrus (pars triangularis), Fusiform gyrus, Supplementary motor area, Calcarine fissure and surrounding cortex, Middle cingulate & paracingulate gyri, Lingual gyrus, Inferior temporal gyrus, Insula, Supramarginal gyrus, Cerebellum (8), Superior parietal gyrus, Angular gyrus, Cuneus, Cerebellum (6), Rolandic operculum, Inferior frontal gyrus (pars opercularis), Superior occipital gyrus, Cerebellum (4,5), Superior temporal gyrus (pole), Putamen, Parahippocampal gyrus, Paracentral lobule, Inferior occipital gyrus, Anterior cingulate cortex (supracallosal), Cerebellum (9), Inferior frontal gyrus (pars orbitalis), Superior frontal gyrus (medial orbital), Hippocampus, Anterior cingulate cortex (pregenual), Middle temporal gyrus (pole), Cerebellum (Crus 2), Anterior orbital gyrus, Posterior orbital gyrus, Vermis (4,5), Cerebellum (7b), Posterior cingulate gyrus, Caudate nucleus, Gyrus rectus, Medial orbital gyrus, Cerebellum (Crus 1), Vermis (6), Heschl's gyrus, Amygdala, Vermis (8), Lateral orbital gyrus, Thalamus (pulvinar medial), Vermis (9), Anterior cingulate cortex (subgenual), Cerebellum (3), Ventral striatum, Vermis (7), Pallidum, Vermis (3), Thalamus (mediodorsal medial magnocellular), Olfactory cortex, Thalamus (ventral posterolateral), Thalamus (mediodorsal lateral parvocellular), Thalamus (pulvinar inferior), Thalamus (pulvinar anterior), Thalamus (ventral lateral), Thalamus (pulvinar lateral), Vermis (10), Thalamus (intralaminar), Thalamus (lateral geniculate), Substantia nigra pars compacta, Vermis (1,2), Thalamus (medial geniculate), Thalamus (anteroventral nucleus), Red nucleus, Thalamus (lateral posterior), Raphe nucleus, Cerebellum (10)<br>**Left**: Ventral tegmental area, Thalamus (ventral anterior) | 43309 | <0.001 | 28.94 | **45 -15 9** |

| | | | | |
|---|---|---|---|---|
| **GCOR: Female > Male** | | | | |
| **Right**: Caudate nucleus | 63 | <0.001 | 17.97 | **12 0 21** |
| **Left**: Hippocampus | 41 | <0.001 | 16.75 | **-30 -39 3** |
| **Left** Caudate nucleus, Thalamus (lateral posterior), Thalamus (pulvinar medial) | 60 | <0.001 | 16.34 | **-15 -12 24** |
| **Right:** Superior frontal gyrus (medial), Superior frontal gyrus (dorsolateral), Superior frontal gyrus (medial orbital) | 60 | <0.001 | 15.82 | **15 69 3** |
| **Left**: Superior frontal gyrus (dorsolateral), Superior frontal gyrus (medial), Superior frontal gyrus (medial orbital) | 46 | <0.001 | 13.85 | **-12 69 6** |
| **Right**: Hippocampus | 32 | <0.001 | 13.59 | **30 -36 6** |
| **Left:** Cerebellum (Crus 1), Cerebellum (Crus 2), Cerebellum (7b) | 295 | <0.001 | 12.94 | **-45 -42 -39** |
| **Right**: Cerebellum (Crus 1), Cerebellum (Crus 2), Cerebellum (7b), Inferior temporal gyrus, Cerebellum (6) | 170 | <0.001 | 12.55 | **54 -63 -33** |
| **Left**: Medial orbital gyrus, Posterior orbital gyrus, Olfactory cortex, Gyrus rectus | 29 | <0.001 | 8.35 | **-12 12 -24** |
| **Right**: Medial orbital gyrus, Olfactory cortex, Gyrus rectus | 22 | <0.001 | 8.07 | **12 12 -24** |

**Supplementary Table 9**: Co-localization of **sex differences in brain functional measures** (unthresholded voxel-wise T-maps) and neurotransmitter systems – *before atrophy correction*.

| | fALFF | | LCOR | | GCOR | |
|---|---|---|---|---|---|---|
| Neurotransmitter system | Rho | $P_{FDR}$ | Rho | $P_{FDR}$ | Rho | $P_{FDR}$ |
| 5-HT1a | 0.01 | 0.9855 | 0.01 | 0.8937 | 0.01 | 0.5794 |
| 5-HT1b | 0.12 | 0.6393 | 0.12 | 0.4761 | 0.12 | 0.1514 |
| 5-HT2a | 0.08 | 0.7707 | 0.08 | 0.8735 | 0.08 | 0.5682 |
| 5-HT4 | -0.13 | 0.3764 | -0.13 | 0.0574 | -0.13 | 0.0809 |
| 5-HT6 | 0.21 | 0.084 | **0.21** | **0.0019** | **0.21** | **0.0005** |
| A4B2 | <0.01 | 0.9855 | <0.01 | 0.2618 | <0.01 | 0.2429 |
| CB1 | -0.12 | 0.7283 | -0.12 | 0.9522 | -0.12 | 0.7879 |
| D1 | -0.07 | 0.6826 | -0.07 | 0.252 | -0.07 | 0.9835 |
| D2 | 0.01 | 0.9855 | 0.01 | 0.554 | 0.01 | 0.6187 |
| DAT | 0.11 | 0.4722 | 0.11 | 0.8735 | 0.11 | 0.6838 |
| GABAa | 0.25 | 0.0522 | **0.25** | **0.0009** | **0.25** | **0.0005** |
| H3 | -0.01 | 0.9855 | -0.01 | 0.9522 | -0.01 | 0.5682 |
| M1 | 0.13 | 0.6254 | 0.13 | 0.5209 | 0.13 | 0.2896 |
| Mu | -0.22 | 0.5527 | -0.22 | 0.5209 | -0.22 | 0.5794 |
| NET | **0.57** | **0.0038** | **0.57** | **0.0009** | **0.57** | **0.0005** |
| NMDA | 0.29 | 0.1086 | 0.29 | 0.252 | 0.29 | 0.113 |
| SERT | 0.21 | 0.084 | 0.21 | 0.9522 | 0.21 | 0.5794 |
| VAChT | 0.23 | 0.057 | 0.23 | 0.252 | 0.23 | 0.5794 |
| mGluR5 | 0.25 | 0.2611 | **0.25** | **0.0043** | **0.25** | **0.0005** |

Bold numbers highlight significant correlations. Rho: Spearman correlation coefficient.

**Supplementary Table 10**: Co-localization of **sex differences in brain functional measures** (unthresholded voxel-wise T-maps) and neurotransmitter systems – *after atrophy correction*.

| | fALFF | | LCOR | | GCOR | |
|---|---|---|---|---|---|---|
| Neurotransmitter system | Rho | $P_{FDR}$ | Rho | $P_{FDR}$ | Rho | $P_{FDR}$ |
| 5-HT1a | -0.01 | 0.9831 | -0.01 | 0.9168 | -0.01 | 0.5968 |
| 5-HT1b | 0.1 | 0.6522 | 0.1 | 0.52 | 0.1 | 0.1485 |
| 5-HT2a | 0.07 | 0.8329 | 0.07 | 0.9019 | 0.07 | 0.5706 |
| 5-HT4 | -0.15 | 0.2344 | -0.15 | 0.0483 | -0.15 | 0.0711 |
| 5-HT6 | 0.2 | 0.1194 | **0.2** | **0.0025** | **0.2** | **0.0005** |
| A4B2 | <0.01 | 0.9831 | <0.01 | 0.2677 | <0.01 | 0.2705 |
| CB1 | -0.14 | 0.6522 | -0.14 | 0.9168 | -0.14 | 0.8053 |
| D1 | -0.08 | 0.6413 | -0.08 | 0.2434 | -0.08 | 0.9567 |
| D2 | -0.01 | 0.9831 | -0.01 | 0.5261 | -0.01 | 0.6028 |
| DAT | 0.11 | 0.5023 | 0.11 | 0.9019 | 0.11 | 0.7061 |
| GABAa | 0.25 | 0.0503 | **0.25** | **0.0009** | **0.25** | **0.0005** |
| H3 | -0.03 | 0.9164 | -0.03 | 0.9168 | -0.03 | 0.5706 |
| M1 | 0.12 | 0.6413 | 0.12 | 0.5261 | 0.12 | 0.281 |
| Mu | -0.23 | 0.5023 | -0.23 | 0.5261 | -0.23 | 0.5968 |
| NET | **0.57** | **0.0019** | **0.57** | **0.0009** | **0.57** | **0.0005** |
| NMDA | 0.28 | 0.1194 | 0.28 | 0.2434 | 0.28 | 0.1086 |
| SERT | 0.19 | 0.1194 | 0.19 | 0.9168 | 0.19 | 0.5968 |
| VAChT | 0.22 | 0.0849 | 0.22 | 0.2434 | 0.22 | 0.5968 |
| mGluR5 | 0.25 | 0.2344 | **0.25** | **0.0047** | **0.25** | **0.0005** |

Bold numbers highlight significant correlations. Rho: Spearman correlation coefficient.

**Supplementary Table 11**: Co-localization strength of **fALFF** with neurotransmitter systems and the effects of age and sex – *before atrophy correction*.

| | Co-localization strength* | | | Linear aging effects** | | | Sex differences** | | |
|---|---|---|---|---|---|---|---|---|---|
| Neurotransmitter system | Median rho | IQR rho | Mean Fisher's z(rho) | Pearson r | Slope | $P_{BH}$ | T | Cohen's d | $P_{BH}$ |
| 5-HT1a | -0.21 | 0.09 | -0.2 | 0.11 | 0.0011 | <0.001 | 11.26 | 0.14 | <0.001 |
| 5-HT1b | 0.43 | 0.07 | 0.46 | -0.02 | -0.0002 | 0.1079 | -3.57 | -0.04 | 0.0069 |
| 5-HT2a | 0.4 | 0.09 | 0.42 | 0.08 | 0.0008 | <0.001 | 8.18 | 0.1 | <0.001 |
| 5-HT4 | -0.29 | 0.09 | -0.29 | 0.08 | 0.0008 | <0.001 | 12.37 | 0.15 | <0.001 |
| 5-HT6 | 0.27 | 0.08 | 0.27 | -0.06 | -0.0005 | <0.001 | -3.74 | -0.05 | 0.0035 |
| A4B2 | 0.1 | 0.1 | 0.11 | 0.02 | 0.0003 | 0.0027 | 5.91 | 0.07 | <0.001 |
| CB1 | -0.14 | 0.14 | -0.13 | 0.09 | 0.0013 | <0.001 | 13.24 | 0.17 | <0.001 |
| D1 | -0.42 | 0.08 | -0.44 | 0.04 | 0.0004 | <0.001 | 9.16 | 0.11 | <0.001 |
| D2 | -0.47 | 0.09 | -0.51 | 0.04 | 0.0005 | <0.001 | -0.81 | -0.01 | 1 |
| DAT | -0.68 | 0.06 | -0.82 | -0.07 | -0.0009 | <0.001 | -9.61 | -0.12 | <0.001 |
| GABAa | 0.4 | 0.07 | 0.43 | -0.06 | -0.0005 | <0.001 | -7.43 | -0.09 | <0.001 |
| H3 | -0.28 | 0.11 | -0.27 | 0.01 | 0.0002 | 0.3196 | 1.96 | 0.02 | 0.9435 |
| M1 | 0.33 | 0.07 | 0.34 | -0.02 | -0.0001 | 0.2195 | 1.34 | 0.02 | 1 |
| μ | -0.5 | 0.13 | -0.53 | 0.11 | 0.0019 | <0.001 | 13.58 | 0.17 | <0.001 |
| NET | 0.51 | 0.12 | 0.57 | -0.13 | -0.0019 | <0.001 | -29.86 | -0.37 | <0.001 |
| NMDA | -0.13 | 0.07 | -0.13 | -0.17 | -0.0013 | <0.001 | -19.67 | -0.25 | <0.001 |
| SERT | -0.5 | 0.07 | -0.54 | -0.17 | -0.0015 | <0.001 | -25.48 | -0.32 | <0.001 |
| VAChT | -0.56 | 0.11 | -0.63 | -0.05 | -0.0008 | <0.001 | -14.58 | -0.18 | <0.001 |
| mGluR5 | 0.3 | 0.1 | 0.32 | <0.01 | 0.0001 | 1 | -1.27 | -0.02 | 1 |

* The distribution of all Fisher's z-transformed Spearman correlation coefficients (Rho) regarding a specific neurotransmitter system were all significantly (one-sample t-test, $P_{BH}$<0.0001) different from a null-distribution.

**Aging effects and sex differences based on linear regression and comparison (t-test, alpha=0.05), respectively, between men and women in individual Fisher's z-transformed Spearman correlation coefficients. Not significant aging effects and sex differences are highlighted in red.

**Supplementary Table 12:** Co-localization strength of **LCOR** with neurotransmitter systems and the effects of age and sex – *before atrophy correction*.

| | Co-localization strength* | | | Linear aging effects** | | | Sex differences** | | |
|---|---|---|---|---|---|---|---|---|---|
| Neurotransmitter system | Median rho | IQR rho | Mean Fisher's z(rho) | Pearson r | Slope | $P_{BH}$ | T | Cohen's d | $P_{BH}$ |
| 5-HT1a | -0.3 | 0.1 | -0.31 | -0.01 | -0.0001 | 1 | -1.57 | -0.02 | 1 |
| 5-HT1b | 0.32 | 0.11 | 0.33 | 0.12 | 0.0015 | <0.001 | 5.78 | 0.07 | <0.001 |
| 5-HT2a | 0.2 | 0.11 | 0.2 | 0.13 | 0.0015 | <0.001 | 9.1 | 0.11 | <0.001 |
| 5-HT4 | -0.4 | 0.1 | -0.42 | 0.01 | 0.0001 | 1 | 5.99 | 0.07 | <0.001 |
| 5-HT6 | 0.29 | 0.12 | 0.31 | -0.06 | -0.0008 | <0.001 | -3.68 | -0.05 | 0.0045 |
| A4B2 | 0.03 | 0.1 | 0.04 | 0.06 | 0.0007 | <0.001 | 2.74 | 0.03 | 0.1164 |
| CB1 | -0.25 | 0.13 | -0.25 | 0.07 | 0.0009 | <0.001 | 7.17 | 0.09 | <0.001 |
| D1 | -0.26 | 0.14 | -0.25 | -0.08 | -0.0013 | <0.001 | 0.83 | 0.01 | 1 |
| D2 | -0.43 | 0.13 | -0.46 | -0.11 | -0.0018 | <0.001 | -6.78 | -0.08 | <0.001 |
| DAT | -0.39 | 0.16 | -0.41 | -0.2 | -0.0037 | <0.001 | -10.78 | -0.14 | <0.001 |
| GABAa | 0.49 | 0.09 | 0.54 | -0.06 | -0.0007 | <0.001 | -1.72 | -0.02 | 1 |
| H3 | -0.21 | 0.16 | -0.2 | -0.06 | -0.001 | <0.001 | -2.68 | -0.03 | 0.1421 |
| M1 | 0.22 | 0.11 | 0.23 | 0.05 | 0.0006 | <0.001 | 7.7 | 0.1 | <0.001 |
| μ | -0.51 | 0.13 | -0.54 | 0.02 | 0.0004 | 0.0071 | 7.9 | 0.1 | <0.001 |
| NET | 0.59 | 0.13 | 0.67 | -0.11 | -0.002 | <0.001 | -23.75 | -0.3 | <0.001 |
| NMDA | 0.15 | 0.14 | 0.15 | -0.2 | -0.0029 | <0.001 | -12.5 | -0.16 | <0.001 |
| SERT | -0.29 | 0.15 | -0.3 | -0.21 | -0.0033 | <0.001 | -15.87 | -0.2 | <0.001 |
| VAChT | -0.33 | 0.19 | -0.34 | -0.17 | -0.0035 | <0.001 | -14.03 | -0.18 | <0.001 |
| mGluR5 | 0.38 | 0.14 | 0.41 | -0.04 | -0.0008 | <0.001 | -3.3 | -0.04 | 0.0183 |

* The distribution of all Fisher's z-transformed Spearman correlation coefficients (Rho) regarding a specific neurotransmitter system were all significantly (one-sample t-test, $P_{BH}$<0.0001) different from a null-distribution. IQR: Interquartile range of Spearman correlation coefficients.

**Aging effects and sex differences based on linear regression and comparison (t-test, alpha=0.05), respectively, between men and women in individual Fisher's z-transformed Spearman correlation coefficients. Not significant aging effects and sex differences are highlighted in red.

**Supplementary Table 13:** Co-localization strength of **GCOR** with neurotransmitter systems and the effects of age and sex – *before atrophy correction*.

| | Co-localization strength* | | | Linear aging effects** | | | Sex differences** | | |
|---|---|---|---|---|---|---|---|---|---|
| Neurotransmitter system | Median rho | IQR rho | Mean Fisher's z(rho) | Pearson r | Slope | $P_{BH}$ | T | Cohen's d | $P_{BH}$ |
| 5-HT1a | -0.24 | 0.21 | -0.24 | -0.1 | -0.0023 | <0.001 | -9.21 | -0.12 | <0.001 |
| 5-HT1b | 0.14 | 0.16 | 0.14 | 0.09 | 0.0015 | <0.001 | -11.82 | -0.15 | <0.001 |
| 5-HT2a | 0.1 | 0.17 | 0.1 | 0.02 | 0.0003 | 0.0405 | -1.57 | -0.02 | 1 |
| 5-HT4 | -0.28 | 0.16 | -0.28 | -0.01 | -0.0002 | 1 | 1.87 | 0.02 | 1 |
| 5-HT6 | 0.19 | 0.19 | 0.19 | -0.05 | -0.001 | <0.001 | -15.69 | -0.2 | <0.001 |
| A4B2 | 0.08 | 0.2 | 0.08 | <0.01 | -0.0001 | 1 | -4.11 | -0.05 | 0.0007 |
| CB1 | -0.26 | 0.22 | -0.25 | -0.04 | -0.0009 | <0.001 | -9.82 | -0.12 | <0.001 |
| D1 | -0.2 | 0.22 | -0.2 | -0.01 | -0.0002 | 1 | -7.96 | -0.1 | <0.001 |
| D2 | -0.29 | 0.21 | -0.29 | -0.07 | -0.0016 | <0.001 | -9.41 | -0.12 | <0.001 |
| DAT | -0.23 | 0.22 | -0.23 | -0.07 | -0.0015 | <0.001 | -8.61 | -0.11 | <0.001 |
| GABAa | 0.31 | 0.16 | 0.32 | -0.04 | -0.0007 | <0.001 | -14.42 | -0.18 | <0.001 |
| H3 | -0.17 | 0.28 | -0.17 | -0.04 | -0.0012 | <0.001 | -10.8 | -0.14 | <0.001 |
| M1 | 0.09 | 0.14 | 0.09 | 0.06 | 0.0009 | <0.001 | -7.28 | -0.09 | <0.001 |
| μ | -0.36 | 0.25 | -0.37 | -0.01 | -0.0003 | 1 | -2.01 | -0.03 | 0.8398 |
| NET | 0.5 | 0.23 | 0.53 | -0.11 | -0.0032 | <0.001 | -19.34 | -0.24 | <0.001 |
| NMDA | 0.16 | 0.19 | 0.16 | -0.07 | -0.0013 | <0.001 | -11.4 | -0.14 | <0.001 |
| SERT | -0.17 | 0.18 | -0.18 | -0.08 | -0.0014 | <0.001 | -8.72 | -0.11 | <0.001 |
| VAChT | -0.15 | 0.28 | -0.15 | -0.12 | -0.0032 | <0.001 | -11.21 | -0.14 | <0.001 |
| mGluR5 | 0.27 | 0.27 | 0.28 | -0.09 | -0.0025 | <0.001 | -18.49 | -0.23 | <0.001 |

* The distribution of all Fisher's z-transformed Spearman correlation coefficients (Rho) regarding a specific neurotransmitter system were all significantly (one-sample t-test, $P_{BH}<0.0001$) different from a null-distribution. IQR: Interquartile range of Spearman correlation coefficients.

**Aging effects and sex differences based on linear regression and comparison (t-test, alpha=0.05), respectively, between men and women in individual Fisher's z-transformed Spearman correlation coefficients. Not significant aging effects and sex differences are highlighted in red.

**Supplementary Table 14:** Co-localization strength of **fALFF** with neurotransmitter systems and the effects of age and sex – *after atrophy correction*.

| | Co-localization strength* | | | Linear aging effects** | | | Sex differences** | | |
|---|---|---|---|---|---|---|---|---|---|
| Neurotransmitter system | Median rho | IQR rho | Mean Fisher's z(rho) | Pearson r | Slope | $P_{BH}$ | T | Cohen's d | $P_{BH}$ |
| 5-HT1a | -0.21 | 0.09 | -0.2 | 0.1 | 0.001 | <0.0001 | 11.86 | 0.15 | <0.0001 |
| 5-HT1b | 0.43 | 0.07 | 0.46 | -0.02 | -0.0002 | 0.0992 | -3 | -0.04 | 0.0512 |
| 5-HT2a | 0.4 | 0.09 | 0.42 | 0.08 | 0.0007 | <0.0001 | 8.91 | 0.11 | <0.0001 |
| 5-HT4 | -0.29 | 0.09 | -0.29 | 0.07 | 0.0007 | <0.0001 | 13.28 | 0.17 | <0.0001 |
| 5-HT6 | 0.26 | 0.08 | 0.27 | -0.04 | -0.0003 | <0.0001 | -4.02 | -0.05 | 0.0011 |
| A4B2 | 0.1 | 0.1 | 0.11 | 0.02 | 0.0002 | 0.0699 | 6.24 | 0.08 | <0.0001 |
| CB1 | -0.14 | 0.14 | -0.13 | 0.08 | 0.0011 | <0.0001 | 13.85 | 0.17 | <0.0001 |
| D1 | -0.42 | 0.08 | -0.44 | 0.06 | 0.0006 | <0.0001 | 8.53 | 0.11 | <0.0001 |
| D2 | -0.47 | 0.08 | -0.51 | 0.05 | 0.0005 | <0.0001 | -0.49 | -0.01 | 1 |
| DAT | -0.68 | 0.06 | -0.82 | -0.04 | -0.0005 | <0.0001 | -10.81 | -0.14 | <0.0001 |
| GABAa | 0.4 | 0.07 | 0.43 | -0.03 | -0.0003 | <0.0001 | -8.64 | -0.11 | <0.0001 |
| H3 | -0.28 | 0.11 | -0.27 | 0.02 | 0.0002 | 0.0665 | 2.15 | 0.03 | 0.6061 |
| M1 | 0.33 | 0.07 | 0.34 | 0 | 0 | 1 | 1.61 | 0.02 | 1 |
| μ | -0.5 | 0.13 | -0.53 | 0.09 | 0.0016 | <0.0001 | 14.11 | 0.18 | <0.0001 |
| NET | 0.51 | 0.12 | 0.57 | -0.11 | -0.0016 | <0.0001 | -30.12 | -0.38 | <0.0001 |
| NMDA | -0.13 | 0.07 | -0.13 | -0.13 | -0.001 | <0.0001 | -20.97 | -0.26 | <0.0001 |
| SERT | -0.5 | 0.06 | -0.54 | -0.14 | -0.0012 | <0.0001 | -26.31 | -0.33 | <0.0001 |
| VAChT | -0.56 | 0.1 | -0.63 | -0.04 | -0.0006 | <0.0001 | -14.69 | -0.18 | <0.0001 |
| mGluR5 | 0.3 | 0.1 | 0.32 | 0.02 | 0.0002 | 0.0218 | -1.76 | -0.02 | 1 |

* The distribution of all Fisher's z-transformed Spearman correlation coefficients (Rho) regarding a specific neurotransmitter system were all significantly (one-sample t-test, $P_{BH}$<0.0001) different from a null-distribution. IQR: Interquartile range of Spearman correlation coefficients.

**Aging effects and sex differences based on linear regression and comparison (t-test, alpha=0.05), respectively, between men and women in individual Fisher's z-transformed Spearman correlation coefficients. Not significant aging effects and sex differences are highlighted in red.

**Supplementary Table 15:** Co-localization strength of **LCOR** with neurotransmitter systems and the effects of age and sex – *after atrophy correction*.

| | Co-localization strength* | | | Linear aging effects** | | | Sex differences** | | |
|---|---|---|---|---|---|---|---|---|---|
| Neurotransmitter system | Median rho | IQR rho | Mean Fisher's z(rho) | Pearson r | Slope | $P_{BH}$ | T | Cohen's d | $P_{BH}$ |
| 5-HT1a | -0.3 | 0.1 | -0.31 | 0 | -0.0001 | 1 | -1.91 | -0.02 | 1 |
| 5-HT1b | 0.32 | 0.11 | 0.33 | 0.11 | 0.0013 | <0.0001 | 6.45 | 0.08 | <0.0001 |
| 5-HT2a | 0.2 | 0.11 | 0.2 | 0.12 | 0.0014 | <0.0001 | 9.47 | 0.12 | <0.0001 |
| 5-HT4 | -0.4 | 0.1 | -0.42 | 0.01 | 0.0002 | 0.4093 | 6.14 | 0.08 | <0.0001 |
| 5-HT6 | 0.29 | 0.12 | 0.31 | -0.05 | -0.0006 | <0.0001 | -3.92 | -0.05 | 0.0017 |
| A4B2 | 0.04 | 0.1 | 0.04 | 0.06 | 0.0006 | <0.0001 | 3.15 | 0.04 | 0.0312 |
| CB1 | -0.25 | 0.13 | -0.25 | 0.06 | 0.0009 | <0.0001 | 7.21 | 0.09 | <0.0001 |
| D1 | -0.26 | 0.14 | -0.25 | -0.07 | -0.001 | <0.0001 | 0.2 | 0 | 1 |
| D2 | -0.43 | 0.13 | -0.46 | -0.09 | -0.0015 | <0.0001 | -7.34 | -0.09 | <0.0001 |
| DAT | -0.39 | 0.16 | -0.41 | -0.17 | -0.0032 | <0.0001 | -11.62 | -0.15 | <0.0001 |
| GABAa | 0.49 | 0.09 | 0.54 | -0.04 | -0.0005 | <0.0001 | -2.53 | -0.03 | 0.2167 |
| H3 | -0.21 | 0.16 | -0.2 | -0.05 | -0.0008 | <0.0001 | -2.91 | -0.04 | 0.0689 |
| M1 | 0.22 | 0.11 | 0.23 | 0.06 | 0.0007 | <0.0001 | 7.82 | 0.1 | <0.0001 |
| μ | -0.51 | 0.13 | -0.54 | 0.02 | 0.0003 | 0.041 | 7.89 | 0.1 | <0.0001 |
| NET | 0.59 | 0.12 | 0.67 | -0.1 | -0.0017 | <0.0001 | -23.99 | -0.3 | <0.0001 |
| NMDA | 0.15 | 0.14 | 0.15 | -0.18 | -0.0026 | <0.0001 | -13.3 | -0.17 | <0.0001 |
| SERT | -0.29 | 0.15 | -0.3 | -0.19 | -0.003 | <0.0001 | -16.56 | -0.21 | <0.0001 |
| VAChT | -0.33 | 0.19 | -0.34 | -0.15 | -0.0032 | <0.0001 | -14.51 | -0.18 | <0.0001 |
| mGluR5 | 0.38 | 0.14 | 0.41 | -0.03 | -0.0005 | 0.0001 | -3.83 | -0.05 | 0.0025 |

* The distribution of all Fisher's z-transformed Spearman correlation coefficients (Rho) regarding a specific neurotransmitter system were all significantly (one-sample t-test, $P_{BH}$<0.0001) different from a null-distribution. IQR: Interquartile range of Spearman correlation coefficients.

**Aging effects and sex differences based on linear regression and comparison (t-test, alpha=0.05), respectively, between men and women in individual Fisher's z-transformed Spearman correlation coefficients. Not significant aging effects and sex differences are highlighted in red.

**Supplementary Table 16:** Co-localization strength of **GCOR** with neurotransmitter systems and the effects of age and sex – *after atrophy correction*.

| | Co-localization strength* | | | Linear aging effects** | | | Sex differences** | | |
|---|---|---|---|---|---|---|---|---|---|
| Neurotransmitter system | Median rho | IQR rho | Mean Fisher's z(rho) | Pearson r | Slope | $P_{BH}$ | T | Cohen's d | $P_{BH}$ |
| 5-HT1a | -0.24 | 0.21 | -0.24 | -0.1 | -0.0021 | <0.0001 | -9.63 | -0.12 | <0.0001 |
| 5-HT1b | 0.14 | 0.16 | 0.14 | 0.08 | 0.0013 | <0.0001 | -11.41 | -0.14 | <0.0001 |
| 5-HT2a | 0.1 | 0.17 | 0.1 | 0.02 | 0.0003 | 0.1381 | -1.41 | -0.02 | 1 |
| 5-HT4 | -0.28 | 0.16 | -0.28 | -0.01 | -0.0002 | 0.8886 | 2 | 0.03 | 0.8675 |
| 5-HT6 | 0.19 | 0.19 | 0.19 | -0.04 | -0.0007 | <0.0001 | -16.16 | -0.2 | <0.0001 |
| A4B2 | 0.08 | 0.2 | 0.08 | -0.01 | -0.0001 | 1 | -3.85 | -0.05 | 0.0023 |
| CB1 | -0.26 | 0.22 | -0.25 | -0.04 | -0.0009 | <0.0001 | -9.9 | -0.12 | <0.0001 |
| D1 | -0.2 | 0.22 | -0.2 | 0 | 0.0001 | 1 | -8.6 | -0.11 | <0.0001 |
| D2 | -0.29 | 0.21 | -0.3 | -0.06 | -0.0014 | <0.0001 | -9.96 | -0.12 | <0.0001 |
| DAT | -0.23 | 0.22 | -0.23 | -0.05 | -0.0011 | <0.0001 | -9.43 | -0.12 | <0.0001 |
| GABAa | 0.31 | 0.16 | 0.32 | -0.02 | -0.0004 | 0.02 | -15.39 | -0.19 | <0.0001 |
| H3 | -0.17 | 0.28 | -0.17 | -0.04 | -0.001 | <0.0001 | -11.03 | -0.14 | <0.0001 |
| M1 | 0.09 | 0.14 | 0.09 | 0.07 | 0.0009 | <0.0001 | -7.33 | -0.09 | <0.0001 |
| μ | -0.37 | 0.25 | -0.37 | -0.01 | -0.0004 | 0.5992 | -2 | -0.03 | 0.8633 |
| NET | 0.5 | 0.23 | 0.53 | -0.1 | -0.0028 | <0.0001 | -19.76 | -0.25 | <0.0001 |
| NMDA | 0.16 | 0.19 | 0.16 | -0.05 | -0.0008 | <0.0001 | -12.25 | -0.15 | <0.0001 |
| SERT | -0.17 | 0.18 | -0.18 | -0.06 | -0.0011 | <0.0001 | -9.47 | -0.12 | <0.0001 |
| VAChT | -0.15 | 0.28 | -0.15 | -0.1 | -0.0028 | <0.0001 | -11.69 | -0.15 | <0.0001 |
| mGluR5 | 0.27 | 0.27 | 0.28 | -0.08 | -0.002 | <0.0001 | -19.23 | -0.24 | <0.0001 |

* The distribution of all Fisher's z-transformed Spearman correlation coefficients (Rho) regarding a specific neurotransmitter system were all significantly (one-sample t-test, $P_{BH}$<0.0001) different from a null-distribution. IQR: Interquartile range of Spearman correlation coefficients.

**Aging effects and sex differences based on linear regression and comparison (t-test, alpha=0.05), respectively, between men and women in individual Fisher's z-transformed Spearman correlation coefficients. Not significant aging effects and sex differences are highlighted in red.

**Supplementary Table 17**: Statistical key figures of the White- and Goldfeld-Quandt-test for heteroskedasticity in the co-localizations (Fisher's z-transformed Spearman correlation coefficients) – *before atrophy correction*.

| | White-test | | | | | | Goldfeld-Quandt-test | | | | | |
|---|---|---|---|---|---|---|---|---|---|---|---|---|
| | fALFF | | LCOR | | GCOR | | fALFF | | LCOR | | GCOR | |
| Neurotrans-mitter system | F | pFDR | F | pFDR | F | pFDR | F | pFDR | F | pFDR | F | pFDR |
| 5-HT1a | **13.43** | **<0.0001** | 2.78 | 0.07 | **23.07** | **<0.0001** | **1.11** | **6.5E-07** | - | - | 0.88 | 1 |
| 5-HT1b | **4.01** | **0.025** | 2.79 | 0.07 | **4.81** | **0.015** | 0.95 | 0.99 | - | - | **1.06** | **0.013** |
| 5-HT2a | **4.66** | **0.015** | **7.46** | **0.0008** | 0.49 | 0.68 | **1.06** | **0.025** | **1.05** | **0.022** | - | - |
| 5-HT4 | **14.68** | **<0.0001** | **28.91** | **<0.0001** | 1.16 | 0.40 | **1.10** | **<0.0001** | **1.15** | **<0.0001** | - | - |
| 5-HT6 | 1.03 | 0.40 | **7.88** | **0.0006** | **4.76** | **0.015** | - | - | **1.08** | **0.0003** | **1.05** | **0.021** |
| A4B2 | 1.13 | 0.38 | 2.64 | 0.07 | **22.04** | **<0.0001** | - | - | - | - | 0.86 | 1 |
| CB1 | **4.63** | **0.015** | **7.9** | **0.0006** | **6.66** | **0.003** | **1.04** | **0.027** | **1.08** | **0.0003** | 0.92 | 1 |
| D1 | **16.01** | **<0.0001** | **4.52** | **0.014** | 1.79 | 0.24 | **1.13** | **<0.0001** | **1.06** | **0.003** | - | - |
| D2 | **19.55** | **<0.0001** | **16.12** | **<0.0001** | 0.22 | 0.80 | **1.14** | **<0.0001** | **1.13** | **<0.0001** | - | - |
| DAT | **14.98** | **<0.0001** | **13.47** | **<0.0001** | 1.27 | 0.38 | **1.14** | **<0.0001** | **1.09** | **<0.0001** | - | - |
| GABAa | **6.19** | **0.004** | **11.34** | **<0.0001** | 1.95 | 0.22 | 1.04 | 0.06 | **1.10** | **<0.0001** | - | - |
| H3 | **13.53** | **<0.0001** | **16.36** | **<0.0001** | 0.44 | 0.68 | **1.10** | **<0.0001** | **1.13** | **<0.0001** | - | - |
| M1 | 0.45 | 0.67 | **4.78** | **0.011** | **10.46** | **0.0001** | - | - | 1.03 | 0.07 | **1.10** | **<0.0001** |
| µ-opioid | **4.1** | **0.02** | 3.13 | 0.052 | **29.49** | **<0.0001** | 1.03 | 0.13 | - | - | 0.86 | 1 |
| NET | **13.24** | **<0.0001** | **50.27** | **<0.0001** | **9.07** | **0.0004** | **1.09** | **<0.0001** | **1.22** | **<0.0001** | **1.09** | **0.0004** |
| NMDA | 2.42 | 0.11 | **17.32** | **<0.0001** | **7.51** | **0.002** | - | - | **1.10** | **<0.0001** | **1.06** | **0.013** |
| SERT | 0.13 | 0.88 | **14.51** | **<0.0001** | **4.97** | **0.015** | - | - | **1.07** | **0.002** | **1.05** | **0.020** |
| VAChT | **37.62** | **<0.0001** | **18.68** | **<0.0001** | 0.59 | 0.66 | **1.18** | **<0.0001** | **1.11** | **<0.0001** | - | - |
| mGluR5 | **10.91** | **<0.0001** | **25.01** | **<0.0001** | **7.29** | **0.002** | **1.10** | **<0.0001** | **1.18** | **<0.0001** | **1.05** | **0.020** |

Bold numbers highlight significant results. F: F-statitstic.

**Supplementary Table 18**: Statistical key figures of the White- and Goldfeld-Quandt-test for heteroskedasticity in the co-localizations (Fisher's z-transformed Spearman correlation coefficients) – *after atrophy correction*.

| | White-test | | | | | | Goldfeld-Quandt-test | | | | | |
|---|---|---|---|---|---|---|---|---|---|---|---|---|
| | fALFF | | LCOR | | GCOR | | fALFF | | LCOR | | GCOR | |
| Neurotrans-mitter system | F | pFDR | F | pFDR | F | pFDR | F | pFDR | F | pFDR | F | pFDR |
| 5-HT1a | **11.59** | **<0.0001** | 2.16 | 0.12 | **26.28** | **<0.0001** | **1.11** | **<0.0001** | - | - | 0.87 | 1 |
| 5-HT1b | **3.77** | **0.034** | 3.12 | 0.053 | 3.6 | 0.058 | 0.95 | 0.99 | - | - | - | - |

| | | | | | | | | | | | | |
|---|---|---|---|---|---|---|---|---|---|---|---|---|
| 5-HT2a | **4.9** | **0.013** | **6.59** | **0.002** | 0.18 | 0.83 | **1.07** | **0.001** | 1.04 | 0.051 | - | - |
| 5-HT4 | **12.72** | **<0.0001** | **23.09** | **<0.0001** | 0.49 | 0.74 | **1.09** | **<0.0001** | **1.14** | **<0.0001** | - | - |
| 5-HT6 | 0.77 | 0.52 | **6.29** | **0.003** | 2.67 | 0.12 | - | - | **1.07** | **0.001** | - | - |
| A4B2 | 1.05 | 0.42 | 1.61 | 0.20 | **25.34** | **<0.0001** | - | - | - | - | 0.85 | 1 |
| CB1 | **3.95** | **0.030** | **6.89** | **0.002** | **7.28** | **0.003** | **1.04** | **0.042** | **1.07** | **0.001** | 0.92 | 1 |
| D1 | **13.64** | **<0.0001** | **3.31** | **0.046** | 0.83 | 0.63 | **1.12** | **<0.0001** | **1.05** | **0.012** | - | - |
| D2 | **17.44** | **<0.0001** | **12.64** | **<0.0001** | 0.78 | 0.63 | **1.13** | **<0.0001** | **1.11** | **<0.0001** | - | - |
| DAT | **15.27** | **<0.0001** | **9.5** | **0.0002** | 0.41 | 0.75 | **1.14** | **<0.0001** | **1.07** | **0.001** | - | - |
| GABAa | **5.54** | **0.007** | **8.01** | **0.0006** | 0.33 | 0.76 | 1.03 | 0.07 | **1.08** | **0.0002** | - | - |
| H3 | **11.57** | **<0.0001** | **14.99** | **<0.0001** | 0.4 | 0.75 | **1.09** | **<0.0001** | **1.12** | **<0.0001** | - | - |
| M1 | 0.34 | 0.75 | **3.64** | **0.036** | **7.87** | **0.002** | - | - | 1.02 | 0.13 | **1.09** | **0.0003** |
| μ-opioid | **3.46** | **0.042** | 3.04 | 0.054 | **29.43** | **<0.0001** | 1.02 | 0.18 | - | - | 0.86 | 1 |
| NET | **10.95** | **<0.0001** | **37.38** | **<0.0001** | **5.24** | **0.017** | **1.08** | **0.0002** | **1.19** | **<0.0001** | **1.06** | **0.008** |
| NMDA | 2.62 | 0.09 | **12.42** | **<0.0001** | **4** | **0.044** | - | - | **1.08** | **0.0004** | 1.04 | 0.13 |
| SERT | 0.06 | 0.94 | **10.68** | **<0.0001** | 2.92 | 0.10 | - | - | **1.05** | **0.018** | - | - |
| VAChT | **35.68** | **<0.0001** | **15.35** | **<0.0001** | 0.94 | 0.62 | **1.18** | **<0.0001** | **1.09** | **<0.0001** | - | - |
| mGluR5 | **11.86** | **<0.0001** | **22.69** | **<0.0001** | **4.03** | **0.044** | **1.11** | **<0.0001** | **1.17** | **<0.0001** | 1.03 | 0.13 |

Bold numbers highlight significant results. F: F-statitstic.

**Supplementary Table 19**: Deviation scores of subjects with manifest PD compared to normative models – *before atrophy correction*.

| | fALFF | | LCOR | | GCOR | |
|---|---|---|---|---|---|---|
| Neurotransmitter system | Median Z | $P_{FDR}$ | Median Z | $P_{FDR}$ | Median Z | $P_{FDR}$ |
| 5-HT1a | -0.13 | 0.4435 | 0.07 | 0.797 | 0.01 | 0.775 |
| 5-HT1b | **-0.4** | **0.0043** | **-0.29** | **0.0478** | **-0.29** | **0.0163** |
| 5-HT2a | -0.26 | 0.1052 | 0 | 0.7309 | -0.01 | 0.5572 |
| 5-HT4 | -0.16 | 0.9167 | -0.08 | 0.2989 | -0.22 | 0.2844 |
| 5-HT6 | **-0.47** | **0.0043** | **-0.39** | **0.0056** | **-0.51** | **0.0001** |
| A4B2 | -0.07 | 0.9167 | 0.13 | 0.2677 | -0.19 | 0.3655 |
| CB1 | -0.2 | 0.4435 | -0.28 | 0.1216 | -0.18 | 0.139 |
| D1 | -0.21 | 0.6035 | **-0.29** | **0.0056** | **-0.43** | **0.0019** |
| D2 | 0.03 | 0.3857 | **-0.56** | **0.0021** | **-0.45** | **0.0005** |
| DAT | -0.39 | 0.4435 | **-0.45** | **0.0212** | **-0.37** | **0.0118** |
| GABAa | **-0.52** | **0.0089** | **-0.5** | **<0.0001** | **-0.64** | **0.0019** |
| H3 | -0.25 | 0.0805 | **-0.3** | **0.0245** | **-0.29** | **0.0306** |
| M1 | **-0.56** | **0.0031** | -0.17 | 0.1156 | **-0.52** | **0.0005** |
| Mu | -0.26 | 0.3275 | -0.27 | 0.2523 | 0 | 0.5121 |
| NET | -0.18 | 0.2474 | **-0.46** | **0.0011** | **-0.31** | **0.0306** |
| NMDA | -0.09 | 0.9167 | **-0.38** | **0.0021** | **-0.5** | **0.0005** |
| SERT | -0.25 | 0.9167 | -0.41 | 0.0772 | **-0.37** | **0.0337** |
| VAChT | -0.23 | 0.3319 | **-0.26** | **0.0478** | -0.22 | 0.1279 |
| mGluR5 | **-0.43** | **0.0061** | **-0.43** | **0.0023** | **-0.33** | **0.0005** |

Bold numbers highlight significant correlations. Median Z: Median deviation score of all subjects with manifest Parkinson's disease.

**Supplementary Table 20**: Deviation scores of subjects with manifest PD compared to normative models – *after atrophy correction*.

| | fALFF | | LCOR | | GCOR | |
|---|---|---|---|---|---|---|
| Neurotransmitter system | Median Z | $P_{FDR}$ | Median Z | $P_{FDR}$ | Median Z | $P_{FDR}$ |
| 5-HT1a | -0.18 | 0.3879 | 0.05 | 0.8283 | 0.01 | 0.803 |
| 5-HT1b | **-0.38** | **0.0052** | -0.31 | 0.0512 | **-0.38** | **0.0137** |
| 5-HT2a | -0.21 | 0.101 | -0.01 | 0.729 | -0.01 | 0.5539 |
| 5-HT4 | -0.2 | 0.9584 | -0.11 | 0.299 | -0.22 | 0.3222 |
| 5-HT6 | **-0.45** | **0.0058** | **-0.41** | **0.0066** | **-0.43** | **0.0001** |
| A4B2 | -0.09 | 0.9584 | 0.13 | 0.2565 | -0.16 | 0.3484 |
| CB1 | -0.22 | 0.3805 | -0.29 | 0.1182 | -0.24 | 0.123 |
| D1 | -0.17 | 0.7212 | **-0.38** | **0.0066** | **-0.49** | **0.002** |
| D2 | 0 | 0.3805 | **-0.52** | **0.0023** | **-0.47** | **0.0007** |
| DAT | -0.31 | 0.4792 | **-0.47** | **0.0244** | **-0.26** | **0.014** |
| GABAa | **-0.58** | **0.0128** | **-0.48** | **0.0001** | **-0.6** | **0.002** |
| H3 | -0.29 | 0.0753 | **-0.31** | **0.0256** | **-0.3** | **0.0347** |
| M1 | **-0.59** | **0.0032** | -0.21 | 0.1203 | **-0.59** | **0.0006** |
| Mu | -0.26 | 0.3115 | -0.29 | 0.2242 | 0 | 0.4845 |
| NET | -0.13 | 0.3805 | **-0.48** | **0.0012** | **-0.29** | **0.0347** |
| NMDA | -0.09 | 0.962 | **-0.35** | **0.0023** | **-0.46** | **0.0006** |
| SERT | -0.25 | 0.9584 | -0.41 | 0.0813 | **-0.35** | **0.043** |
| VAChT | -0.19 | 0.3805 | -0.24 | 0.0512 | -0.22 | 0.1384 |
| mGluR5 | **-0.42** | **0.0058** | **-0.39** | **0.0027** | **-0.3** | **0.0007** |

Bold numbers highlight significant correlations. Median Z: Median deviation score of all subjects with manifest Parkinson's disease.

**Supplementary Table 21**: Correlation between deviation scores and reported disease duration in subjects with manifest PD – *before atrophy correction*.

| | fALFF | | LCOR | | GCOR | |
|---|---|---|---|---|---|---|
| Neurotransmitter system | Pearson r | $P_{FDR}$ | Pearson r | $P_{FDR}$ | Pearson r | $P_{FDR}$ |
| 5-HT1a | - | - | - | - | - | - |
| 5-HT1b | -0.16 | 0.3969 | -0.2 | 0.6141 | -0.13 | 0.9659 |
| 5-HT2a | - | - | - | - | - | - |
| 5-HT4 | - | - | - | - | - | - |
| 5-HT6 | -0.06 | 0.7641 | -0.11 | 0.7756 | -0.03 | 0. 9659 |
| A4B2 | - | - | - | - | - | - |
| CB1 | - | - | - | - | - | - |
| D1 | - | - | -0.16 | 0.6141 | -0.07 | 0. 9659 |
| D2 | - | - | -0.03 | 0.9121 | 0.02 | 0. 9659 |
| DAT | - | - | -0.01 | 0.9121 | -0.02 | 0. 9659 |
| GABAa | -0.16 | 0. 3969 | **-0.38** | **0.0317** | -0.11 | 0. 9659 |
| H3 | - | - | -0.02 | 0.9121 | 0.01 | 0. 9659 |
| M1 | -0.22 | 0. 3969 | - | - | -0.21 | 0. 9659 |
| Mu | - | - | - | - | - | - |
| NET | - | - | 0.13 | 0.7508 | 0.02 | 0. 9659 |
| NMDA | - | - | -0.02 | 0.9121 | -0.06 | 0. 9659 |
| SERT | - | - | - | - | 0.07 | 0. 9659 |
| VAChT | - | - | 0.18 | 0.6141 | - | - |
| mGluR5 | 0.04 | 0. 7641 | -0.09 | 0.8153 | 0.01 | 0. 9659 |

Bold numbers highlight significant correlations. Empty cells indicate pairs of functional measure and neurotransmitter system whose co-localization in PD was not significantly different from the norm.

**Supplementary Table 22**: Correlation between deviation scores and reported disease duration in subjects with manifest PD – *after atrophy correction*.

| | fALFF | | LCOR | | GCOR | |
|---|---|---|---|---|---|---|
| Neurotransmitter system | Pearson r | $P_{FDR}$ | Pearson r | $P_{FDR}$ | Pearson r | $P_{FDR}$ |
| 5-HT1a | - | - | - | - | - | - |
| 5-HT1b | -0.14 | 0.4986 | - | - | -0.12 | 0.9744 |
| 5-HT2a | - | - | - | - | - | - |
| 5-HT4 | - | - | - | - | - | - |
| 5-HT6 | -0.05 | 0.7495 | -0.09 | 0.9335 | -0.03 | 0. 9744 |
| A4B2 | - | - | - | - | - | - |
| CB1 | - | - | - | - | - | - |
| D1 | - | - | -0.16 | 0. 9335 | -0.07 | 0. 9744 |
| D2 | - | - | -0.02 | 0. 9335 | 0.03 | 0. 9744 |
| DAT | - | - | -0.02 | 0. 9335 | -0.02 | 0. 9744 |
| GABAa | -0.16 | 0. 4986 | **-0.38** | **0.0295** | -0.10 | 0. 9744 |
| H3 | - | - | -0.01 | 0. 9335 | 0.004 | 0. 9744 |
| M1 | -0.20 | 0. 4986 | - | - | -0.20 | 0. 9744 |
| Mu | - | - | - | - | - | - |
| NET | - | - | 0.13 | 0. 9335 | 0.02 | 0. 9744 |
| NMDA | - | - | -0.02 | 0. 9335 | -0.06 | 0. 9744 |
| SERT | - | - | - | - | 0.07 | 0. 9744 |
| VAChT | - | - | - | - | - | - |
| mGluR5 | 0.04 | 0. 7495 | -0.08 | 0. 9335 | 0.01 | 0. 9744 |

Bold numbers highlight significant correlations. Empty cells indicate pairs of functional measure and neurotransmitter system whose co-localization in PD was not significantly different from the norm.

**Supplementary Table 23:** Statistics on the correlation of effect sizes (Cohen's d) in fALFF, LCOR, and GCOR between PD and HCmatched and the regional contributions ($\Delta\rho^2$) – *before atrophy correction*. This analysis was performed only for the pairs of brain measure and neurotransmitter system in which PD deviated significantly from the norm.

| | fALFF | | LCOR | | GCOR | |
|---|---|---|---|---|---|---|
| Neurotransmitter map | Pearson r | $P_{FDR}$ | Pearson r | $P_{FDR}$ | Pearson r | $P_{FDR}$ |
| 5-HT1b | 0.02 | 0.80 | \|r\|<0.01 | 0.98 | -0.1 | 0.42 |
| 5-HT6 | 0.11 | 0.48 | 0.12 | 0.34 | 0.15 | 0.31 |
| D1 | - | - | 0.05 | 0.72 | \|r\|<0.01 | 0.97 |
| D2 | - | - | -0.06 | 0.68 | -0.05 | 0.69 |
| DAT | - | - | 0.16 | 0.28 | 0.14 | 0.31 |
| GABAa | 0.07 | 0.57 | 0.14 | 0.28 | 0.06 | 0.69 |
| H3 | - | - | -0.09 | 0.55 | -0.02 | 0.91 |
| M1 | 0.19 | 0.21 | - | - | -0.11 | 0.39 |
| NET | - | - | 0.14 | 0.28 | 0.11 | 0.39 |
| NMDA | - | - | **0.37** | **0.0003** | **0.44** | **<0.0001** |
| SERT | - | - | **-** | **-** | 0.20 | 0.11 |
| VAChT | - | - | -0.04 | 0.72 | - | - |
| mGluR5 | -0.10 | 0.48 | **-0.28** | **0.013** | **0.28** | **0.014** |

Bold numbers highlight significant correlation between effect sizes and regional contribution.

**Supplementary Table 24:** Statistics on the correlation of effect sizes (Cohen's d) in fALFF, LCOR, and GCOR between PD and HCmatched and the regional contributions ($\Delta\rho^2$) – *after atrophy correction*. This analysis was performed only for the pairs of brain measure and neurotransmitter system in which PD deviated significantly from the norm.

| | fALFF | | LCOR | | GCOR | |
|---|---|---|---|---|---|---|
| Neurotransmitter map | Pearson r | $P_{FDR}$ | Pearson r | $P_{FDR}$ | Pearson r | $P_{FDR}$ |
| 5-HT1b | 0.02 | 0.80 | - | - | -0.13 | 0.27 |
| 5-HT6 | 0.12 | 0.49 | 0.10 | 0.48 | **0.25** | **0.04** |
| D1 | - | - | 0.06 | 0.52 | 0.01 | 0.98 |
| D2 | - | - | -0.07 | 0.52 | -0.06 | 0.63 |
| DAT | - | - | 0.17 | 0.20 | 0.14 | 0.27 |
| GABAa | 0.04 | 0.79 | 0.06 | 0.52 | 0.12 | 0.31 |
| H3 | - | - | -0.08 | 0.52 | \|r\|<0.01 | 0.98 |
| M1 | **0.25** | **0.03** | - | - | -0.08 | 0.50 |
| NET | - | - | 0.12 | 0.44 | 0.14 | 0.27 |
| NMDA | - | - | **0.35** | **0.0009** | **0.42** | **<0.0001** |
| SERT | - | - | - | - | 0.20 | 0.10 |
| VAChT | - | - | - | - | - | - |
| mGluR5 | -0.09 | 0.48 | **-0.28** | **0.0092** | 0.22 | 0.06 |

Bold numbers highlight significant correlation between effect sizes and regional contribution.

**Supplementary Table 25:** Statistics on regional differences in fALFF, LCOR, and GCOR between PD and HCmatched assessed using the Mann-Whitney-U test – *before atrophy correction*.

| | fALFF | | | | LCOR | | | | GCOR | | | |
|---|---|---|---|---|---|---|---|---|---|---|---|---|
| | Left | | Right | | Left | | Right | | Left | | Right | |
| Region | U statistic | $P_{FDR}$ | U statistic | $P_{FDR}$ | U statistic | $P_{FDR}$ | U statistic | $P_{FDR}$ | U statistic | $P_{FDR}$ | U statistic | $P_{FDR}$ |
| Accumbens | 1.64 | 0.4662 | 1.11 | 0.6261 | **4.1** | **0.0013** | 1.85 | 0.1216 | **3.47** | **0.0087** | **2.56** | **0.0199** |
| Caudate | 2.18 | 0.3162 | 1.69 | 0.4662 | **3.12** | **0.0119** | 1.81 | 0.1302 | **3.95** | **0.0084** | **3.22** | **0.0101** |
| Putamen | 1.26 | 0.5564 | 0.55 | 0.8665 | **3.97** | **0.0017** | **2.66** | **0.0322** | **3.81** | **0.0084** | **3.11** | **0.0101** |
| Pallidum | 1.25 | 0.5564 | 0.09 | 0.996 | **3.4** | **0.0047** | 2.22 | 0.0622 | **2.63** | **0.0186** | **2.91** | **0.011** |
| Amygdala | 0.67 | 0.8421 | -0.13 | 0.996 | **2.46** | **0.0475** | 0.89 | 0.4933 | **2.6** | **0.019** | 2.03 | 0.0509 |
| Hippocampus | 0.72 | 0.8345 | -0.08 | 0.996 | **2.66** | **0.0322** | 0.93 | 0.4761 | **3.52** | **0.0087** | **2.13** | **0.0418** |
| Ant Cing Gyrus | 0.96 | 0.7272 | 0.76 | 0.8139 | 1.96 | 0.0981 | 2.04 | 0.084 | **2.38** | **0.0289** | **2.75** | **0.0152** |
| Mid Cing Gyrus | 0.7 | 0.8421 | 0.63 | 0.8421 | 1.23 | 0.3314 | 1.32 | 0.287 | **2.96** | **0.0106** | **3.03** | **0.0101** |
| Post Cing Gyrus | -0.16 | 0.996 | -0.03 | 0.996 | 0.85 | 0.5198 | 0.56 | 0.6829 | **2.21** | **0.0372** | **2.55** | **0.0199** |
| Parahip. Gyrus | -0.07 | 0.996 | 0.87 | 0.7707 | 1.58 | 0.1947 | 1.52 | 0.2104 | **2.42** | **0.0267** | **2.6** | **0.019** |
| Subcallosal Area | 0.21 | 0.996 | 0.24 | 0.996 | 2.23 | 0.0622 | -0.35 | 0.8179 | **2.07** | **0.0467** | 0.48 | 0.63 |
| Thalamus | 1.21 | 0.5573 | 0.66 | 0.8421 | 1.69 | 0.157 | 0.99 | 0.4458 | **2.76** | **0.015** | **2.3** | **0.0326** |
| Basal Forebrain | 0.15 | 0.996 | 1.26 | 0.5564 | **2.4** | **0.0496** | 2.38 | 0.0504 | **2.1** | **0.0445** | **2.87** | **0.0121** |
| Frontal Pole | 0.37 | 0.9503 | -0.19 | 0.996 | 0.51 | 0.7019 | 0.3 | 0.8449 | 0.78 | 0.4404 | 0.94 | 0.3558 |
| Gyrus Rectus | 0.43 | 0.9185 | -0.08 | 0.996 | 1.08 | 0.4018 | -0.07 | 0.9763 | 1.38 | 0.1815 | **2.58** | **0.0195** |
| Frontal Operculum | 1.81 | 0.4662 | 1.51 | 0.4874 | **3.03** | **0.0134** | 2.21 | 0.0622 | **3.12** | **0.0101** | **2.56** | **0.0199** |
| Ant Orbital | 1.39 | 0.5339 | 0.8 | 0.7994 | **2.62** | **0.0347** | 2.2 | 0.0622 | **2.93** | **0.0109** | **2.14** | **0.0409** |
| Lat Orbital | 1.57 | 0.4662 | 1.45 | 0.5282 | **3.04** | **0.0134** | **3.06** | **0.0133** | **3.06** | **0.0101** | **2.84** | **0.0123** |
| Med Orbital | 0.79 | 0.7994 | 0.93 | 0.7346 | 2.24 | 0.0622 | **2.39** | **0.0496** | 1.75 | 0.0917 | **2.91** | **0.011** |
| Post Orbital | 1.41 | 0.5339 | 1.24 | 0.5564 | **2.72** | **0.0297** | 2.35 | 0.0532 | **2.61** | **0.019** | **2.34** | **0.03** |
| Frontal (Inf Tri) | 1.76 | 0.4662 | 2.49 | 0.217 | **2.87** | **0.0202** | **2.52** | **0.043** | **2.66** | **0.0179** | **2.9** | **0.0112** |
| Frontal (Inf Oper) | 1.76 | 0.4662 | 1.24 | 0.5564 | **2.43** | **0.0493** | **2.98** | **0.0147** | **2.35** | **0.0299** | **2.46** | **0.0241** |
| Frontal (Inf Orbit) | 1.67 | 0.4662 | 1.95 | 0.4647 | **2.61** | **0.0347** | **2.4** | **0.0496** | **3.14** | **0.0101** | **2.6** | **0.019** |
| Frontal (Med) | 1.33 | 0.5564 | 0.63 | 0.8421 | **2.5** | **0.0443** | 1.36 | 0.2773 | **2.23** | **0.0365** | **2.38** | **0.0289** |
| Frontal (Mid) | 0.99 | 0.7108 | 0.67 | 0.8421 | 1.51 | 0.2131 | 1.78 | 0.1351 | **2.55** | **0.0199** | **2.21** | **0.0372** |
| Frontal (Sup med) | 0.86 | 0.7707 | 1.17 | 0.5875 | 2.32 | 0.0554 | 2.25 | 0.0622 | **2.23** | **0.0365** | **2.66** | **0.0179** |
| Frontal (Sup) | -0.01 | 0.996 | -0.15 | 0.996 | 0.27 | 0.8477 | 0.14 | 0.9356 | **2.15** | **0.0409** | 1.65 | 0.1107 |
| SMA | 1.0 | 0.7108 | 0.52 | 0.8748 | 1.11 | 0.3891 | 0.82 | 0.5302 | **2.61** | **0.019** | **2.38** | **0.0289** |
| Precentral (Med) | -0.16 | 0.996 | 0.29 | 0.9944 | 0.6 | 0.656 | 0.79 | 0.5442 | 1.33 | 0.1941 | 1.36 | 0.1859 |
| Precentral | 0.64 | 0.8421 | 0.49 | 0.894 | 1.34 | 0.2822 | 2.17 | 0.0654 | **2.36** | **0.0293** | **2.51** | **0.0217** |
| Postcentral (Med) | 0.39 | 0.9402 | -0.18 | 0.996 | 1.18 | 0.3473 | 0.64 | 0.6366 | 2.03 | 0.0509 | 1.25 | 0.2195 |

| | | | | | | | | | | | | |
|---|---|---|---|---|---|---|---|---|---|---|---|---|
| Postcentral | 0.57 | 0.8643 | -0.14 | 0.996 | 1.23 | 0.3314 | 1.55 | 0.2033 | **2.28** | **0.0334** | **2.24** | **0.0365** |
| Central Operculum | 2.67 | 0.1977 | 2.22 | 0.3118 | **3.62** | **0.003** | **3.81** | **0.0028** | **3.21** | **0.0101** | **3.08** | **0.0101** |
| Parietal Operculum | 2.28 | 0.3092 | 3.07 | 0.1977 | **3.68** | **0.003** | **4.43** | **0.001** | **2.97** | **0.0104** | **2.85** | **0.0123** |
| Angular Gyrus | -0.01 | 0.996 | 0.79 | 0.7994 | **2.46** | **0.0475** | **3.45** | **0.0044** | **3.08** | **0.0101** | **3.44** | **0.0087** |
| Supramarginal | 0.73 | 0.8345 | 0.99 | 0.7108 | **2.67** | **0.0322** | **4.23** | **0.001** | **3.49** | **0.0087** | **3.29** | **0.0101** |
| Parietal (Sup) | -0.91 | 0.7457 | -0.59 | 0.8598 | 1.87 | 0.1192 | 2.06 | 0.0824 | **3.11** | **0.0101** | **2.82** | **0.0129** |
| Precuneus | 0.14 | 0.996 | 0.32 | 0.9797 | **2.26** | **0.0622** | 2.34 | 0.0532 | **3.03** | **0.0101** | **3.17** | **0.0101** |
| Insula (Ant) | 1.43 | 0.5339 | 1.34 | 0.5564 | **2.76** | **0.0273** | **2.42** | **0.0493** | **3.48** | **0.0087** | **2.97** | **0.0104** |
| Insula (Post) | 2.49 | 0.217 | 1.76 | 0.4662 | **3.72** | **0.003** | **3.08** | **0.013** | **3.26** | **0.0101** | **2.98** | **0.0104** |
| Entorhinal | 0.46 | 0.9028 | 0.01 | 0.996 | 1.71 | 0.1513 | 0.45 | 0.7488 | **1.77** | **0.088** | **2.34** | **0.0302** |
| Fusiform Gyrus | -0.06 | 0.996 | 0.01 | 0.996 | 2.0 | 0.0927 | 1.2 | 0.3402 | **3.02** | **0.0101** | **3.16** | **0.0101** |
| Temporal (Inf) | 0.08 | 0.996 | -0.2 | 0.996 | 2.21 | 0.0622 | 0.34 | 0.8185 | **2.23** | **0.0365** | 1.51 | 0.1456 |
| Temporal (Mid) | 0.53 | 0.8748 | 0.59 | 0.8598 | 2.08 | 0.0789 | 2.22 | 0.0622 | **2.55** | **0.0199** | **3.03** | **0.0101** |
| Temporal (Sup) | 1.28 | 0.5564 | 1.92 | 0.4647 | 2.19 | 0.0634 | **3.44** | **0.0044** | **3.06** | **0.0101** | **3.1** | **0.0101** |
| Temporal Pole | 0.55 | 0.8665 | -0.16 | 0.996 | 1.85 | 0.1216 | 1.38 | 0.2682 | **2.73** | **0.0154** | **2.16** | **0.0402** |
| Transverse temporal | 2.27 | 0.3092 | 1.74 | 0.4662 | **3.66** | **0.003** | **3.61** | **0.003** | **3.18** | **0.0101** | **2.93** | **0.0109** |
| Planum Polare | 2.64 | 0.1977 | 2.07 | 0.3776 | **3.71** | **0.003** | **3.46** | **0.0044** | **3.24** | **0.0101** | **3.47** | **0.0087** |
| Planum Temporale | 2.89 | 0.1977 | 2.77 | 0.1977 | **3.48** | **0.0044** | **4.22** | **0.001** | **2.7** | **0.0163** | **2.54** | **0.0199** |
| Calcarine Fissure | -1.57 | 0.4662 | -1.22 | 0.5573 | -0.21 | 0.8828 | 0.06 | 0.9763 | **2.18** | **0.0387** | **2.31** | **0.0321** |
| Cuneus | -1.63 | 0.4662 | -1.6 | 0.4662 | 0.03 | 0.9763 | 0.26 | 0.8556 | **2.83** | **0.0126** | **2.65** | **0.0179** |
| Lingual Gyrus | -0.93 | 0.7346 | -0.42 | 0.9204 | 0.04 | 0.9763 | 0.44 | 0.7493 | 1.91 | 0.0651 | **2.23** | **0.0365** |
| Occipital fusiform | -1.12 | 0.6261 | -0.81 | 0.7994 | -0.1 | 0.9582 | 0.79 | 0.5442 | 2.01 | 0.0519 | **2.37** | **0.0293** |
| Occipital (Inf) | -1.39 | 0.5339 | -1.28 | 0.5564 | 0.04 | 0.9763 | 0.71 | 0.5947 | 2.03 | 0.0509 | **3.0** | **0.0104** |
| Occipital (Mid) | -1.54 | 0.4786 | -0.65 | 0.8421 | 1.0 | 0.4458 | 0.9 | 0.4927 | **3.06** | **0.0101** | **2.73** | **0.0154** |
| Occipital (Sup) | -1.71 | 0.4662 | -1.71 | 0.4662 | 0.54 | 0.6917 | 0.28 | 0.8477 | **2.2** | **0.0372** | 1.98 | 0.056 |
| Occipital Pole | -1.73 | 0.4662 | -1.59 | 0.4662 | -0.7 | 0.6021 | -0.93 | 0.4761 | 1.28 | 0.2103 | 1.4 | 0.1777 |
| Cerebellum Exterior | -0.05 | 0.996 | 0.46 | 0.9028 | 1.8 | 0.1302 | 1.77 | 0.1372 | **2.15** | **0.0409** | **2.14** | **0.041** |
| Vermis I-V | -0.35 | 0.9567 | - | - | 0.52 | 0.7002 | - | - | 1.39 | 0.1791 | - | - |
| Vermis VI-VII | -0.04 | 0.996 | - | - | 0.68 | 0.6113 | - | - | 1.04 | 0.31 | - | - |
| Vermis VIII-X | -0.01 | 0.996 | - | - | 1.02 | 0.4354 | - | - | 0.75 | 0.4562 | - | - |

Bold numbers highlight regions with significant differences in a specific measure (fALFF, LCOR, or GCOR) between PD and the matched subgroup of HC.

**Supplementary Table 26:** Statistics on regional differences in fALFF, LCOR, and GCOR between PD and HCmatched assessed using the Mann-Whitney-U test – *after atrophy correction*.

| | fALFF | | | | LCOR | | | | GCOR | | | |
|---|---|---|---|---|---|---|---|---|---|---|---|---|
| | Left | | Right | | Left | | Right | | Left | | Right | |
| Region | U statistic | $P_{FDR}$ | U statistic | $P_{FDR}$ | U statistic | $P_{FDR}$ | U statistic | $P_{FDR}$ | U statistic | $P_{FDR}$ | U statistic | $P_{FDR}$ |
| Accumbens | 1.64 | 0.4111 | 1.09 | 0.6462 | **4.08** | **0.0013** | 1.81 | 0.1329 | **3.47** | **0.0103** | **2.54** | **0.0206** |
| Caudate | 2.07 | 0.4111 | 1.63 | 0.4111 | **2.86** | **0.021** | 1.66 | 0.1709 | **3.8** | **0.0087** | **3.19** | **0.0103** |
| Putamen | 1.32 | 0.5456 | 0.56 | 0.8821 | **3.98** | **0.0016** | **2.65** | **0.0347** | **3.8** | **0.0087** | **3.11** | **0.0103** |
| Pallidum | 1.26 | 0.5714 | 0.1 | 0.9934 | **3.42** | **0.0052** | 2.23 | 0.0629 | **2.66** | **0.0183** | **2.92** | **0.011** |
| Amygdala | 0.64 | 0.8632 | -0.14 | 0.9934 | 2.43 | 0.0515 | 0.87 | 0.5169 | **2.58** | **0.02** | 2.03 | 0.051 |
| Hippocampus | 0.55 | 0.8821 | -0.2 | 0.9934 | 2.41 | 0.0515 | 0.83 | 0.5323 | **3.31** | **0.0103** | 2.02 | 0.0519 |
| Ant Cing Gyrus | 0.95 | 0.7393 | 0.79 | 0.8157 | 1.98 | 0.0971 | 2.07 | 0.08 | **2.39** | **0.0291** | **2.77** | **0.0146** |
| Mid Cing Gyrus | 0.76 | 0.8191 | 0.65 | 0.8632 | 1.26 | 0.3172 | 1.32 | 0.2931 | **2.97** | **0.011** | **3.03** | **0.0103** |
| Post Cing Gyrus | -0.19 | 0.9934 | -0.03 | 0.9934 | 0.86 | 0.5178 | 0.57 | 0.6759 | **2.21** | **0.0385** | **2.55** | **0.0206** |
| Parahip. Gyrus | -0.15 | 0.9934 | 0.8 | 0.8112 | 1.51 | 0.2193 | 1.48 | 0.2264 | **2.37** | **0.0299** | **2.53** | **0.0206** |
| Subcallosal Area | 0.23 | 0.9934 | 0.26 | 0.9934 | 2.29 | 0.0593 | -0.34 | 0.8207 | **2.06** | **0.0481** | 0.47 | 0.6365 |
| Thalamus | 1.16 | 0.6016 | 0.64 | 0.8632 | 1.63 | 0.1783 | 0.95 | 0.4721 | **2.73** | **0.0156** | **2.29** | **0.0338** |
| Basal Forebrain | 0.09 | 0.9934 | 1.2 | 0.5797 | 2.34 | 0.0577 | 2.29 | 0.0593 | **2.06** | **0.0481** | **2.77** | **0.0146** |
| Frontal Pole | 0.38 | 0.9431 | -0.16 | 0.9934 | 0.48 | 0.723 | 0.29 | 0.8306 | 0.76 | 0.4553 | 0.94 | 0.3567 |
| Gyrus Rectus | 0.41 | 0.943 | -0.09 | 0.9934 | 1.04 | 0.4274 | -0.07 | 0.9743 | 1.35 | 0.193 | **2.58** | **0.02** |
| Frontal Operculum | 1.77 | 0.4111 | 1.49 | 0.5098 | **3.0** | **0.0153** | 2.21 | 0.0633 | **3.1** | **0.0103** | **2.54** | **0.0206** |
| Ant Orbital | 1.38 | 0.5419 | 0.8 | 0.8112 | **2.63** | **0.0349** | 2.27 | 0.0606 | **2.94** | **0.011** | **2.19** | **0.0388** |
| Lat Orbital | 1.63 | 0.4111 | 1.44 | 0.5406 | **3.07** | **0.0142** | **3.04** | **0.0142** | **3.05** | **0.0103** | **2.85** | **0.0127** |
| Med Orbital | 0.76 | 0.8191 | 0.93 | 0.7401 | 2.25 | 0.062 | 2.4 | 0.0515 | 1.71 | 0.0987 | **2.93** | **0.011** |
| Post Orbital | 1.41 | 0.5419 | 1.21 | 0.5797 | **2.7** | **0.0321** | 2.32 | 0.0583 | **2.6** | **0.0196** | **2.33** | **0.0312** |
| Frontal (Inf Tri) | 1.73 | 0.4111 | 2.4 | 0.2822 | **2.87** | **0.021** | **2.49** | **0.0471** | **2.66** | **0.0183** | **2.91** | **0.011** |
| Frontal (Inf Oper) | 1.76 | 0.4111 | 1.25 | 0.5714 | 2.4 | 0.0515 | **2.99** | **0.0153** | **2.33** | **0.0312** | **2.45** | **0.0247** |
| Frontal (Inf Orbit) | 1.71 | 0.4111 | 1.91 | 0.4111 | **2.62** | **0.0351** | 2.38 | 0.0524 | **3.14** | **0.0103** | **2.6** | **0.0196** |
| Frontal (Med) | 1.36 | 0.5419 | 0.58 | 0.8821 | **2.52** | **0.0444** | 1.32 | 0.2931 | **2.23** | **0.038** | **2.34** | **0.0312** |
| Frontal (Mid) | 0.93 | 0.7401 | 0.59 | 0.8821 | 1.5 | 0.2193 | 1.74 | 0.1467 | **2.55** | **0.0206** | **2.2** | **0.0388** |
| Frontal (Sup med) | 0.84 | 0.7973 | 1.16 | 0.6016 | 2.31 | 0.0593 | 2.23 | 0.0629 | **2.22** | **0.038** | **2.64** | **0.0185** |
| Frontal (Sup) | 0.01 | 0.9934 | -0.14 | 0.9934 | 0.29 | 0.8306 | 0.17 | 0.9147 | **2.15** | **0.0411** | 1.69 | 0.1023 |
| SMA | 1.02 | 0.7021 | 0.5 | 0.887 | 1.12 | 0.3805 | 0.81 | 0.5409 | **2.62** | **0.0192** | **2.38** | **0.0294** |
| Precentral (Med) | -0.19 | 0.9934 | 0.21 | 0.9934 | 0.58 | 0.6715 | 0.74 | 0.5916 | 1.32 | 0.1981 | 1.33 | 0.1959 |
| Precentral | 0.54 | 0.8821 | 0.39 | 0.9431 | 1.29 | 0.3065 | 2.12 | 0.0757 | **2.32** | **0.0313** | **2.46** | **0.0247** |
| Postcentral (Med) | 0.35 | 0.9582 | -0.17 | 0.9934 | 1.17 | 0.3627 | 0.66 | 0.6204 | 2.03 | 0.0512 | 1.29 | 0.2048 |

| | | | | | | | | | | | | |
|---|---|---|---|---|---|---|---|---|---|---|---|---|
| Postcentral | 0.53 | 0.8821 | -0.19 | 0.9934 | 1.21 | 0.343 | 1.51 | 0.2193 | **2.24** | **0.0373** | **2.2** | **0.0388** |
| Central Operculum | 2.61 | 0.2126 | 2.22 | 0.3521 | **3.57** | **0.0035** | **3.81** | **0.0027** | **3.19** | **0.0103** | **3.09** | **0.0103** |
| Parietal Operculum | 2.25 | 0.3521 | 3.03 | 0.2126 | **3.63** | **0.0035** | **4.4** | **0.0013** | **2.92** | **0.011** | **2.81** | **0.0137** |
| Angular Gyrus | -0.17 | 0.9934 | 0.53 | 0.8821 | 2.4 | 0.0515 | **3.35** | **0.0056** | **3.07** | **0.0103** | **3.41** | **0.0103** |
| Supramarginal | 0.64 | 0.8632 | 0.84 | 0.7973 | **2.65** | **0.0347** | **4.17** | **0.0013** | **3.49** | **0.0103** | **3.26** | **0.0103** |
| Parietal (Sup) | -0.98 | 0.7201 | -0.72 | 0.8502 | 1.83 | 0.1284 | 1.99 | 0.095 | **3.06** | **0.0103** | **2.79** | **0.0143** |
| Precuneus | 0.03 | 0.9934 | 0.23 | 0.9934 | 2.2 | 0.0633 | 2.28 | 0.0593 | **3.02** | **0.0103** | **3.17** | **0.0103** |
| Insula (Ant) | 1.38 | 0.5419 | 1.32 | 0.5456 | **2.72** | **0.0307** | 2.43 | 0.0515 | **3.44** | **0.0103** | **2.97** | **0.011** |
| Insula (Post) | 2.41 | 0.2822 | 1.65 | 0.4111 | **3.67** | **0.0035** | **3.05** | **0.0142** | **3.22** | **0.0103** | **2.94** | **0.011** |
| Entorhinal | 0.47 | 0.9075 | 0.05 | 0.9934 | 1.71 | 0.1537 | 0.48 | 0.723 | 1.75 | 0.0924 | **2.33** | **0.0312** |
| Fusiform Gyrus | -0.2 | 0.9934 | -0.08 | 0.9934 | 1.87 | 0.1227 | 1.13 | 0.3781 | **2.91** | **0.011** | **3.1** | **0.0103** |
| Temporal (Inf) | 0.02 | 0.9934 | -0.23 | 0.9934 | 2.08 | 0.08 | 0.29 | 0.8306 | **2.18** | **0.0392** | 1.5 | 0.1492 |
| Temporal (Mid) | 0.5 | 0.887 | 0.52 | 0.887 | 2.08 | 0.08 | 2.2 | 0.0633 | **2.56** | **0.0206** | **3.02** | **0.0103** |
| Temporal (Sup) | 1.22 | 0.5797 | 1.82 | 0.4111 | 2.14 | 0.0718 | **3.39** | **0.0056** | **3.03** | **0.0103** | **3.05** | **0.0103** |
| Temporal Pole | 0.67 | 0.8632 | -0.08 | 0.9934 | 1.85 | 0.1242 | 1.37 | 0.2739 | **2.72** | **0.0156** | **2.17** | **0.0395** |
| Transverse temporal | 2.17 | 0.3536 | 1.65 | 0.4111 | **3.59** | **0.0035** | **3.57** | **0.0035** | **3.1** | **0.0103** | **2.89** | **0.0114** |
| Planum Polare | 2.62 | 0.2126 | 2.04 | 0.4111 | **3.67** | **0.0035** | **3.48** | **0.0046** | **3.23** | **0.0103** | **3.47** | **0.0103** |
| Planum Temporale | 2.77 | 0.2126 | 2.63 | 0.2126 | **3.37** | **0.0056** | **4.12** | **0.0013** | **2.62** | **0.0192** | **2.49** | **0.0232** |
| Calcarine Fissure | -1.71 | 0.4111 | -1.32 | 0.5456 | -0.29 | 0.8306 | -0.01 | 0.9917 | **2.11** | **0.0445** | **2.25** | **0.037** |
| Cuneus | -1.67 | 0.4111 | -1.68 | 0.4111 | 0.02 | 0.9917 | 0.24 | 0.8628 | **2.83** | **0.0131** | **2.65** | **0.0185** |
| Lingual Gyrus | -1.01 | 0.7059 | -0.54 | 0.8821 | -0.01 | 0.9917 | 0.37 | 0.809 | 1.86 | 0.0722 | **2.18** | **0.0392** |
| Occipital fusiform | -1.14 | 0.6093 | -0.9 | 0.7516 | -0.14 | 0.9293 | 0.72 | 0.596 | 2.01 | 0.0528 | **2.35** | **0.0307** |
| Occipital (Inf) | -1.36 | 0.5419 | -1.29 | 0.5565 | 0.02 | 0.9917 | 0.7 | 0.5995 | **2.06** | **0.0481** | **3.01** | **0.0104** |
| Occipital (Mid) | -1.67 | 0.4111 | -0.71 | 0.8506 | 0.97 | 0.4633 | 0.89 | 0.5055 | **3.06** | **0.0103** | **2.73** | **0.0156** |
| Occipital (Sup) | -1.74 | 0.4111 | -1.77 | 0.4111 | 0.54 | 0.6931 | 0.29 | 0.8306 | **2.19** | **0.0388** | 1.98 | 0.0556 |
| Occipital Pole | -1.74 | 0.4111 | -1.57 | 0.4442 | -0.7 | 0.5995 | -0.92 | 0.4885 | 1.29 | 0.2048 | 1.42 | 0.17 |
| Cerebellum Exterior | -0.05 | 0.9934 | 0.46 | 0.9075 | 1.78 | 0.1385 | 1.77 | 0.1407 | **2.11** | **0.0445** | **2.12** | **0.0441** |
| Vermis I-V | -0.38 | 0.9431 | - | - | 0.48 | 0.723 | - | - | 1.35 | 0.1916 | - | - |
| Vermis VI-VII | -0.05 | 0.9934 | - | - | 0.66 | 0.6204 | - | - | 1.01 | 0.3212 | - | - |
| Vermis VIII-X | 0.01 | 0.9934 | - | - | 1.03 | 0.4299 | - | - | 0.75 | 0.4557 | - | - |

Bold numbers highlight regions with significant differences in a specific measure (fALFF, LCOR, or GCOR) between PD and the matched subgroup of HC.

### Supplementary Figures

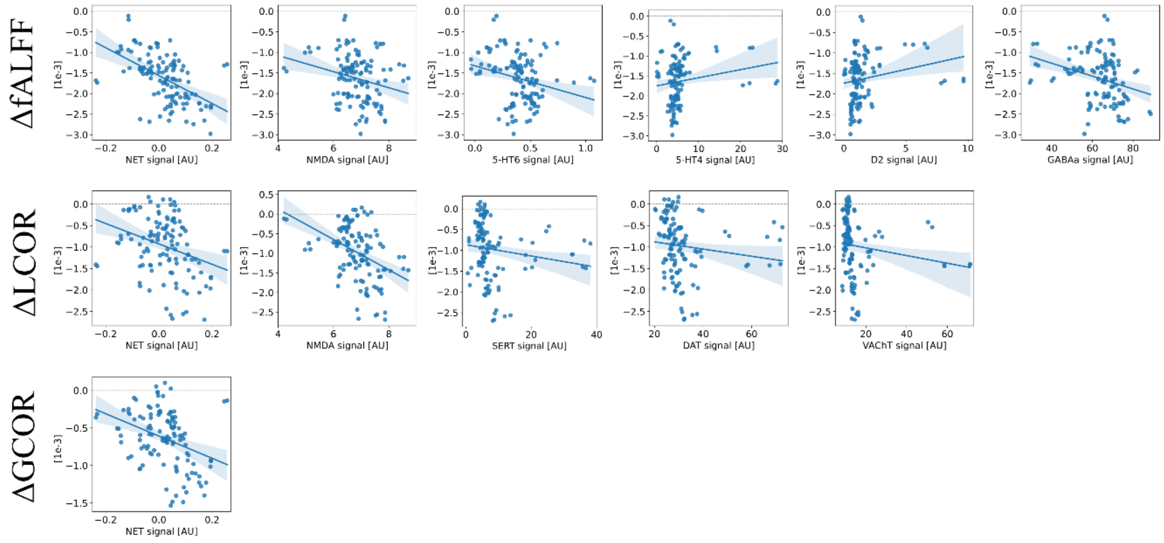


**Supplementary Figure 1**: Significant linear correlations between unthresholded age-effect (slope) maps in fALFF, LCOR, and GCOR, and neurotransmitter systems – *before atrophy correction*. Corresponding correlation coefficients are visualized in Figure 2B. ΔMeasure corresponds to the annual rate of in- or decrease in the respective measure of brain function (fALFF, LCOR, or GCOR) within one region.

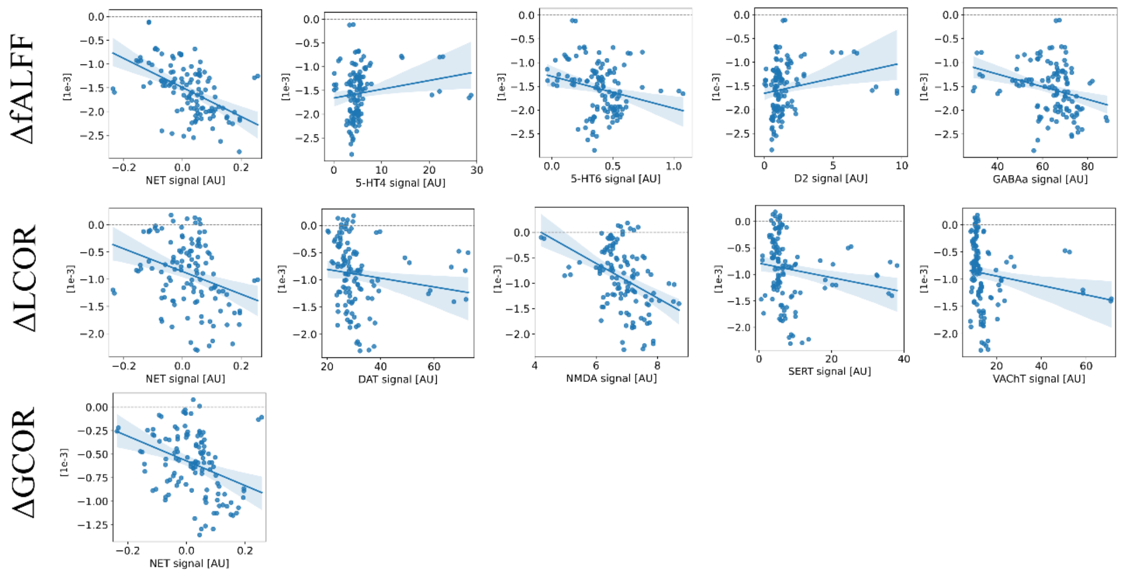


**Supplementary Figure 2**: Significant linear correlations between of unthresholded age-effects (slope maps) in fALFF, LCOR, and GCOR, and multiple neurotransmitter systems – *after atrophy correction*. ΔMeasure corresponds to the annual rate of in- or decrease in the respective measure of brain function (fALFF, LCOR, or GCOR) within one region.

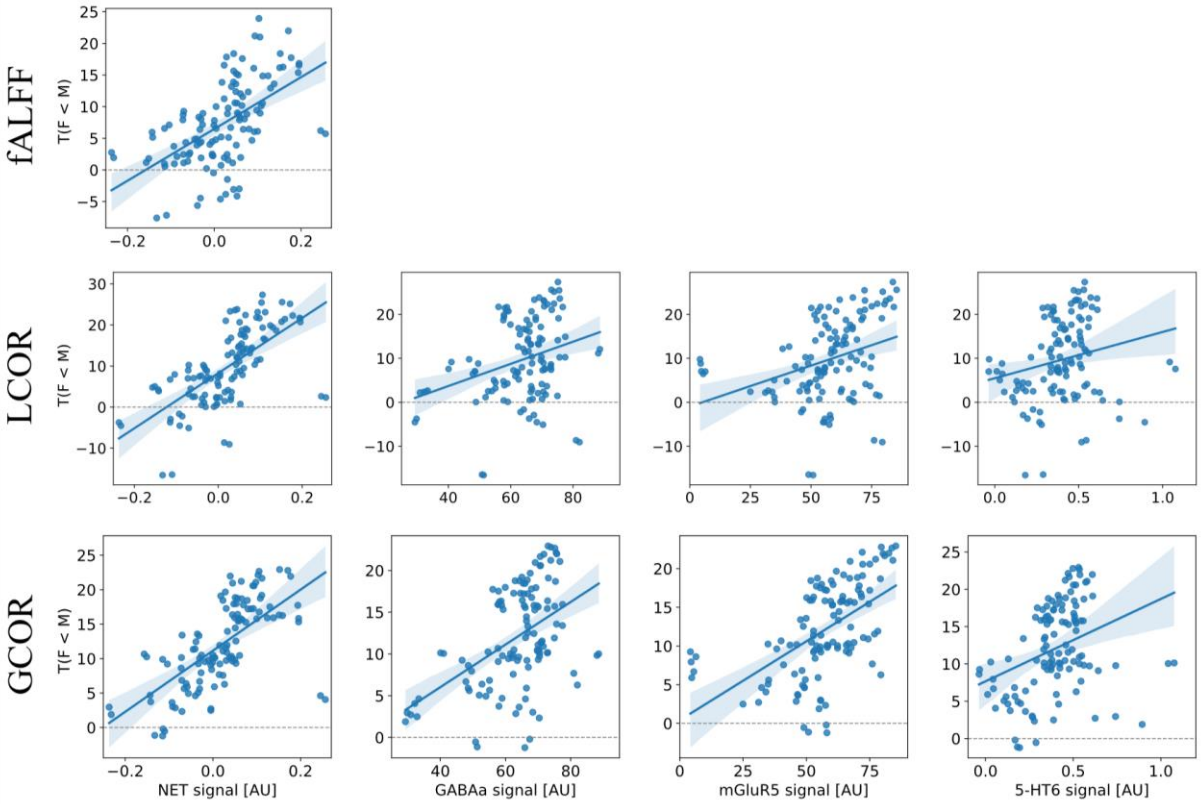


**Supplementary Figure 3:** Significant linear correlations of unthresholded sex effects (T-values) in fALFF, LCOR, and GCOR and NET, GABAa, mGluR5, and 5-HT6 availability – *before atrophy correction*.

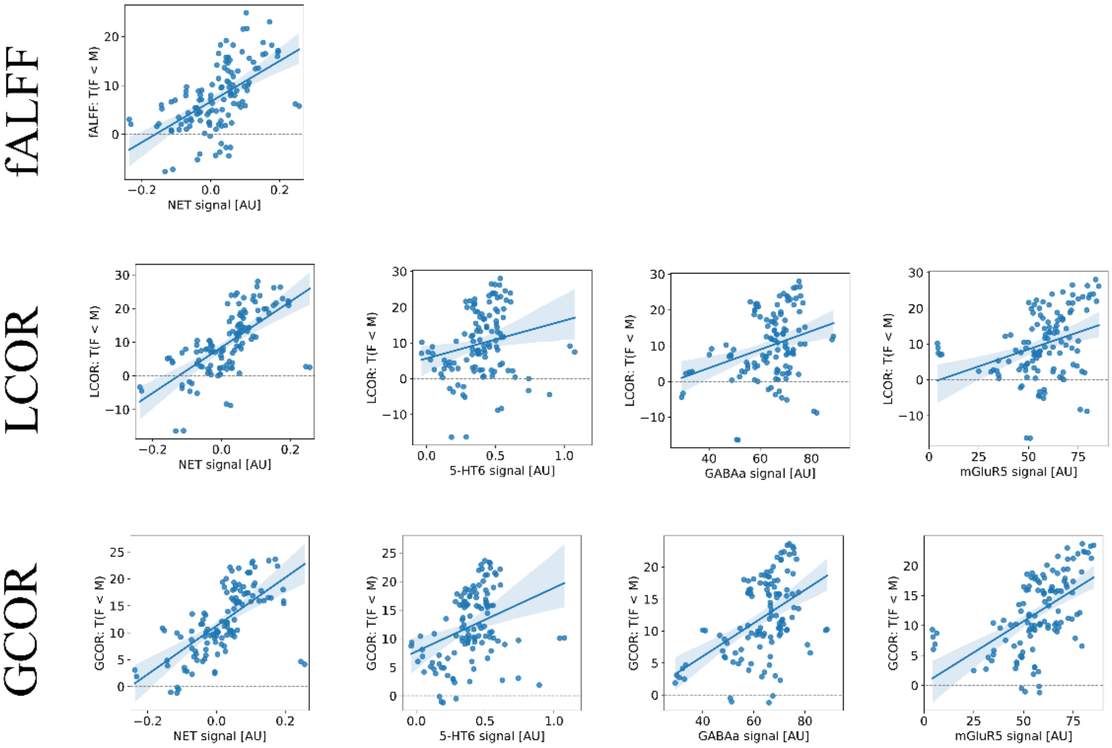


**Supplementary Figure 4:** Significant linear correlations of unthresholded sex effects (T-values) in fALFF, LCOR, and GCOR and NET, GABAa, mGluR5, and 5-HT6 availability – *after atrophy correction*.

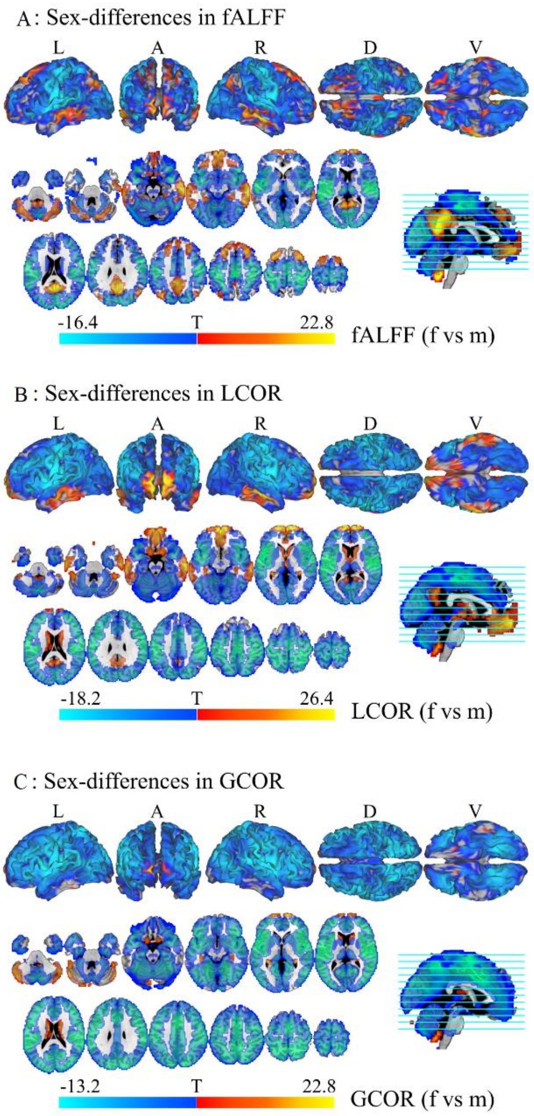


**Supplementary Figure 5**: Sex differences (T-values) in brain functional measures, thresholded. Red voxels indicate higher values in women compared to men and blue voxels indicate the inverse.

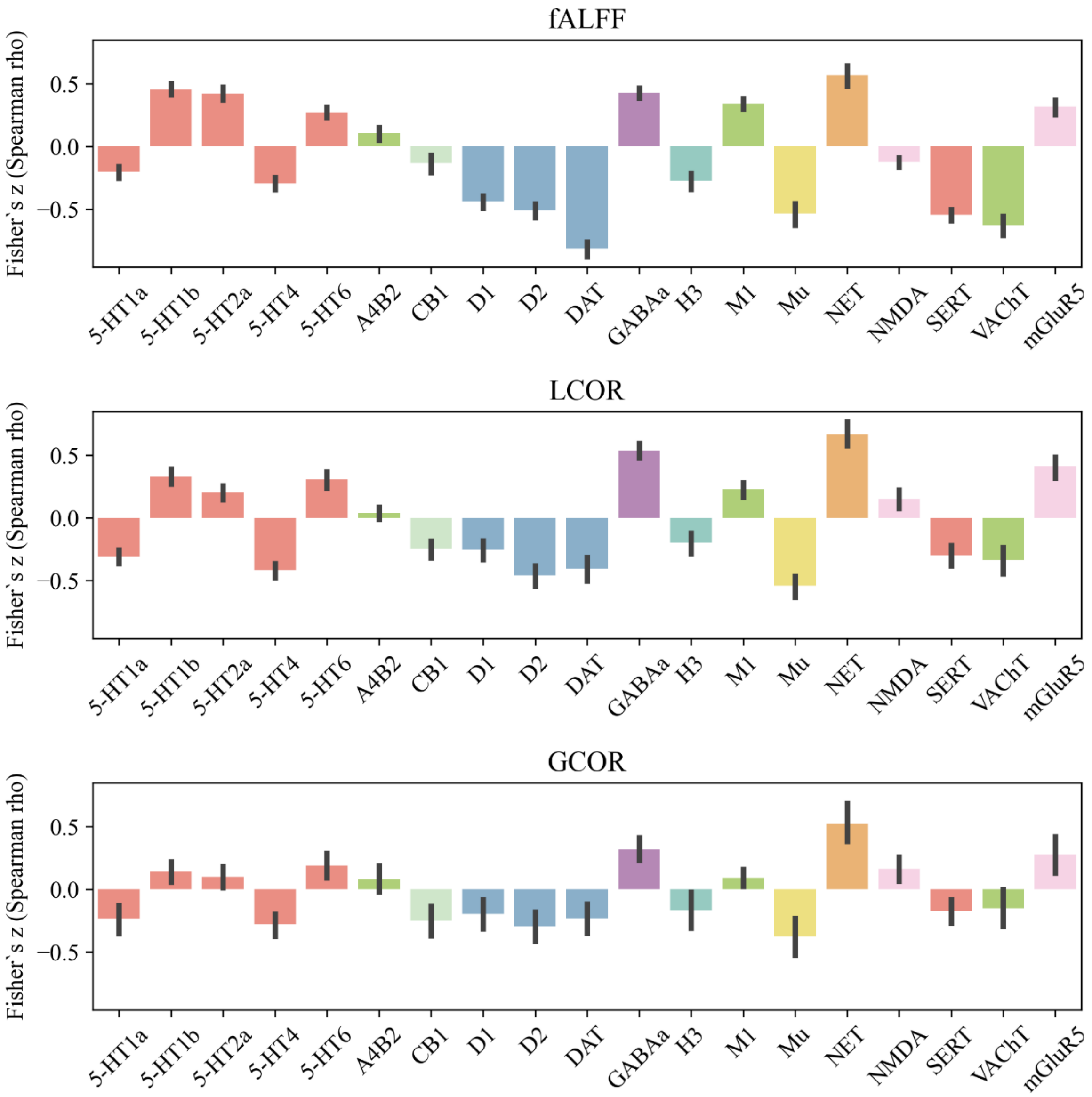


**Supplementary Figure 6**. Bar plots show the healthy control cohort's mean Fisher's z-transformed Spearman correlation coefficients. These coefficients were derived from the spatial correlation analyses of individual fALFF (top), LCOR (middle), and GCOR (bottom) maps and 19 PET maps of neurotransmitter systems. Vertical bars indicate the 50% quantile. Colors group receptors and transporters of the same neurotransmitter system, i.e., serotonin (red), dopamine (blue), acetylcholine (green), glutamate (pink) and GABA (purple), cannabinoid (mint), opioid (yellow), norepinephrine (orange), and histamine (turquoise).

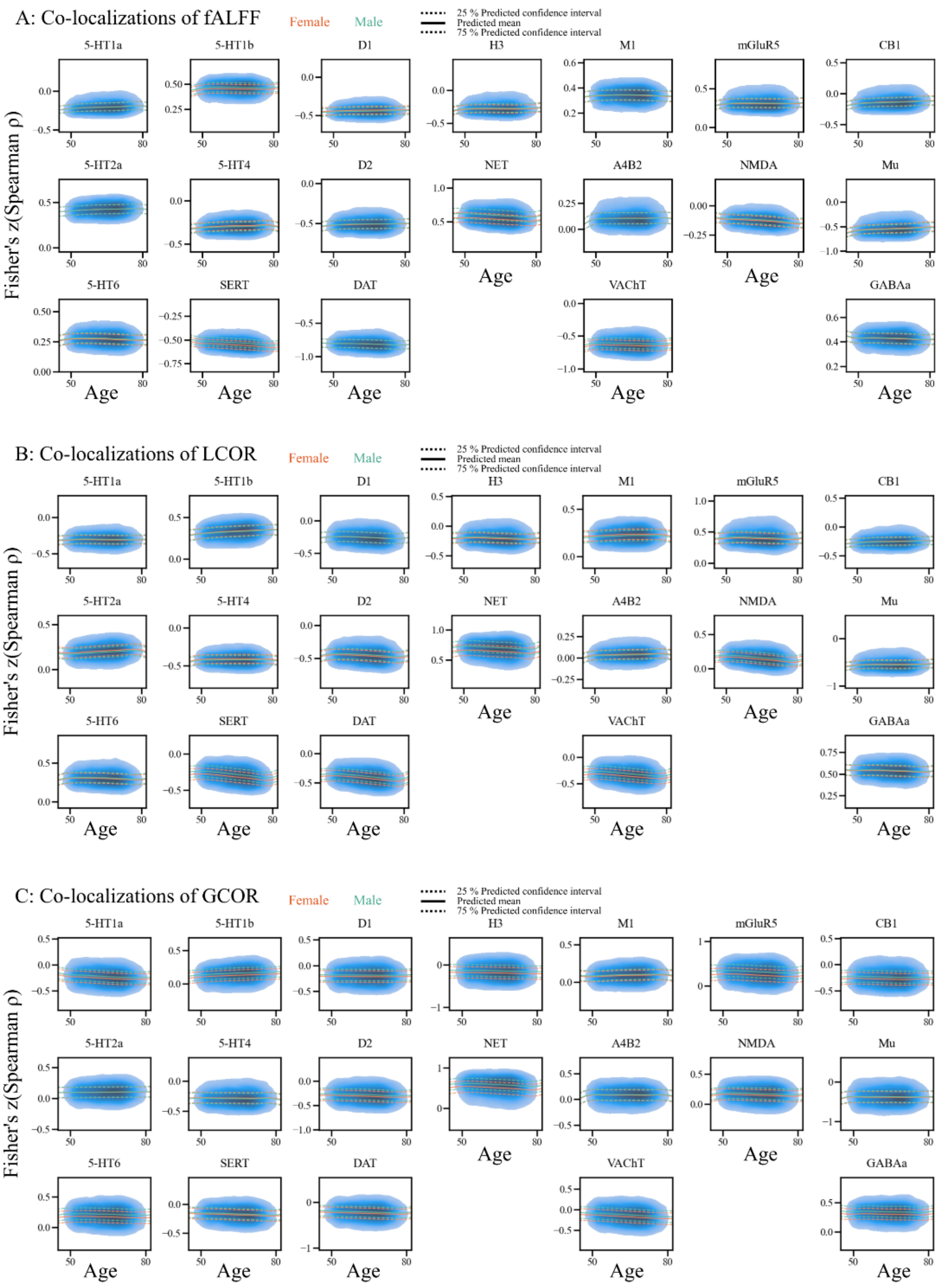


**Supplementary Figure 7:** Normative models of co-localizations between brain function (A: fALFF, B: LCOR, C: GCOR) and 19 PET maps – *before atrophy correction*. Within each subplot, the blue cloud (kernel density estimation) visualized the distribution of Fisher's z-transformed Spearman correlation coefficients of all healthy controls. Solid and dashed lines indicate the predicted median and 25 and 75% confidence interval of men (turquoise) and women (orange).

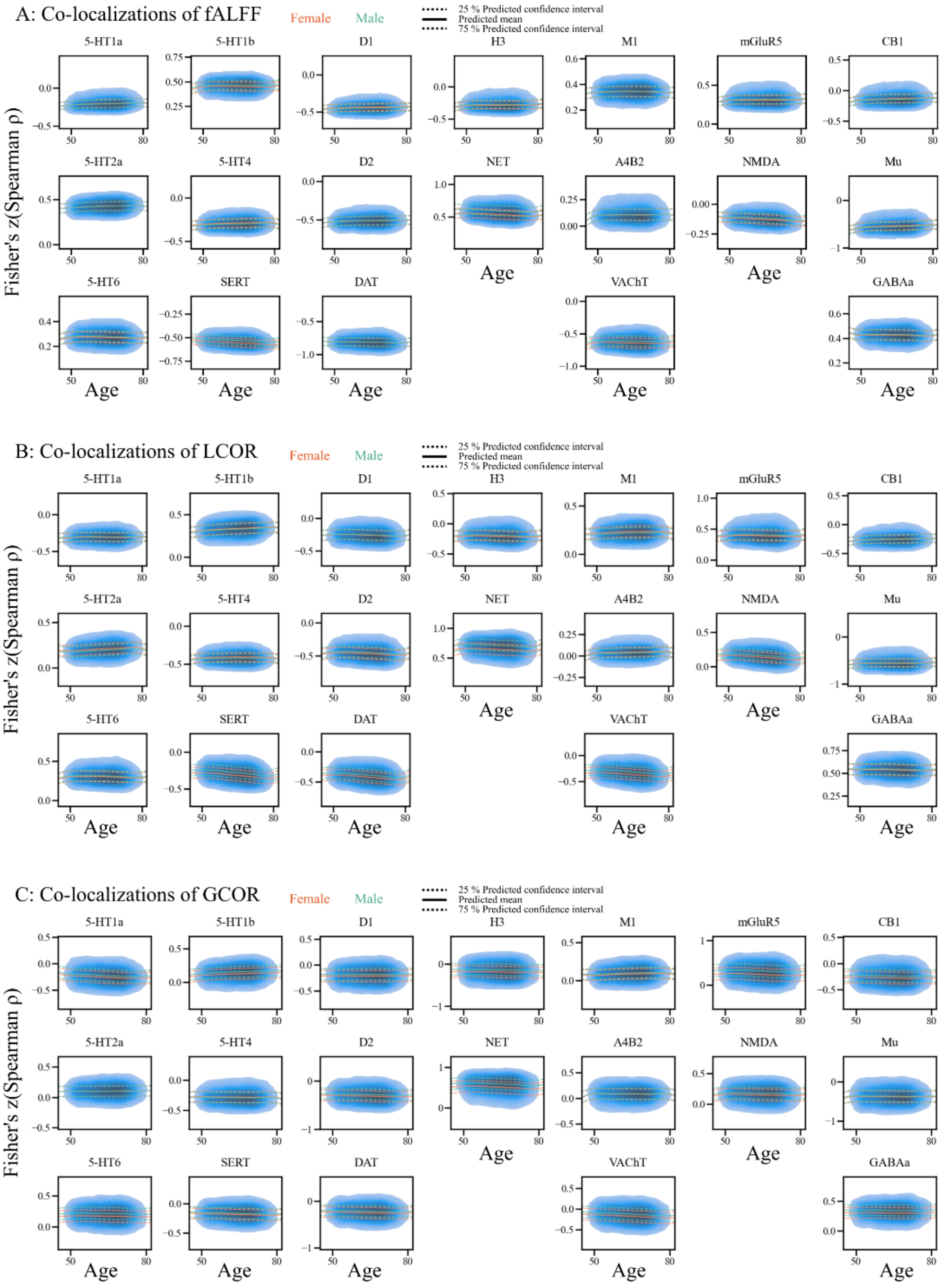


**Supplementary Figure 8:** Normative models of co-localizations between brain function (A: fALFF, B: LCOR, C: GCOR) and 19 PET maps – *after atrophy correction*. Within each subplot, the blue cloud (kernel density estimation) visualized the distribution of Fisher's z-transformed Spearman correlation coefficients of all healthy controls. Solid and dashed lines indicate the predicted median and 25 and 75% confidence interval of men (turquoise) and women (orange).

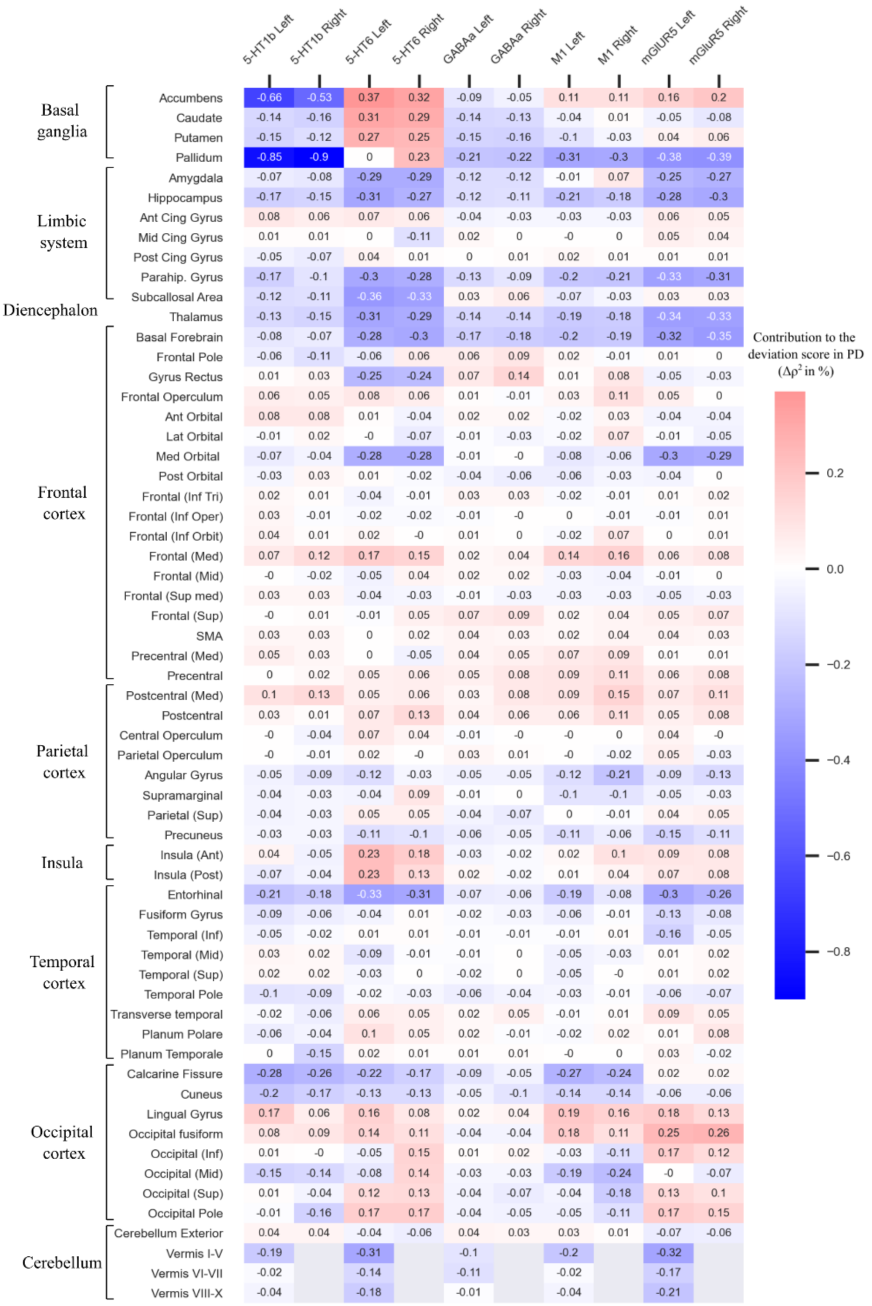


**Supplementary Figure 9**: Contribution of each region to the deviation in **fALFF** co-localizations in subjects with manifest Parkinson's disease – *before atrophy correction*. The contribution is quantified by the mean change in squared spatial correlation coefficient (mean $\Delta\rho^2$) after leaving the specific region out from the spatial correlation analysis. Values of regions of the left or right hemisphere are arranged next to each other (column-wise) for each neurotransmitter system. The rows are sorted from top to the bottom: Basal ganglia, limbic system, diencephalon, frontal cortex, motor area, parietal cortex, temporal cortex, occipital cortex, cerebellum. Red cells, i.e. positive values, indicate that leaving this specific region out in the individual co-localization analysis led to a correlation coefficient that was closer to the norm.

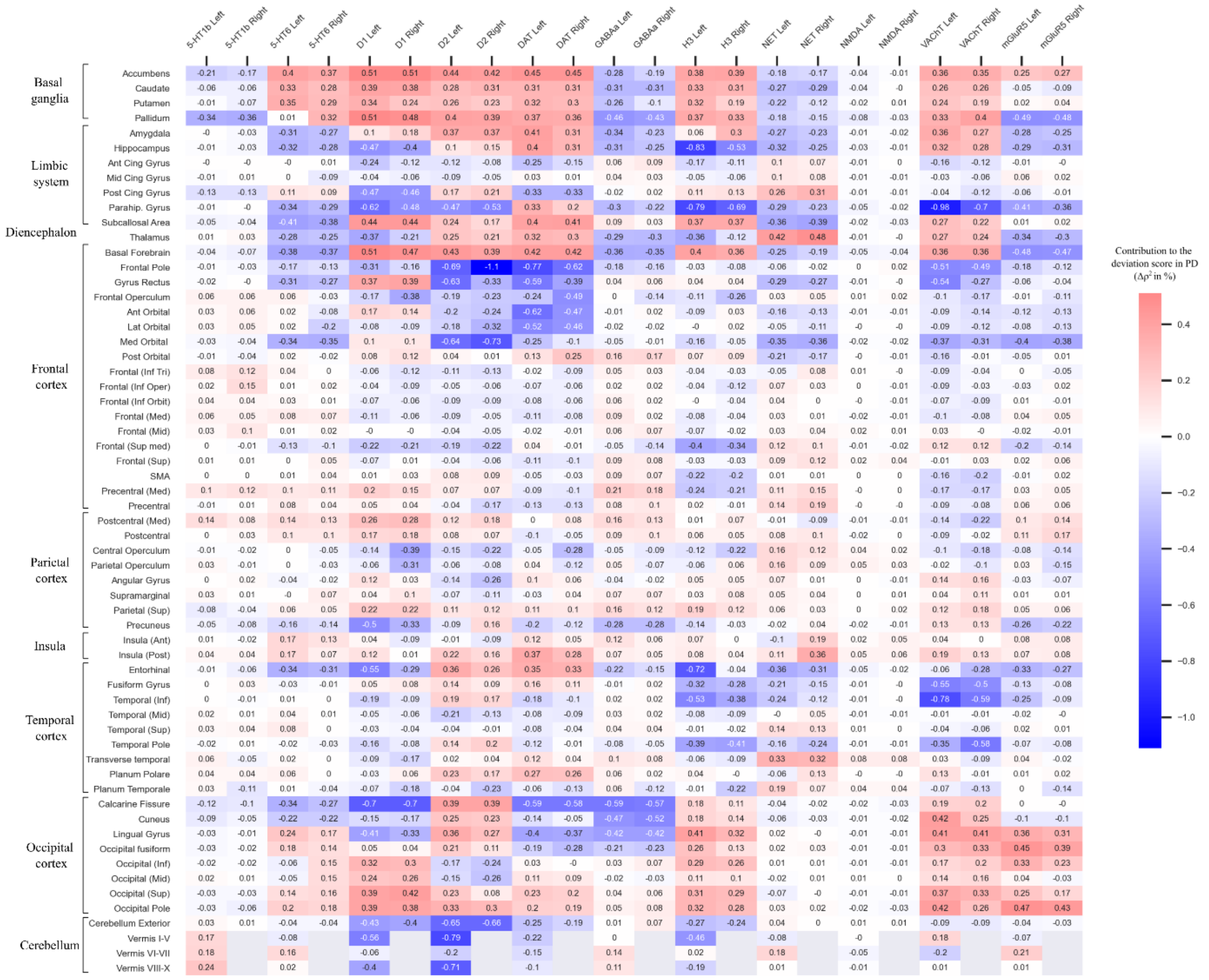


**Supplementary Figure 10:** Contribution of each region to the deviation in **LCOR** co-localizations in subjects with manifest Parkinson's disease – *before atrophy correction*. The contribution is quantified by the mean change in squared spatial correlation coefficient (mean $\Delta\rho^2$) after leaving the specific region out from the spatial correlation analysis. Values of regions of the left or right hemisphere are arranged next to each other (column-wise) for each neurotransmitter system. The rows are sorted from top to the bottom: Basal ganglia, limbic system, diencephalon, frontal cortex, parietal cortex, temporal cortex, occipital cortex, cerebellum. Red cells, i.e. positive values, indicate that leaving this specific region out in the individual co-localization analysis led to a correlation coefficient that was closer to the norm.

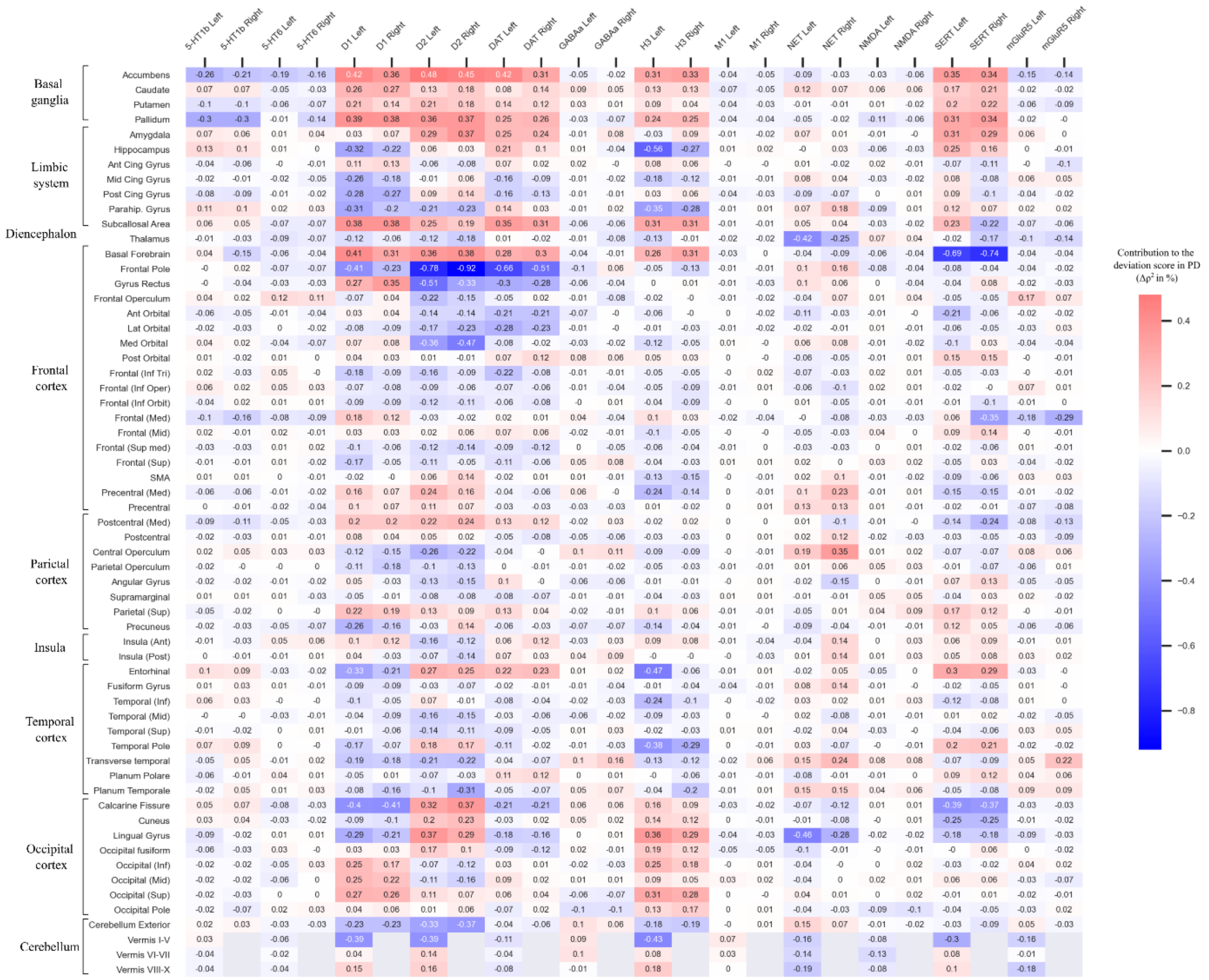


**Supplementary Figure 11**: Contribution of each region to the deviation in **GCOR** co-localizations in subjects with manifest Parkinson's disease – *before atrophy correction*. The contribution is quantified by the mean change in squared spatial correlation coefficient (mean $\Delta\rho^2$) after leaving the specific region out from the spatial correlation analysis. Values of regions of the left or right hemisphere are arranged next to each other (column-wise) for each neurotransmitter system. The rows are sorted from top to the bottom: Basal ganglia, limbic system, diencephalon, frontal cortex, parietal cortex, temporal cortex, occipital cortex, cerebellum. Red cells, i.e. positive values, indicate that leaving this specific region out in the individual co-localization analysis led to a correlation coefficient that was closer to the norm.

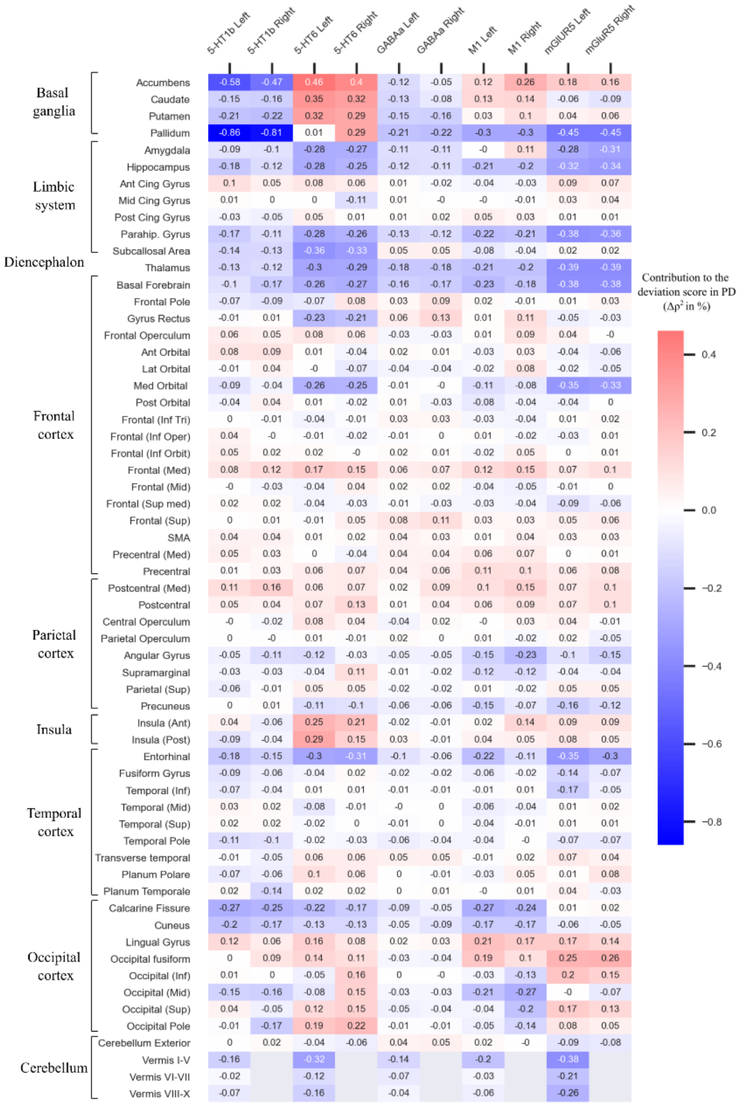


**Supplementary Figure 12**: Contribution of each region to the deviation in **fALFF** co-localizations in subjects with manifest Parkinson's disease – *after atrophy correction*. The contribution is quantified by the mean change in squared spatial correlation coefficient (mean $\Delta\rho^2$) after leaving the specific region out from the spatial correlation analysis. Values of regions of the left or right hemisphere are arranged next to each other (column-wise) for each neurotransmitter system. The rows are sorted from top to the bottom: Basal ganglia, limbic system, diencephalon, frontal cortex, parietal cortex, temporal cortex, occipital cortex, cerebellum. Red cells, i.e. positive values, indicate that leaving this specific region out in the individual co-localization analysis led to a correlation coefficient that was closer to the norm.

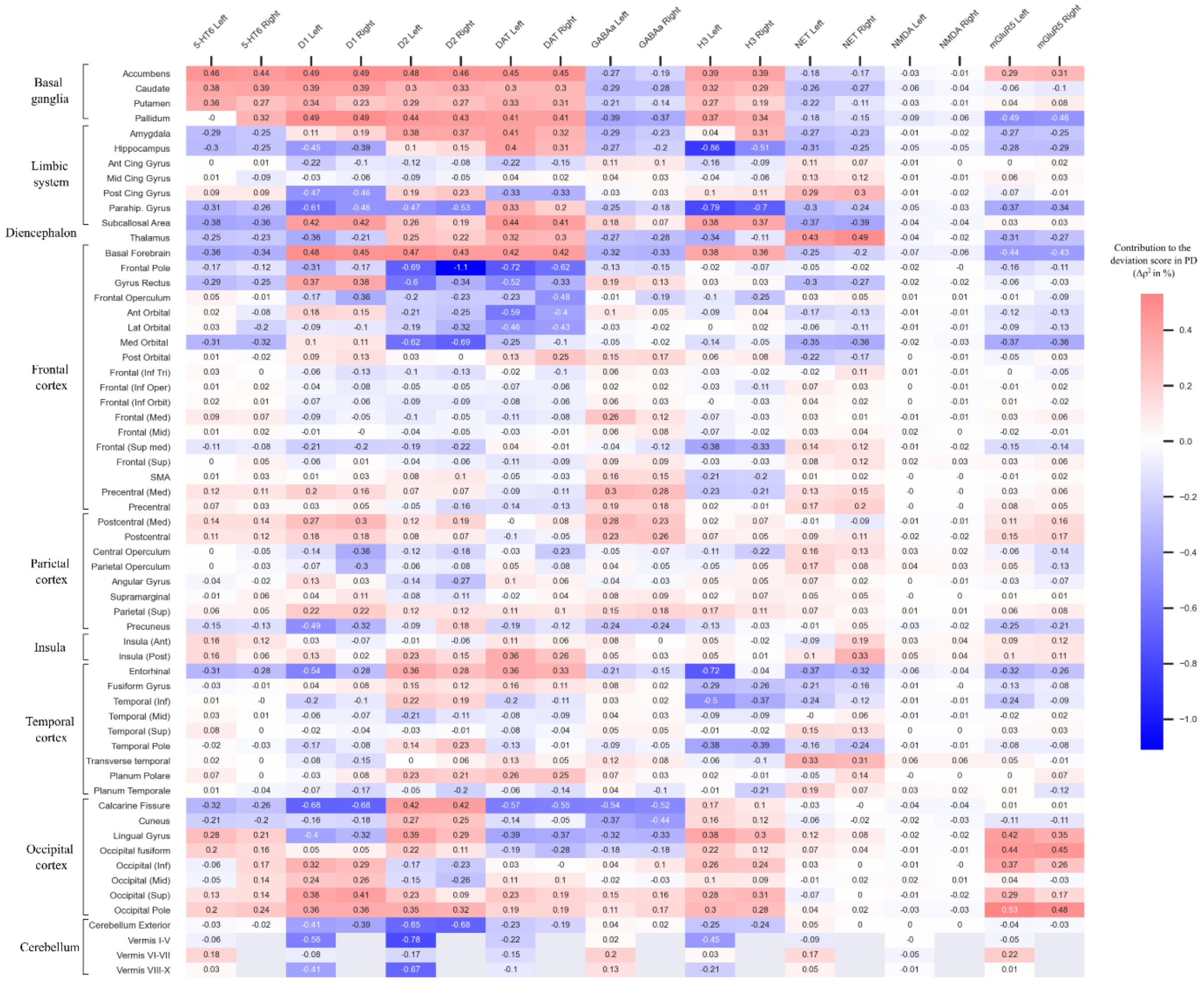


**Supplementary Figure 13:** Contribution of each region to the deviation in **LCOR** co-localizations in subjects with manifest Parkinson's disease – *after atrophy correction*. The contribution is quantified by the mean change in squared spatial correlation coefficient (mean $\Delta\rho^2$) after leaving the specific region out from the spatial correlation analysis. Values of regions of the left or right hemisphere are arranged next to each other (column-wise) for each neurotransmitter system. The rows are sorted from top to the bottom: Basal ganglia, limbic system, diencephalon, frontal cortex, parietal cortex, temporal cortex, occipital cortex, cerebellum. Red cells, i.e. positive values, indicate that leaving this specific region out in the individual co-localization analysis led to a correlation coefficient that was closer to the norm.

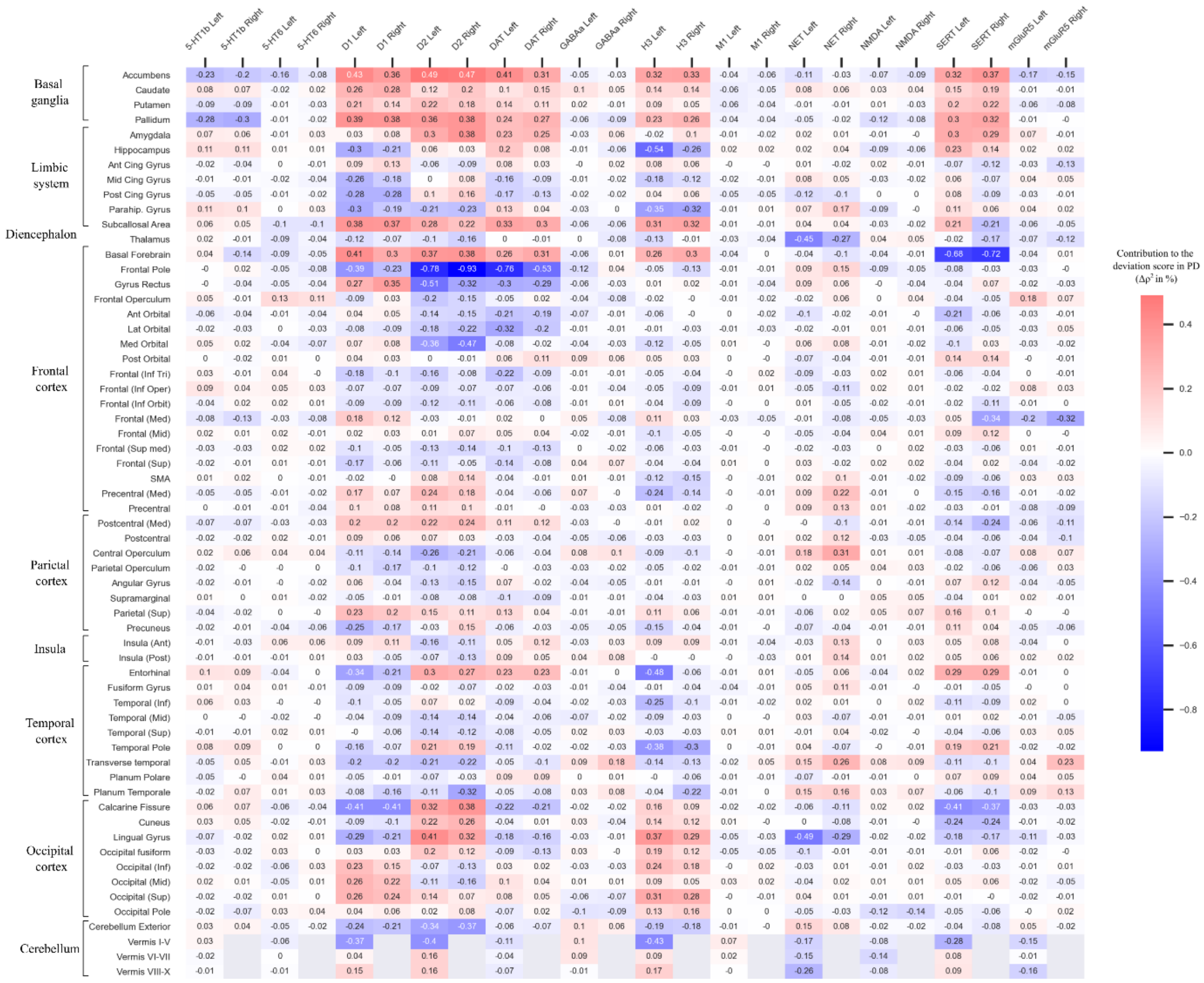


**Supplementary Figure 14**: Contribution of each region to the deviation in **GCOR** co-localizations in subjects with manifest Parkinson's disease – *after atrophy correction*. The contribution is quantified by the mean change in squared spatial correlation coefficient (mean $\Delta\rho^2$) after leaving the specific region out from the spatial correlation analysis. Values of regions of the left or right hemisphere are arranged next to each other (column-wise) for each neurotransmitter system. The rows are sorted from top to the bottom: Basal ganglia, limbic system, diencephalon, frontal cortex, parietal cortex, temporal cortex, occipital cortex, cerebellum. Red cells, i.e. positive values, indicate that leaving this specific region out in the individual co-localization analysis led to a correlation coefficient that was closer to the norm.

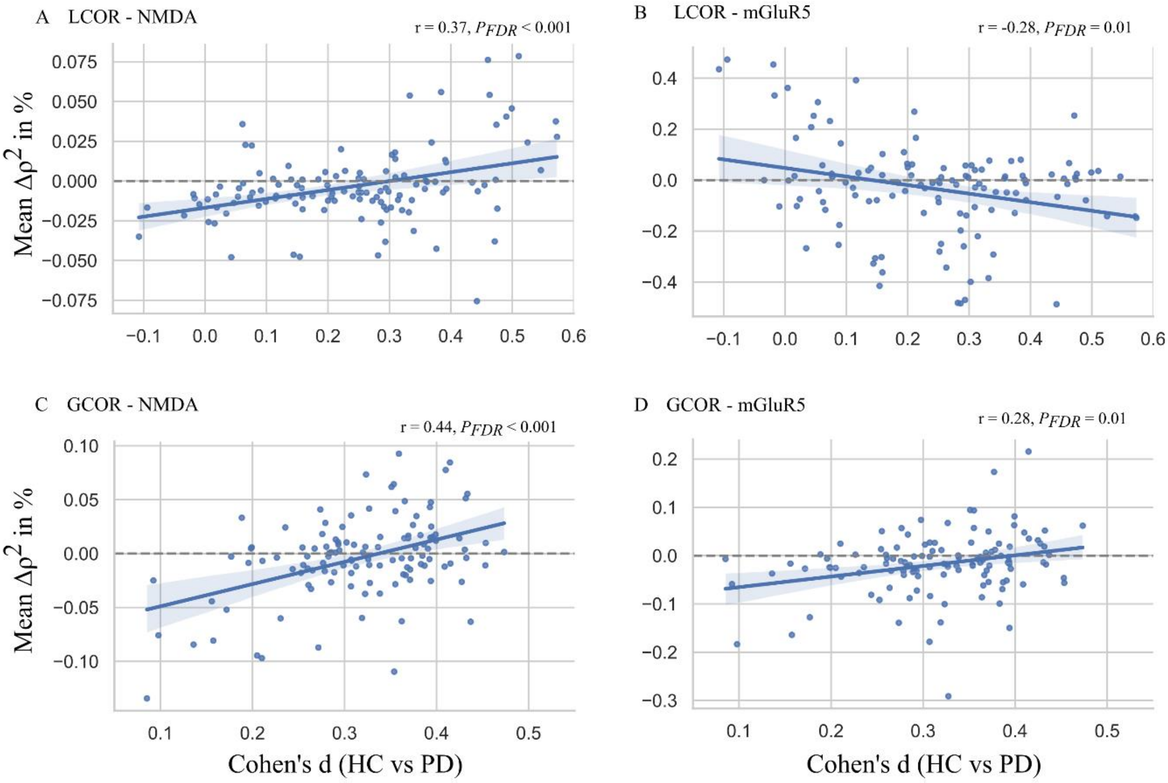


**Supplementary Figure 15:** Linear correlation of regional contribution (mean $\Delta\rho^2$) to the deviation score and functional differences (Cohen's d) between PD and the matched subcohort of healthy controls – *before atrophy correction*.

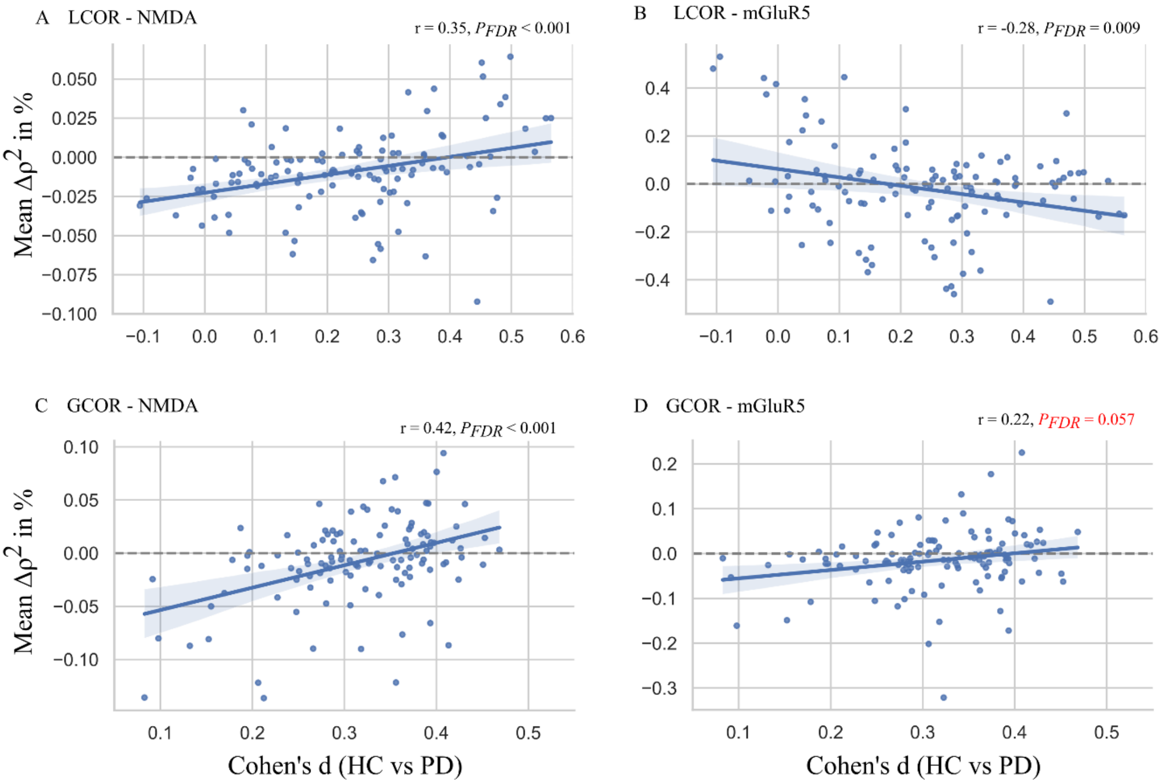


**Supplementary Figure 16**: Linear correlation of regional contribution (Mean $\Delta\rho^2$) to deviation score and functional differences (Cohen's d) between PD and the matched subcohort of healthy controls – *after atrophy correction*.

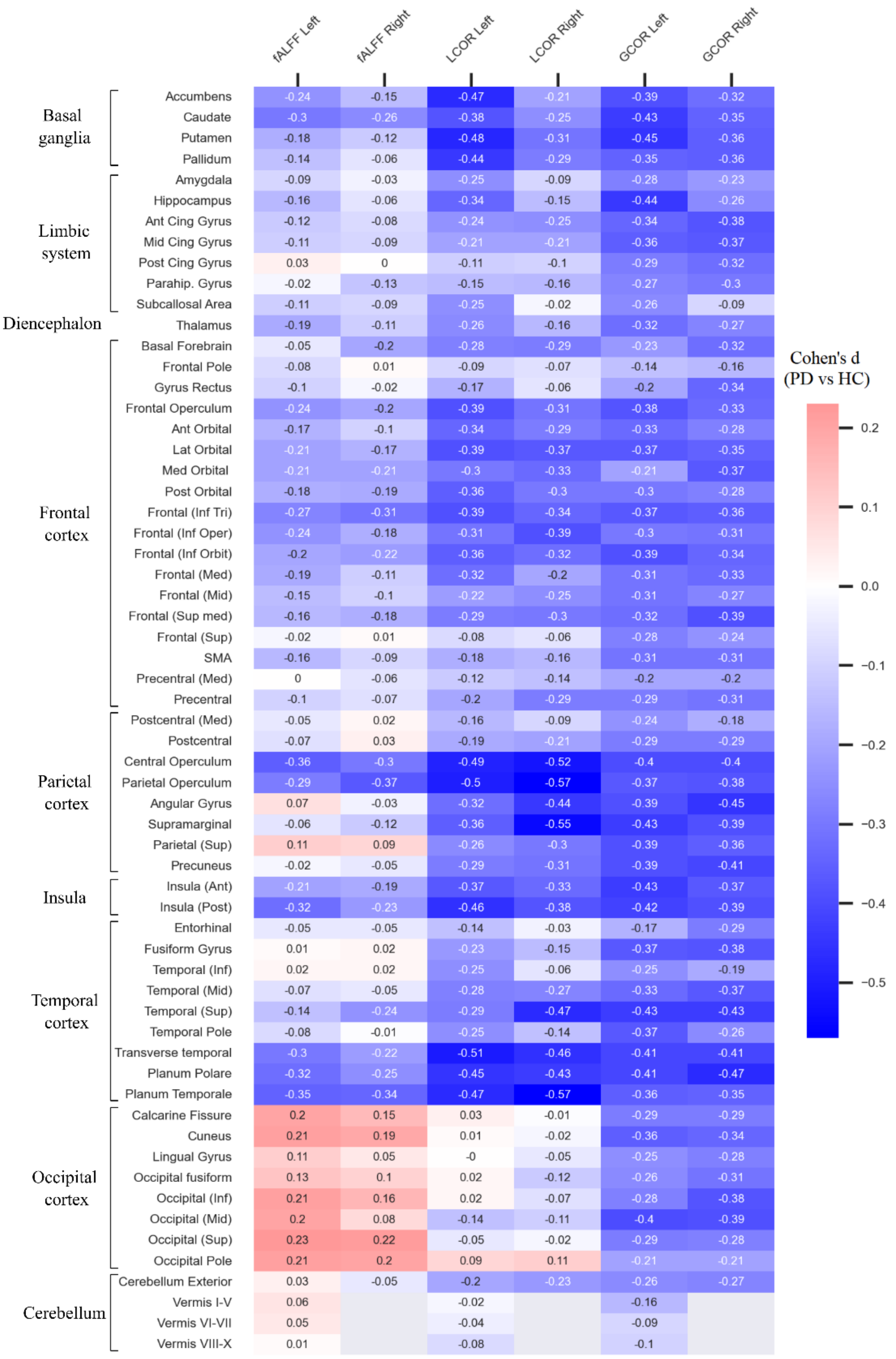


**Supplementary Figure 17:** Regional differences (effect sizes, Cohen's d) in fALFF, LCOR, and GCOR between manifest PD and the age- and sex-matched control group – *before atrophy correction*. Positive values (red cells) indicate higher values of brain functional measures in PD and negative values (blue cells) indicate the inverse. Values of regions of the left or right hemisphere are arranged next to each other (column-wise). The list is sorted from top to the bottom: Basal ganglia, limbic system, diencephalon, frontal cortex, parietal cortex, temporal cortex, occipital cortex, cerebellum. Regional values are visualized in Supplementary Figure 19A.

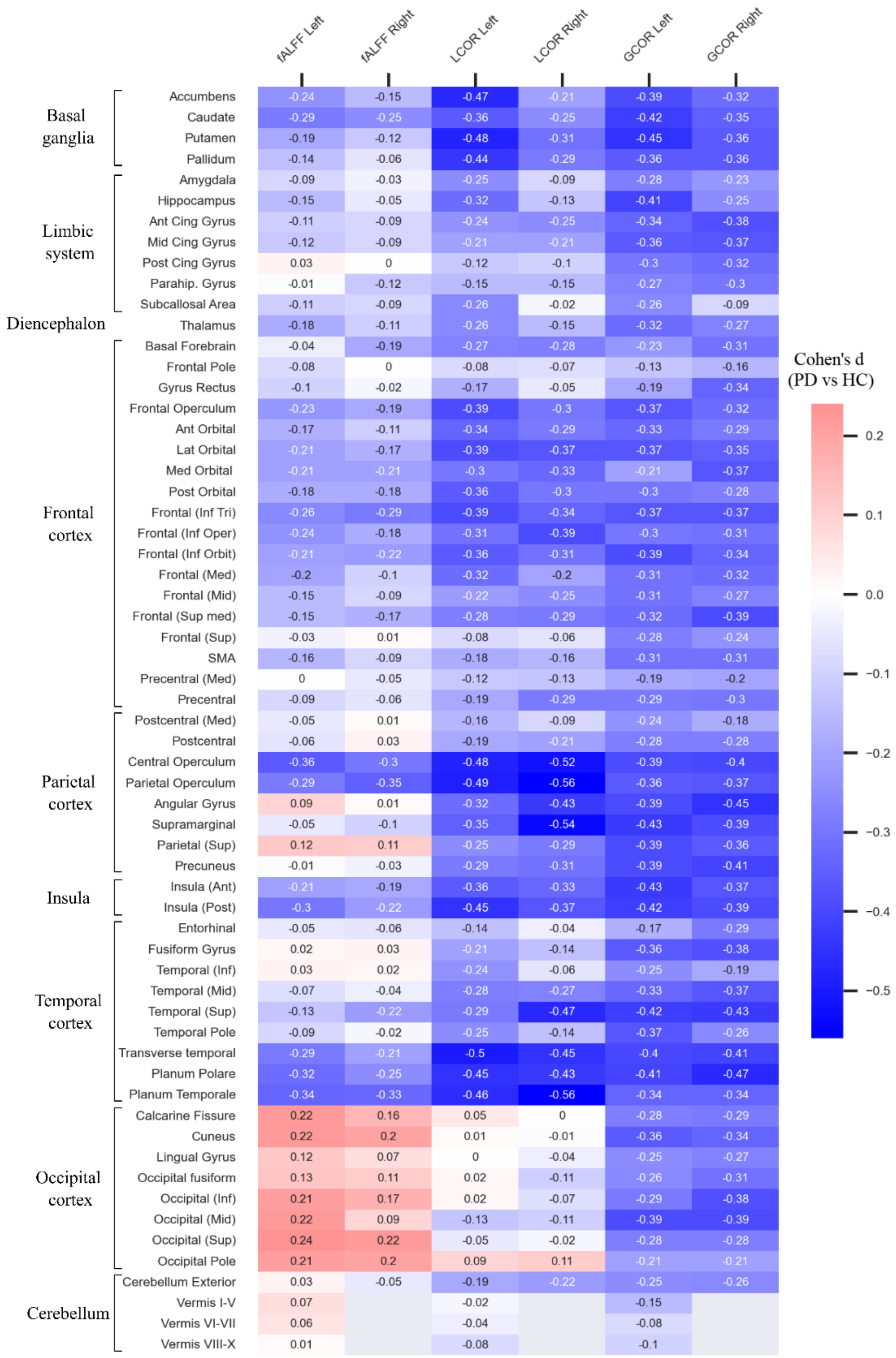


**Supplementary Figure 18:** Regional differences (effect sizes, Cohen's d) in fALFF, LCOR, and GCOR between the age- and sex-matched control group and PD – *after atrophy correction*. Positive values (red cells) indicate higher values of brain functional measures in PD and negative values (blue cells) indicate the inverse. Values of regions of the left or right hemisphere are arranged next to each other (column-wise). The list is sorted from top to the bottom: Basal ganglia, limbic system, diencephalon, frontal cortex, motor area, parietal cortex, temporal cortex, occipital cortex, cerebellum. Regional values are visualized in Supplementary Figure 19B.

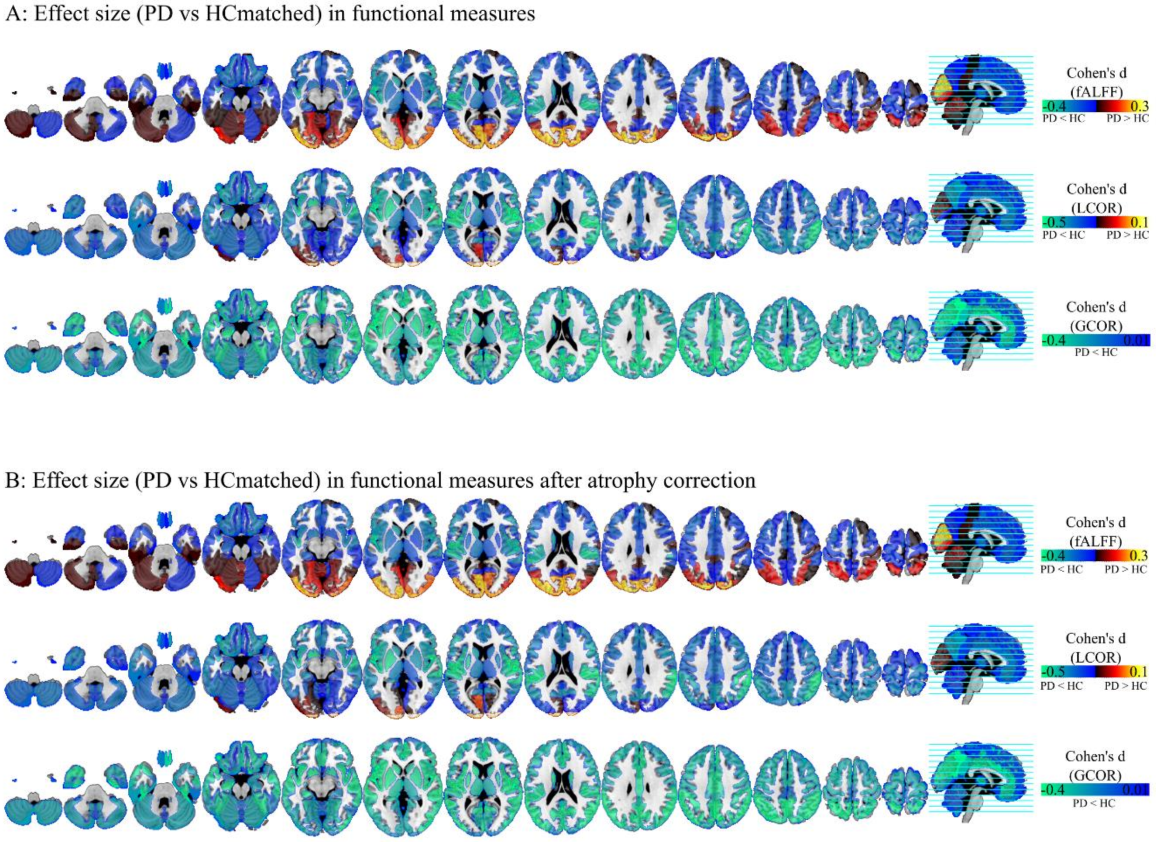


**Supplementary Figure 19**: Maps of regional differences in fALFF, LCOR, and GCOR between the age- and sex-matched control group and PD. Values of regions correspond to those of Supplementary Figure 17 and 18 (atrophy corrected).
